# Gene interaction perturbation network deciphers a high-resolution taxonomy in colorectal cancer

**DOI:** 10.1101/2022.09.02.506442

**Authors:** Zaoqu Liu, Siyuan Weng, Qin Dang, Hui Xu, Yuqing Ren, Chunguang Guo, Zhe Xing, Zhenqiang Sun, Xinwei Han

**Author notes:** **Correspondence to:** Department of Interventional Radiology, The First Affiliated Hospital of Zhengzhou University, Zhengzhou, China. Email Address (Xinwei Han). Department of Colorectal Surgery, The First Affiliated Hospital of Zhengzhou University, Zhengzhou, China. Email Address (Zhenqiang Sun).

## Abstract

Molecular subtypes of colorectal cancer (CRC) are currently identified via the snapshot transcriptional profiles, largely ignoring the dynamic changes of gene expressions. Conversely, biological networks remain relatively stable irrespective of time and condition. Here, we introduce an individual-specific gene interaction perturbation network-based (GIN) approach and identify six GIN subtypes (GINS1-6) with distinguishing features: (i) GINS1 (proliferative, 24%∼34%), elevated proliferative activity, high tumor purity, immune-desert, *PIK3CA* mutations, and immunotherapeutic resistance; (ii) GINS2 (stromal-rich, 14%∼22%), abundant fibroblasts, immune-suppressed, stem-cell-like, *SMAD4* mutations, unfavorable prognosis, high potential of recurrence and metastasis, immunotherapeutic resistance, and sensitive to fluorouracil-based chemotherapy; (iii) GINS3 (*KRAS*-inactivated, 13%∼20%), high tumor purity, immune-desert, activation of *EGFR* and ephrin receptors, chromosomal instability (CIN), fewer *KRAS* mutations, *SMOC1* methylation, immunotherapeutic resistance, and sensitive to cetuximab and bevacizumab; (iv) GINS4 (mixed, 10%∼19%), moderate level of stromal and immune activities, transit-amplifying-like, and *TMEM106A* methylation; (v) GINS5 (immune-activated, 12%∼24%), stronger immune activation, plentiful tumor mutation and neoantigen burden, microsatellite instability and high CpG island methylator phenotype, *BRAF* mutations, favorable prognosis, and sensitive to immunotherapy and *PARP* inhibitors; (vi) GINS6, (metabolic, 5%∼8%), accumulated fatty acids, enterocyte-like, and *BMP* activity. Overall, the novel high-resolution taxonomy derived from an interactome perspective could facilitate more effective management of CRC patients.

## Introduction

Colorectal cancer (CRC) is a worldwide health issue, representing a heterogeneous and aggressive disease with the leading cause of tumor-associated lethality(***Sung et al., 2021***). Currently, pathological staging is broadly but inadequately used to guide clinical management due to diverse clinical outcomes of patients within the same stage(***Liu et al., 2022***). The inherent heterogeneity between patients hampers the individualized treatment of CRC. Development of molecular classification takes the plunge toward more effective interventions and provides critical insights into CRC heterogeneity(***Guinney et al., 2015***; ***Isella et al., 2017***; ***De Sousa et al., 2013***; ***Sadanandam et al., 2013***; ***Marisa et al., 2013***). However, molecular subtypes with distinctive peculiarities and outcomes are mainly identified based on the snapshot transcriptional profiles, largely ignoring the dynamic changes of gene expressions in a biological system(***Guinney et al., 2015***; ***Isella et al., 2017***; ***De Sousa et al., 2013***; ***Sadanandam et al., 2013***; ***Marisa et al., 2013***; ***Chen et al., 2021***)^9^. Indeed, gene expressions are commonly variable at distinct time points or conditions, so that the subtypes based on expression data are unstable and difficult to reproduce(***Chen et al., 2021***). Conversely, biological networks remain relatively stable irrespective of time and condition, and could more reliably characterize the biological state of bulk tissues(***Chen et al., 2021***; ***Sahni et al., 2015***; ***Li et al., 2019***). Previous studies have demonstrated that network analysis is well documented and applied in high-dimensional data, performing more robustly and effectively than single-gene approach(***Chen et al., 2021***; ***Sahni et al., 2015***). Nevertheless, most network-based methods merely focus on gene nodes in the biological network, but ignore the interactions among genes.

To tackle this issue, we introduced a rank-based individual-specific gene interaction perturbation approach(***Chen et al., 2021***), which not only leveraged gene node information but also included vital interaction information in the biological network. Gene interactions are highly conservative in normal samples but broadly perturbed in diseased tissues(***Sahni et al., 2015***). The interaction perturbation within the network can quantify the interaction change for each gene pair. Thus, the overall perturbation of all gene pairs in the background network is reasonably and effectively utilized to characterize the pathological condition at the individual level. Using the individual-specific gene interaction perturbation network-based program, we identified and diversely validated six gene interaction network-based subtypes (GINS1-6) with distinct clinical and molecular peculiarities. Our results provided a high-resolution classification system and improved the understanding of CRC heterogeneity from an interactome perspective.

## Results

### Six CRC subtypes were identified from the gene interaction-perturbation network

To decipher the heterogeneous subtypes from the interaction-perturbation matrix (***Methods, Figure 1***), we selected the representative features that significantly distinguished tumor from normal samples and maintained high variability within all tumor samples for clustering analysis, which formed a network with 1,390 genes and 2,225 interactions (***Supplementary Methods***). This new network also met the scale-free distribution (*R* =-0.994, *P* <2.2e-16; ***Figure 1-figure supplement 1E***) and was visualized in ***Figure 1-figure supplement 1F***.

Consensus clustering analysis(***Wilkerson and Hayes, 2010***) on the discovery cohort with 2,167 CRC samples and 2,225 gene interactions, initially tested potential clustering numbers (*K* =2-10). The cumulative distribution function (CDF) curve and the proportion of ambiguous clustering (PAC) score(***Senbabaoglu et al., 2014***) of the consensus score matrix suggested the optimal *K* =6, which was also achieved from the Nbclust assessment (***Figure 2A and Figure 2-figure supplement 1A-C***). The silhouette statistic was utilized to identify the samples that best represented one of six gene interaction-perturbation network subtypes (GINS), yielding a core set of 1,957 CRC samples (***Figure 2-figure supplement 1D***). The Uniform Manifold Approximation and Projection (UMAP)(***Becht et al., 2018***) cast all samples in two-dimensional spatial coordinates, showing good discrimination (***Figure 2B***).

**Figure 1.**
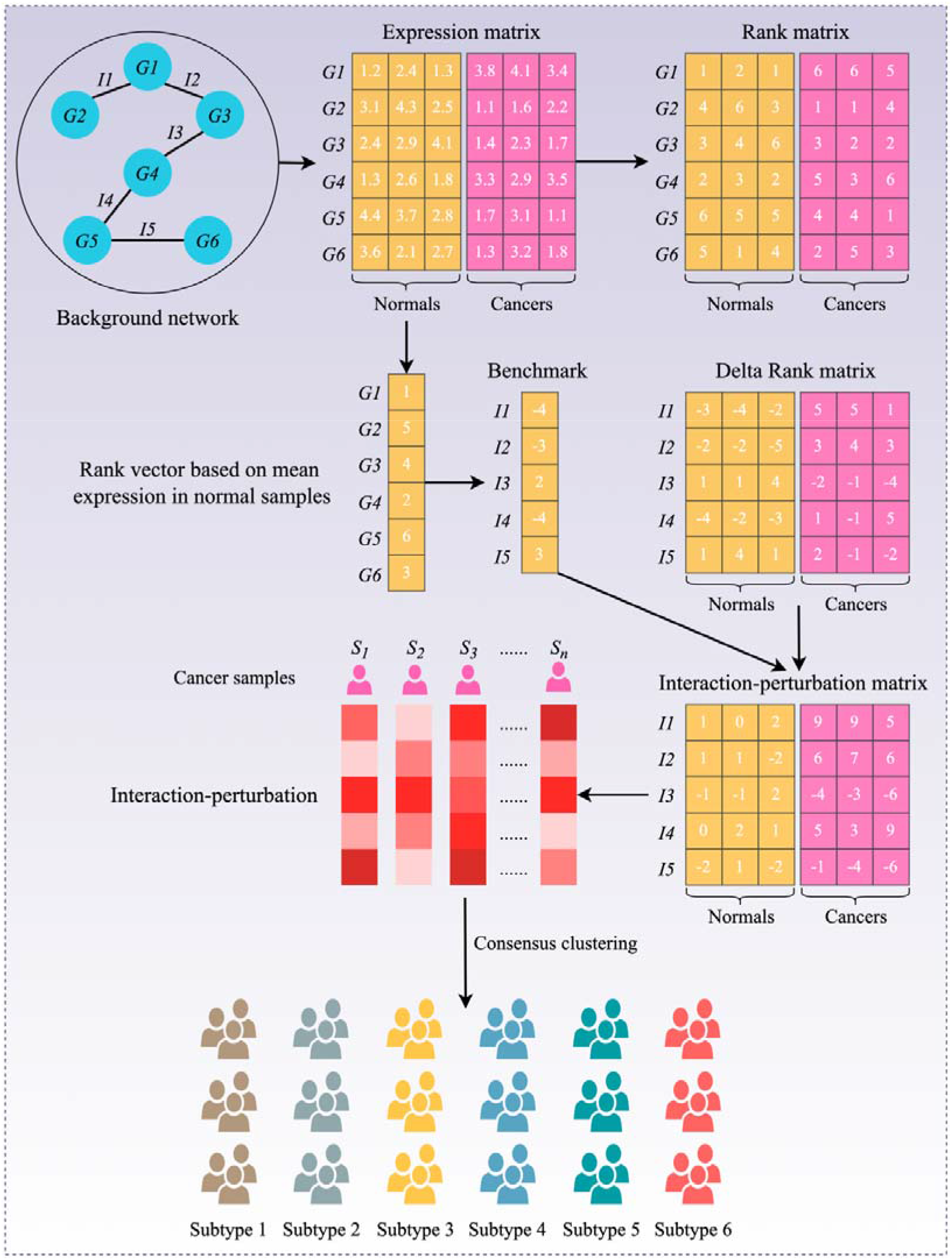
Flowchart of the interaction-perturbation-based program. As an example, the background network consists of six genes and five interactions. There were three normal samples (yellow) and three cancer samples (pink). A rank matrix was obtained by ranking the genes according to the expression value of each sample. The rank matrix was converted to a delta rank matrix with five rows and six columns representing interactions and samples, respectively. The benchmark delta rank vector was calculated as the delta rank of the average expression value in all normal samples. The interaction-perturbation matrix was obtained by subtracting the benchmark delta rank vector from the delta rank matrix.

**Figure 2.**
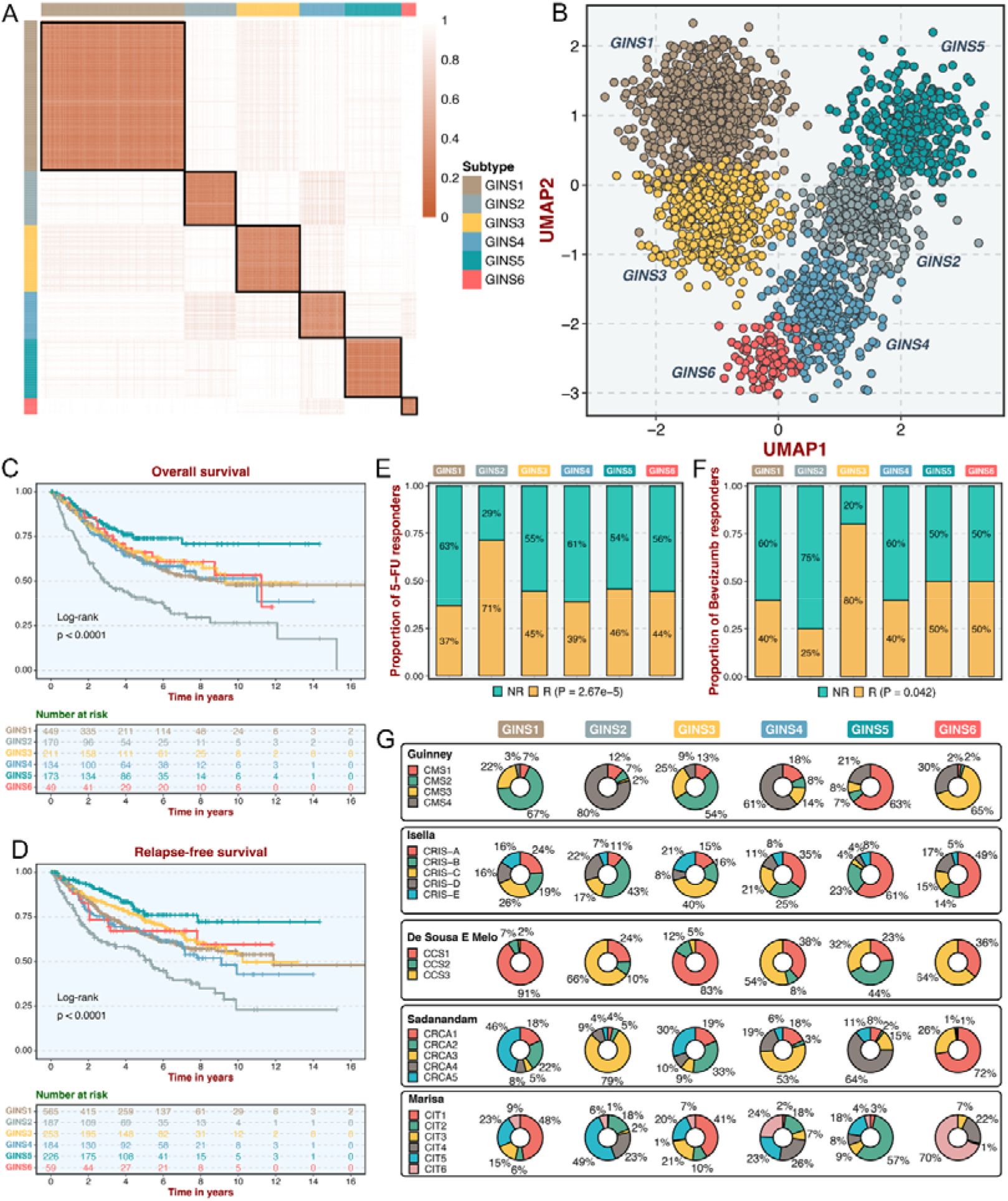
Six CRC subtypes were identified from the gene interaction-perturbation network. **A.** The consensus score matrix of all samples when *K*□achieved 6. A higher consensus score between two samples indicates they are more likely to be grouped into the same cluster in different iterations. **B.** The UMAP analysis cast all samples in two-dimensional spatial coordinates, showing good discrimination. **C-D.** Kaplan-Meier curves of overall survival and relapse-free survival with log-rank test for six GINS subtypes. Log-rank test. **E-F.** Barplots showed the distribution of fluorouracil-based adjuvant chemotherapy (**E**) and bevacizumab (**F**) responders in six subtypes. Fisher’s exact test. **G.** Pie charts showed the proportion of other CRC subtypes in the current GINS taxonomy.

In the six subtypes, age and gender did not differ in distribution (*P* >0.05; ***Figure 2-figure supplement 2A-B***), whereas the clinicopathological stage was more advanced in GINS2 than the other subtypes (*P* <0.05; ***Figure 2-figure supplement 2C-F***). Microsatellite instability (MSI), a well-established biomarker in CRC(***Raskov et al., 2020***), was prominently enriched in GINS5 (*P* <2.2e-16; ***Figure 2-figure supplement 2G***). Kaplan-Meier survival analysis demonstrated significant survival differences among six subtypes. GINS2 had the worst prognostic outcomes of overall survival (OS) and relapse-free survival (RFS), whereas GINS5 portended the most favorable prognosis, and the other four subtypes displayed intermedium OS and RFS (OS, *P* <0.0001; RFS, *P* <0.0001; ***Figure 2C-D***). Additionally, GINS2 benefited more from fluorouracil-based adjuvant chemotherapy (ACT) in the discovery cohort with 79 responders and 187 non-responders (*P* =2.67e-5, ***Figure 2E***). We further explored the association of GINS subtypes with ACT after surgery for 585 patients in one subseries of the discovery cohort, GSE39582, which stored complete ACT information. The six subtypes presented concordant distribution in survival across all samples (*P* =0.0021, ***Figure 2-figure supplement 3A***). Subsequent analysis focused on each subtype and revealed that only GINS2 tumors had significantly improved survival after ACT treatment (***Figure 2-figure supplement 3B-G***), suggesting these patients were preferentially responsive to ACT. Conversely, GINS2 might benefit less from bevacizumab in the discovery cohort (25 responders and 29 non-responders), whereas GINS3 possessed a large proportion of responders (*P* =0.042; ***Figure 2F***).

To compare our subtypes with previously reported CRC classifications, the discovery cohort was reclassified according to the previous subtype criteria, including consensus molecular subtypes (CMS)(***Guinney et al., 2015***), CRC intrinsic subtypes (CRIS)(***Isella et al., 2017***), colon cancer subtypes (CCS)(***De Sousa et al., 2013***), CRCAssigner (CRCA)(***Sadanandam et al., 2013***), and Cartes d’Identit é des Tumeurs(***Marisa et al., 2013***), respectively. Noteworthy connections were observed between our subtypes and these previous classifications, indicating a biological convergence (***Figure 2G*)**. Specifically, GINS1 was related to the canonical CMS2, CCS1, CRCA5, and CIT1; GINS2 was associated with the more aggressive subtypes, including CMS4, CRIS-B, CCS3, CRCA3, and CIT5; GINS3 was linked to CMS2, CRIS-C, CCS1, CRCA2/5, and CIT1; GINS4 was correlated with CMS4, CRIS-A, CCS1/3, and CRCA3; GINS5 was predominantly enriched in MSI-like subtypes, containing CMS1, CRIS-A, CCS2, CRCA4, and CIT2; GINS6 was associated with CMS3, CRIS-A, CCS3, CRCA1, and CIT6. Overall, the aggressiveness properties shown in other classifications were consistent with our six subtypes. Notably, only approximately 50% of our classifier genes overlapped with the signature genes of all previous CRC classifications (***Figure 2-figure supplement 4***), suggesting a significant molecular convergence, but also leaving a rich exploration space for our classification.

### Six subtypes were reproductive and stable in 19 independent datasets

To identify GINS subtypes in novel datasets using a small list of genes, a gene centroid classifier was developed (***Supplementary Methods***). We first identified genes correlated with the six subtypes using significance analysis of microarrays(***Tusher et al., 2001***), followed by prediction analysis for microarrays(***Tibshirani et al., 2002***) to determine 289 subtype-discriminant genes with the lowest misclassification error (1.8%) (***Supplementary File 1***). Subsequently, a 289-gene centroid-based classifier based on the diagonal quadratic discriminant analysis (DQDA) rule(***Marisa et al., 2013***) was developed, and validation datasets were independently assigned to six subtypes. The validation works focused on the following four contexts: (1) data from the same platform (GPL570); (2) data from different platforms and sequencing techniques (microarray or RNA-seq); (3) microdissected or whole tumors; (4) in-house clinical setting.

Initially, significant subtype assignments were performed on seven datasets from the same platform via the 289-gene centroid-based classifier (***Supplementary Methods***). Six subtypes were confidently identified, and Subclass Mapping (SubMap) analysis(***Hoshida et al., 2007***) confirmed that each subtype was associated with similar underlying transcriptional traits in the discovery cohort (***Figure 3-figure supplement 1***). The same results were achieved on seven microarrays from different platforms and one RNA-seq dataset (TCGA-CRC Illumina; ***Figure 3-figure supplement 2***). Two datasets, GSE26682 and GSE24551, each chip from two different platforms (GPL570 & GPL96 for GSE26682 and GPL5175 & GPL11028 for GSE24551), also displayed superimposable classification patterns sustained by similar transcriptional traits (***Figure 3-figure supplement 3***). Proverbially, spatial genetic and phenotypic diversity within solid tumors has been well documented, which is also dubbed as intra-tumor heterogeneity(***Li et al., 2022***). To address this issue, our analysis using additional datasets (GSE12945 and GSE21510) containing samples from both microdissected and whole tumors, and from tumor RNAs profiled on different microarray platforms, consistently reproduced six subtypes with particular molecular traits (***Figure 3A-B***). This is similar to what has been suggested in breast cancer, where subtypes are routinely identified despite possible intra-tumoral heterogeneity(***Polyak, 2011***). In the discovery cohort and 19 validation datasets, we found comparable fractions of patients being assigned to each subtype (***Figure 3C***), which demonstrated that our classification was stable and universal within different datasets. In addition to the identified and attributed subtypes sharing similar transcriptional traits, clinical features were also characterized in validation datasets. Likewise, GINS2 possessed more advanced tumors (***Supplementary File 2***), preferentially metastasized (***Figure 3-figure supplement 4A-F***), and behaved adverse OS (***Figure 3D-H***) and RFS (***Figure 3I and Figure 3-figure supplement 4G-H***). The MSI tumors were prone to occur in GINS5 (***Figure 3J-P***) with the most favorable OS (***Figure 3D-H***) and RFS (***Figure 3I and Figure 3-figure supplement 4G-H***). ACT treatment also exhibited the identical response distribution, with GINS2 achieving more clinical benefit (***Figure 3Q***). Cetuximab with function to target *EGFR*(***Raskov et al., 2020***), performed better in GINS3 (***Figure 3R***). Overall, six subtypes not only maintained comparable proportions, but also shared analogical transcriptional and clinical traits in the discovery cohort and 19 validation datasets.

**Figure 3.**
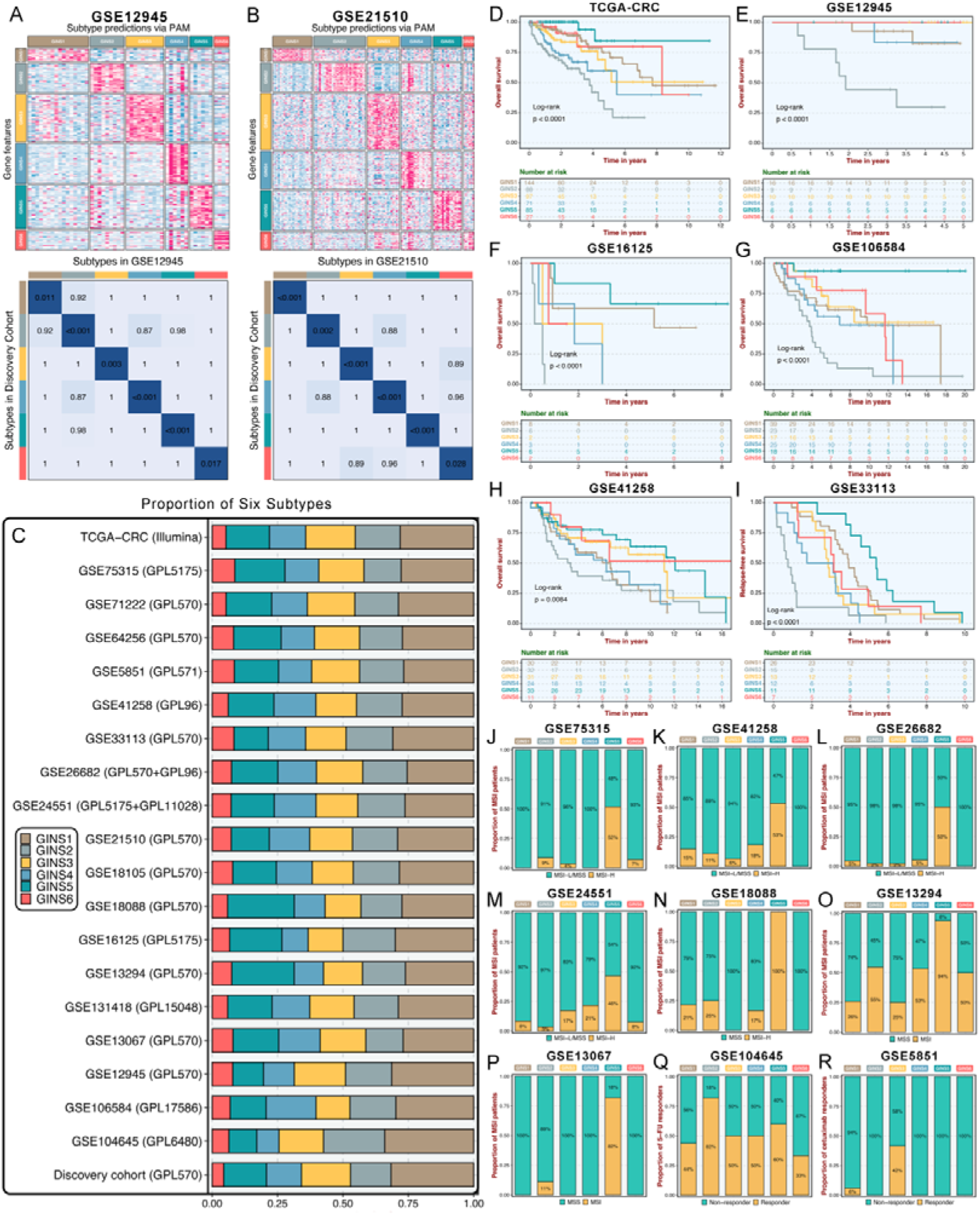
Six subtypes were reproductive and stable in 19 independent datasets. **A-B.** The GSE12945 (**A**) and GSE21510 (**B**) were assigned in six subtypes according to the classifier. The top and left bars indicated the subtypes. In the heatmap, rows indicated genes from the classifier and columns represent patients. The heatmap was color-coded on the basis of median-centered log_2_ gene expression levels (red, high expression; blue, low expression). SubMap plots, located in the bottom panel, assessed expressive similarity between corresponding subtypes from two different cohorts. **C.** Barplots showed comparable fractions of patients being assigned to each subtype in the discovery cohort and 19 validation datasets. **D-H.** Kaplan-Meier curves of overall survival for six GINS subtypes in TCGA-CRC (**D**), GSE12945 (**E**), GSE16125 (**F**), GSE106584 (**G**), and GSE41258 (**H**). Log-rank test. **I.** Kaplan-Meier curves of relapse-free survival for six GINS subtypes in GSE33113. Log-rank test. **J-P.** Barplots showed the distribution of MSI patients across six subtypes in GSE75315 (**J**), GSE41258 (**K**), GSE26682 (**L**), GSE24551 (**M**), GSE18088 (**N**), GSE13294 (**O**), and GSE13067 (**P**). Fisher’s exact test. **Q-R.** Barplots showed the distribution of responders to six subtypes of fluorouracil-based adjuvant chemotherapy in GSE104645 (**Q**) and cetuximab in GSE5851 (**R**). Fisher’s exact test.

### Subtype validation in an in-house clinical cohort

As an initial attempt to facilitate the GINS taxonomy into a clinically translatable tool amenable to clinical applications, we developed a quantitative PCR (qPCR) miniclassifier and further validated our subtypes in 214 clinical CRC samples from our hospital (***Supplementary File 3***). Using 289 genes from the PAM classifier, we firstly identified 93 subtype-specific robust genes via paired differential expression analysis (all *P* <0.01) and bootstrap logistic regression (1000 iterations and all *P* <0.05) (***Figure 4A***). Subsequently, the LASSO framework based on 10-fold cross-validation and one-standard-error rule determined the 14 most informative genes that integratively fitted a random forest model (***Figure 4A and Supplementary File 4***). Initial model development was conducted in the training dataset (70% of the discovery cohort) and then validated in the testing dataset (30% of the discovery cohort). Confusion matrix displayed the general tendency of classification effect, with a misclassification error of 7.8% and 13.0% in the training and testing datasets, respectively (***Figure 4-figure supplement 1A-B***). The accuracy, precision, recall, F1-score, and specificity of the random forest model reached a quite respectable level, suggesting this miniclassifier comprised of 14 key genes was robust to assign six subtypes in a new cohort (***Figure 4B***).

**Figure 4.**
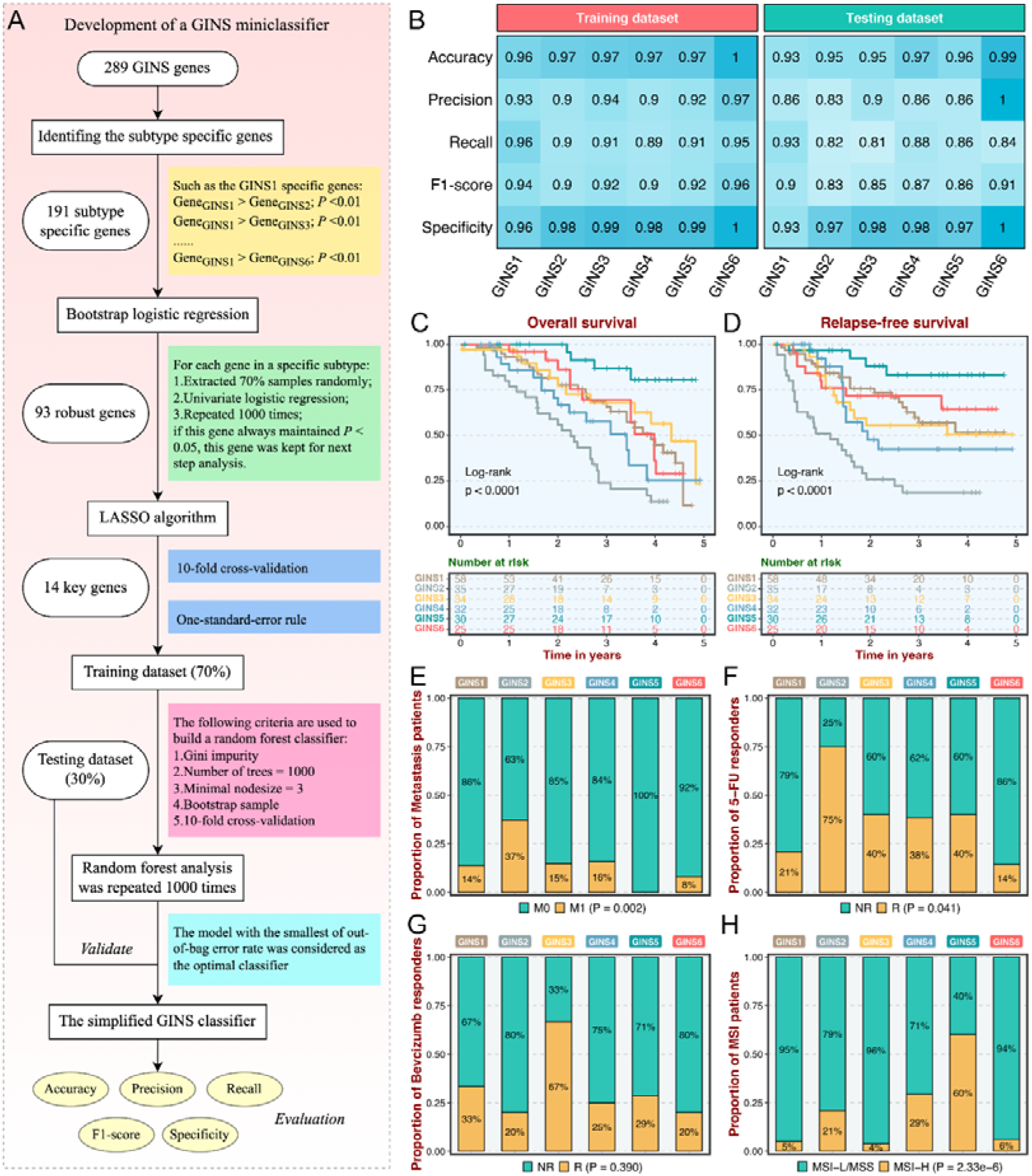
Subtype validation in an in-house clinical cohort. **A.** Overview of the miniclassifier development procedures. **B.** Performance of the miniclassifier in the training and testing datasets. **C-D.** Kaplan-Meier curves of overall survival and relapse-free survival with log-rank test for six GINS subtypes. Log-rank test. **E-H.** Barplots showed the distribution of metastasis patients (**E**), fluorouracil-based adjuvant chemotherapy responders (**F**), bevacizumab responders (**G**), and MSI patients (**H**) in six subtypes. Fisher’s exact test.

To test the clinical interpretation of this miniclassifier, another validation based on qPCR results from 214 frozen CRC tissues was deployed to verify its feasibility in clinical settings. With the expression profiles of 14 key genes in each patient, the miniclassifier successfully isolated six subtypes (***Supplementary File 5***). In line with our prior findings, GINS2 had shorter OS and RFS (*P* <0.0001, ***Figure 4C-D***), behaved a stronger propensity to invade and metastasize (*P* =0.002, ***Figure 4E and Supplementary File 5***), but was sensitive to ACT treatment (*P* =0.041, ***Figure 4F***). Bevacizumab responders were predominantly concentrated in GINS3, whereas GINS2 still failed to achieve clinical efficacy (***Figure 4G***), although not statistically significant (*P* =0.390) due to the small sample size (n =42). A subset of CRC, GINS5 (14%), displayed prolonged prognosis (*P* <0.0001, ***Figure 4C-D***) and enriched more MSI tumors (***Figure 4H***). Hence, the 14-gene miniclassifier could afford the stability and interpretation in clinical practice.

### Biological peculiarities of six subtypes

To better delineate the biological attributes inherent to GINS subtypes, we leveraged the ‘Hallmark’ genesets (***Supplementary File 6***), a comprehensive picture of biological features representing essential oncogenic pathways in cancers(***Hanahan, 2022***). For each sample and signature, an integrated score was computed by subtracting the average expression of genes negatively correlated with the subtype from the average expression of genes positively correlated with the subtype. To assess the extent to which six subtypes captured samples with stronger transcriptional signatures, we introduced a framework termed ‘Sample Set Enrichment Analysis’ (SSEA)(***Isella et al., 2017***). In SSEA, all samples are ranked by the integrated scores, and the ranked sample list is further subjected to the gene set enrichment analysis (GSEA) procedure to test whether the ‘sample set’ for each GINS subtype enriches high-ranking samples. Subsequently, another unsupervised algorithm, gene set variation analysis (GSVA)(***Hanzelmann et al., 2013***), estimated differences in pathway activity across six subtypes.

According to the SSEA and GSVA phenotypic analysis, GINS1 was distinguished by up-regulated cell cycle pathways, suggesting proliferative characteristics for these tumors (***Figure 5A-B, Figure 5-figure supplement 1 and Supplementary File 7***). We next proved that GINS1 also strikingly overexpressed *MKI67* and *PCNA* (*P* <2.2e-16, ***Figure 5-figure supplement 2A-B***), which were identified as important cell cycle-specific antigens in tumors. GINS3 exhibited an inferior level of *KRAS* signaling that was mainly driven by *KRAS* mutations (***Figure 5A-B and Supplementary File 7***)(***Raskov et al., 2020***). Activation of metabolisms (mainly lipid metabolisms) was featured by GINS6, suggesting canonical metabolic reprogramming across these tumors (***Figure 5A-B, Figure 5-figure supplement 1 and Supplementary File 7***). Intriguingly, interactive stromal and immune activation trends shifted in GINS2/4/5 (***Figure 5A-B, Figure 5-figure supplement 1 and Supplementary File 7***). GINS2 was endowed with higher stromal activity and lower immune activity, whereas GINS5 conveyed the opposite trend entirely, concordant with the tumor invasiveness and prognosis of two subtypes, and GINS4 was characterized by a mixed phenotype that displayed moderate level of stromal and immune pathways. *ESTIMATE*(***Yoshihara et al., 2013***), a tool that uses gene expression profiles to infer immune and stromal constituents within the tumor microenvironment (TME), further validated these phenomena (*P* <2.2e-16, ***Figure 5-figure supplement 2C-D***). As three subtypes with abundant TME components, GINS2/4/5 may mutually evolve in stromal and immune functions. Thus, we intended to extract consistently upregulated and downregulated genes among these three subtypes, using *Mfuzz* package, a noise-robust soft clustering analysis with the fuzzy c-means form(Kumar and M, 2007). The *Mfuzz* analysis revealed 10 gene clusters, and gene cluster 3 and 10 displayed the stable expression pattern from GINS2 to GINS5 (***Figure 5C and Supplementary File 8***). As expected, gene cluster 3 was prevailingly associated with immune infiltration and activation (***Figure 5D***), whereas gene cluster 10 was prominently characterized by stromal activation and remodeling (***Figure 5E***), which further supported our findings. This also indicated that TME had profound impacts on the progression and prognosis of tumors, and GINS2/5 acted as two extremes of TME components, indeed showing diametrically opposite clinical outcomes. Of note, GINS1/3 displayed scarce stromal and immune components (***Figure 5A-B, Figure 5-figure supplement 1, and Figure 5-figure supplement 2C-D***), instead, tumors within these subtypes possessed higher purity (***Figure 5-figure supplement 2E***).

**Figure 5.**
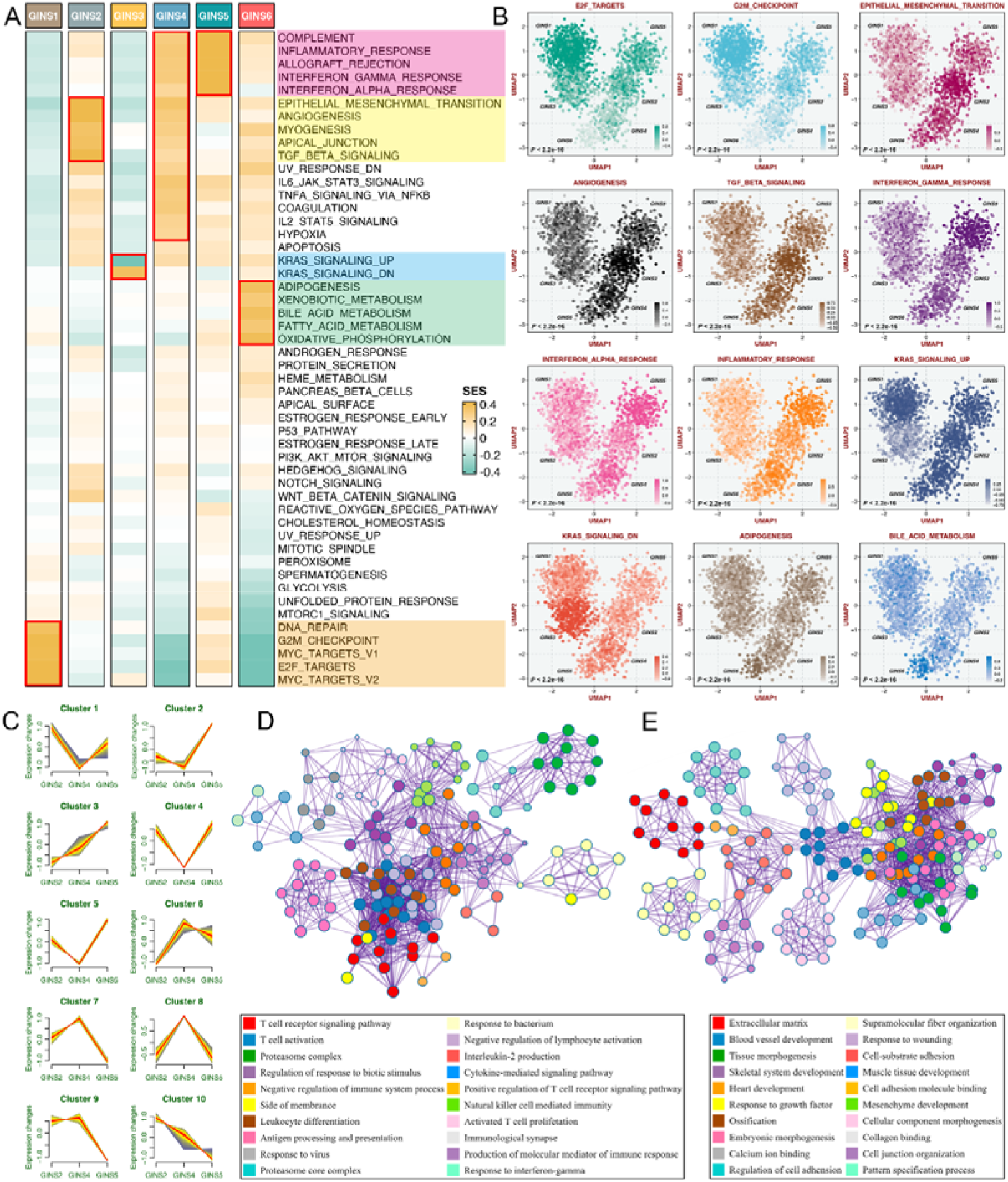
Biological peculiarities of six subtypes. **A.** SSEA-based analysis delineated the biological attributes inherent to GINS subtypes. **B.** GSVA further estimated differences in pathway activity across six subtypes. Anova test. **C.** Ten gene clusters were obtained via the soft clustering method (*Mfuzz*) in GINS2/4/5. **D-E**. Enrichment analysis of gene cluster 3 (**D**) and 10 (**E**).

### Immune landscape and immunotherapeutic potential of six subtypes

To further investigate the immune regulations of GINS subfamilies, we profiled five classes of immunomodulators (145 molecules in total), including antigen presentation molecules, immunoinhibitors, immunostimulators, chemokines, and receptors. These immunomodulators are crucial for cancer immunotherapy with specific agonists and antagonists in clinical oncology(***Tang et al., 2018; Thorsson et al., 2018***). Our results delineated that transcriptional expression of immunomodulators varied across GINS subtypes, and tumors with high expression pattern of immunomodulators were predominantly assigned to GINS5 (***Figure 6A and Supplementary File 9***). To better illustrate this at protein level, we took advantage of the proteome (Reverse Phase Protein Array) data available from the TCGA portal(***Cancer Genome Atlas, 2012***), but with only 26 immunomodulators (***Supplementary File 10***). Using PAM-centroid distance classifier, all samples were attributed to corresponding subtypes. Differential analysis with the thresholds of Benjamini-Hochberg false discovery rate <0.05 and log_2_ (fold change) >1 was performed between GINS5 and other subtypes, and we observed 13/26 of immunomodulators were up-regulated in GINS5 (***Figure 6B***). More specifically, 12/13 of significant immunomodulators are involved in antigen presentation, another protein was *IDO1*, an emerging immune checkpoint that overexpresses in multiple cancers(***Zhai et al., 2018***). GINS5 was also characterized by a stronger immunogenicity that harbored remarkably higher tumor mutation burden (TMB) and neoantigen load (NAL) (P <0.001, ***Figure 6C***), possibly further inducing abundant immune elements and regulations.

**Figure 6.**
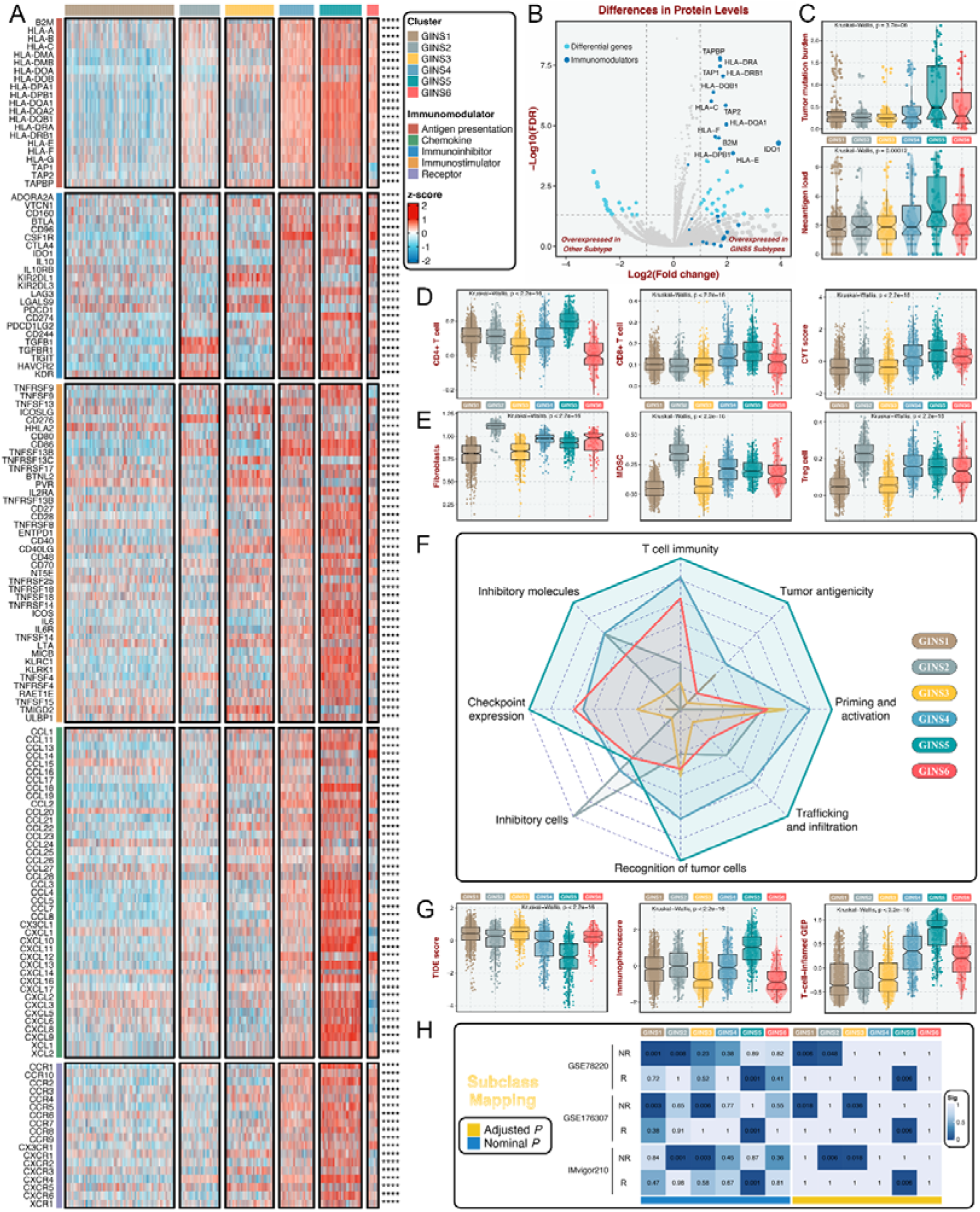
Immune landscape and immunotherapeutic potential of six subtypes. **A.** The expression distribution of 145 immunomodulators among six subtypes. \*\*\*\**P* <0.0001. **B.** Differential analysis was performed between GINS5 and other subtypes, and 13/26 of immunomodulators were up-regulated in GINS5. **C.** The distribution of TMB and NAL score among six subtypes. Kruskal-Wallis test. **D.** The distribution of CD4+ T cells, CD8+ T cells, and CYT score among six subtypes. Kruskal-Wallis test. **E.** The distribution of fibroblasts, MDSC, and Treg cells among six subtypes. Kruskal-Wallis test. **F.** Radar plots showed the immunogram patterns of the six subtypes. Kruskal-Wallis test. **G.** The distribution of TIDE score, immunophenoscore, and T-cell-inflamed gene expression profiles (GEP) among six subtypes. Kruskal-Wallis test. **H.** SubMap analysis delineated the similar expression pattern between GISN5 tumors and immunotherapeutic responders from three cohorts with treatment annotations, and GINS1/2/3 shared the transcriptional modes with non-responders from 2/3 of immunotherapeutic cohorts. ‘R’ represents responder, whereas ‘NR’ represents non-responder.

Previous reports introduced several bioinformatics tools based on gene expression profiles to quantify the infiltration and activation of immune cells in solid tumors(***Charoentong et al., 2017***; ***Newman et al., 2019***; ***Becht et al., 2016***; ***Rooney et al., 2015***). Using these tools, we found that rich infiltration and strong immune killing of T cells were particularly evident in GINS5, coincident with the abovementioned findings (***Figure 6D***). Moreover, GINS5 also possessed the abundant infiltration of Th1, Th2, and M1 macrophages(***Mills et al., 2016***) (***Figure 6-figure supplement 1A-C***), which could secrete proinflammatory cytokines and enhance immune activation. Conversely, M2 traditionally regarded as promoting tumor growth by suppressing cell-mediated immunity and subsequent cancer cell killing(***Mills et al., 2016***), was significantly elevated in GINS2 (***Figure 6-figure supplement 1D***). In line with this, three other classical immunosuppressive cells, including fibroblasts, myeloid-derived suppressor cells (MDSC), and Treg cells(***Hicks et al., 2022***), were also significantly enriched in GINS2 (***Figure 6E***). Apart from the immune activation represented by GINS5, GINS1/2/3 displayed sparse infiltration of cells that promote immune activity (***Figure 6D and Figure 6-figure supplement 1A-C***), but unlike GINS2, GINS1/3 were also characterized by rare immunosuppressive cells (***Figure 6E and Figure 6-figure supplement 1D***), consistent with their high tumor purity. GINS4/6 subfamilies were featured as the mixed phenotypes with immune activating and inhibitory components (***Figure 6D-E and Figure 6-figure supplement 1A-D***).

To systematically evaluate immunotherapeutic potential of six subtypes, we built an immunogram for the cancer-immunity cycle (CIC) (***Figure 6F***), which was based on the rationale that immunity within tumors is a dynamic process and proposed by Karasaki and colleagues(***Karasaki et al., 2017***). Together, we annotated six subtypes by specific immune features: (i) GINS1/3, thereafter designated the ‘immune-desert’ phenotype, was endowed with scarce immune fractions; (ii) GINS2, thereafter designated the ‘immune-suppressed’ phenotype, was enriched for abundant inhibitory cells; (iii) GINS5, thereafter designated the ‘immune-activated’ phenotype, was dramatically linked to superior tumor immunogenicity and extensive immune activation; and (iv) GINS4/6, thereafter designated the ‘mixed’ phenotype, was characterized by moderate levels of immunity cycle score (***Figure 6F***).

Among these six subtypes, patients with lower tumor immune dysfunction and exclusion score(***Jiang et al., 2018***), higher immunophenoscore(***Charoentong et al., 2017***) and T-cell-inflamed gene expression profiles(***Ott et al., 2019***), were proven to favor benefit from immunotherapy, and predominantly assigned to GINS5 (***Figure 6G***). SubMap analysis(***Hoshida et al., 2007***) also delineated the similar expression pattern between GISN5 tumors and immunotherapeutic responders from three cohorts with treatment annotations, and GINS1/2/3 shared the transcriptional modes with non-responders from 2/3 of immunotherapeutic cohorts (***Figure 6H***). Collectively, GINS5 tumors might generate clinical benefit from immunotherapy, whereas GINS1/2/3 were not suitable for this treatment due to potential immune-related adverse events and high cost.

### GINS6 tumors conveyed rich lipid metabolisms

Prior results indicated that GINS6 was characterized by activation of metabolism pathways. To investigate an extensive spectrum of metabolic reprogramming in GINS6, we executed GSEA against 69 metabolic pathways from the Kyoto Encyclopedia of Genes and Genomes (KEGG) database(***Chen et al., 2021***) (***Supplementary File 11***). In total, 20 pathways were significantly enriched in GINS6 versus other subtypes, and most pathways were upregulated (***Figure 7A and Supplementary File 11***). Notably, GSEA demonstrated that lipid metabolisms were the most significant metabolic processes in GINS6 (***Figure 7A-B and Supplementary File 11***). Using principal component analysis (PCA), we found that only the lipid metabolism profiles could distinguish GINS6 from other subtypes in spatial distribution (***Figure 7C and Figure 7-figure supplement 1***). The SSEA-based framework further confirmed that GINS6 predominantly enriched high-ranking samples with stronger lipid signature scores (***Figure 7D***). Subsequently, we established a metabolite-protein interaction network (MPIN)(***Chen et al., 2021***) via nine GINS6-specific genes with broad and tight connections with lipid metabolites (***Figure 7E and Supplementary File 12***). Indeed, 7/9 of these genes belonged to lipid metabolic pathways. To explore the metabolic profiles from the perspective of metabolomics, we enrolled 55 CRC cell lines with both transcriptome and metabolomics data (including 225 metabolites) from Cancer Cell Line Encyclopedia (CCLE)(***Li et al., 2019***). All cell lines were assigned to corresponding subtypes via our PAM-centroid distance classifier. We compared the metabolite abundances between GINS6 and other subtypes, and found that GINS6 exhibited higher levels in four fatty acids including α-glycerophosphate, adipate, taurocholate, and aconitate. Additionally, four carnitines containing stearoylcarnitine, myristoylcarnitine, valerylcarnitine, and malonylcarnitine, that serve as vital compounds in lipid metabolism processes, were also dramatically accumulated in GINS6 (***Figure 7F***). These findings validated that GINS6 was closely associated with metabolic reprogramming and accumulated fatty acids, suggesting GINS6 tumors might be more sensitive to metabolic inhibitors targeting fatty acid metabolisms.

**Figure 7.**
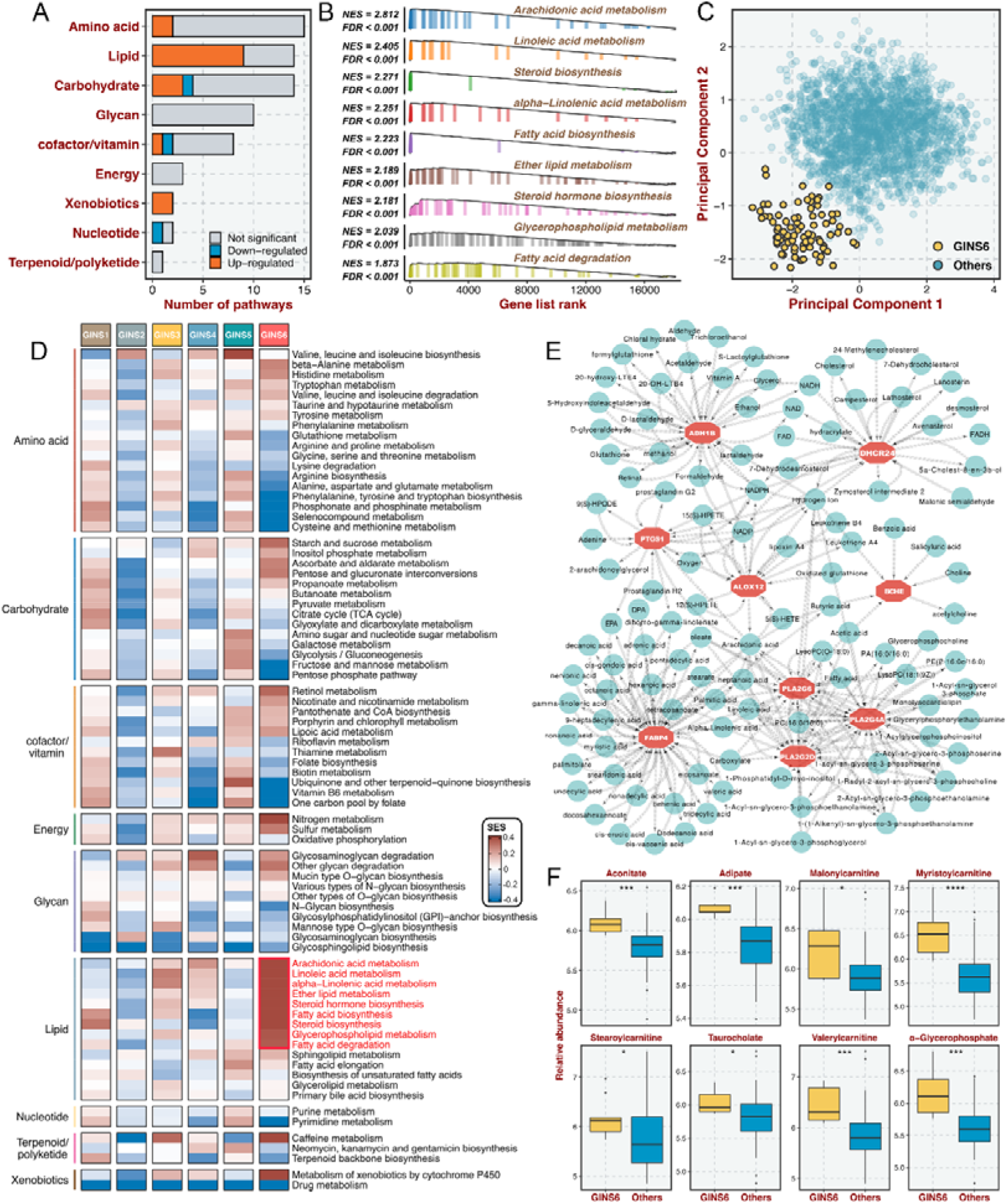
GINS6 tumors conveyed rich lipid metabolisms. **A.** The number of metabolic pathways that was significantly upregulated or downregulated (FDR < 0.05) in GINS6 versus the other subtypes among each of nine metabolic categories. **B.** GSEA plots of nine lipid metabolism pathways (FDR < 0.001). **C.** Principal component analysis of all samples in the discovery cohort for the mRNA expression of lipid metabolic genes. **D.** SSEA-based framework further confirmed that GINS6 predominantly enriched high-ranking samples with stronger lipid signature scores. **E.** Metabolite-protein interaction network (MPIN) was established via nine GINS6-specific genes with broad and tight connections with lipid metabolites. **F.** Metabolomics results further demonstrated that GINS6 featured by abundant fatty acids metabolites. Wilcoxon test. \**P* <0.05, \*\*\**P* <0.001, \*\*\*\**P* <0.0001.

### GINS subtypes were associated with cellular phenotypes and autocrine loops

Using previously supervised signatures derived from cellular phenotypes(***Sadanandam et al., 2013***; ***Marisa et al., 2013***; ***Kosinski et al., 2007***), we identified phenotype origins peculiar to individual GINS classes. In this study, GINS2 appeared highly enriched for stem-cell-like tumors (91%), whereas GINS4 was endowed with transit-amplifying-like phenotype (86%) (***Figure 8A***). GINS5 was characterized by inflammatory (***Figure 8A***), coincident with its biological and immune features. GINS6 featured an enterocyte-like phenotype (***Figure 8A***). Specifically, serrated-like CRC arising from serrated neoplasia pathway(***Marisa et al., 2013***), were predominantly assigned to GINS5 and to a lesser extent to other subtypes (***Figure 8A***). Conversely, conventional-like tumors were mainly shared by non-GINS5 subtypes. From the unsupervised perspective, GSVA further verified the cellular phenotypic differences across six subtypes (***Figure 8B-H***). As previously reported(***Sadanandam et al., 2013***), we also delineated that these cellular phenotypes were linked to distinctive WNT signaling activity and anatomical regions of the colon crypts (***Figure 8I-J***). Moreover, the nearest template prediction (NTP) algorithm(***Hoshida, 2010***) based on published signatures(***Kosinski et al., 2007***) assigned each sample into the crypt base and top phenotypes in the discovery cohort (***Figure 8K***). Consistently, tumors with the crypt base phenotype were particularly evident for GINS2, whereas other subtypes were mainly concentrated on tumors with the crypt top phenotype, especially GINS6. We next curated 26 published stemness signatures from the StemChecker webserver(***Pinto et al., 2015***) and further employed GSVA to quantify the signature score of each pathway. Overall, GINS2 displayed superior abundance relative to other subtypes, which was in line with its malignant traits (***Figure 8L***).

**Figure 8.**
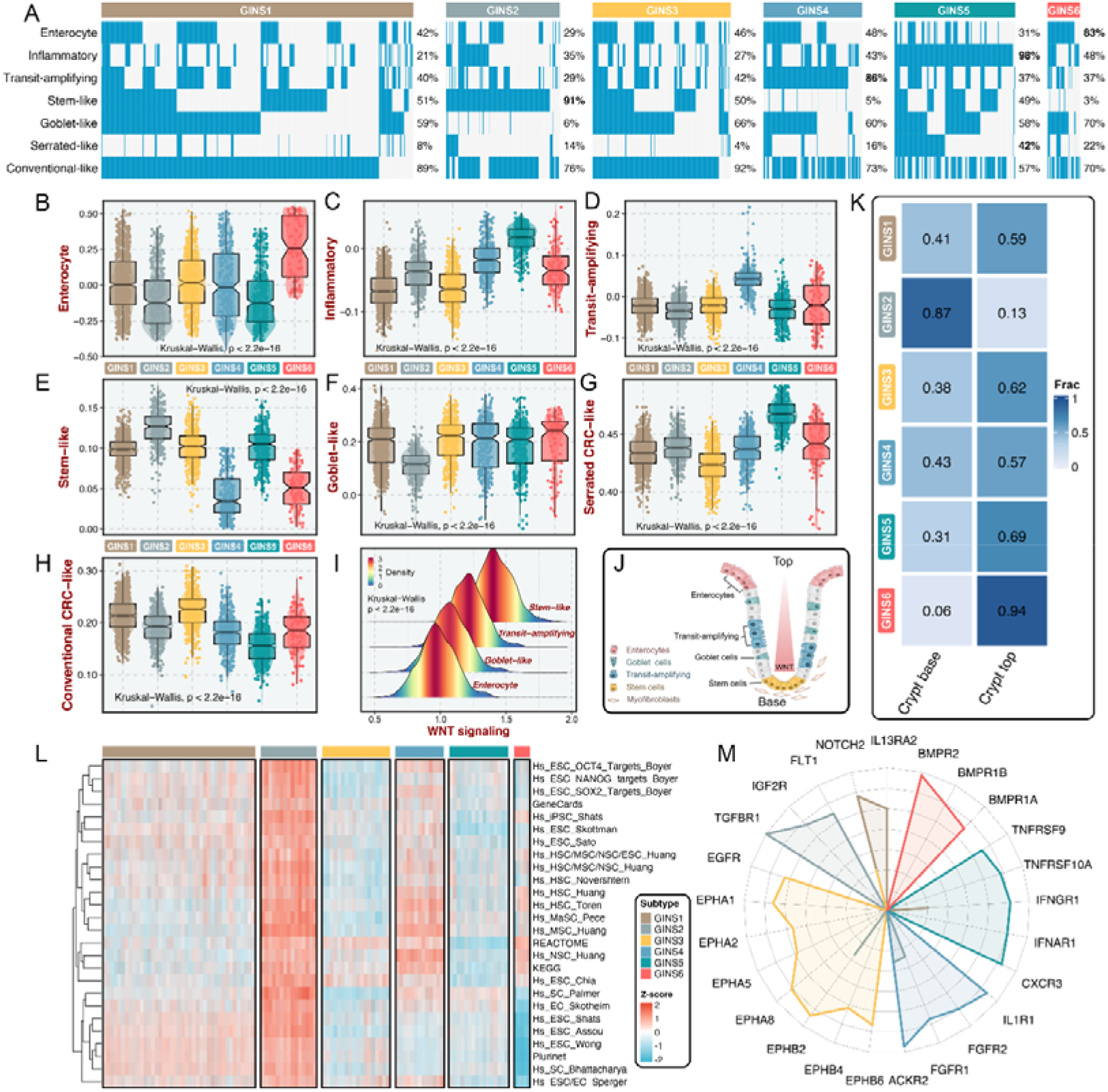
GINS subtypes were associated with cellular phenotypes and autocrine loops. **A.** Supervised approach identified phenotype origins peculiar to individual GINS classes. **B-H.** Unsupervised-based GSVA showed the distribution of enterocyte (**B**), inflammatory (**C**), transit-amplifying (**D**), stem-like (**E**), goblet-like (**F**), serrated CRC-like (**G**), conventional CRC-like (**H**) scores among six subtypes. Kruskal-Wallis test. **I.** The distribution of WNT signaling score in different cell-like tumors. Kruskal-Wallis test. **J.** CRC cellular phenotypes correlated with colon-crypt location and WNT signaling. **K.** Fractions of the crypt base and top phenotypes among six subtypes. Nearest template prediction (NTP) algorithm based on published signatures assigned each sample into the crypt base and top phenotypes in the discovery cohort. **L.** GSVA analysis revealed that GINS2 displayed superior stemness abundance relative to other subtypes. **M.** Radar plot showed autocrine stimulation loops in GINS subtypes.

Using SSEA, we also assessed the biological significance of each subtype in mitogenic/anti-apoptotic autocrine loops(***Isella et al., 2017***), as a proxy of growth factor-dependent oncogenic signaling (***Supplementary File 13***). All samples in SSEA framework were ranked according to ‘receptor activation index’, which were computed by averaging the expression of receptor and its ligands. As results, GINS1 was mainly associated with elevated *NOTCH2* and *IL13RA2* autocrine stimulation loops (***Figure 8M and Figure 8-figure supplement 1***). GINS2 displayed high intrinsic *TGFBR1* and *IGF2R* stimulation (***Figure 8M and Figure 8-figure supplement 1***). Of note, GINS3 was characterized by activations of ephrin receptors (*EPHA* and *EPHB* signaling) (***Figure 8M and Figure 8-figure supplement 1***), a set of receptors that are activated via binding to Eph receptor interacting proteins and form the largest subfamily of receptor tyrosine kinases (RTKs). In line with prior findings, GINS3 featured a high *EGFR* activity (***Figure 8M and Figure 8-figure supplement 1***), corresponding to its sensitive response to cetuximab. GINS4 exhibited marked traits of high activities in *ACKR2*, *FGFR1*, and *IL1R1* stimulation loops (***Figure 8M and Figure 8-figure supplement 1***). Accordingly, our results attributed immune-related autocrine loops including *CXCR3*, *IFNAR1*, *IFNGR1*, *TNFRSF9*, and *TNFRSF10A* to GINS5 tumors (***Figure 8M and Figure 8-figure supplement 1***), concordant with inflammatory traits of this subtype. GINS6 was linked to *BMP* activity (***Figure 8M and Figure 8-figure supplement 1***), which was reported to restrict stem cell expansion and upregulated at the crypt top with a decreasing gradient towards the crypt base(***Kosinski et al., 2007***). Taken together, these findings further provided a higher resolution of GINS taxonomy.

### Multi-omics alteration characteristics of six subtypes

To identify the genetic traits peculiar to individual GINS subfamilies, we characterized the multi-omics landscape in the TCGA-CRC cohort (***Figure 9A***). *PIK3CA* mutations could activate *PI3K/AKT* signaling and further enhance the proliferation and invasion of cancer cells(***Raskov et al., 2020***), which was prevalent in GINS1 (45%) (***Figure 9A***). GINS2 enriched plentiful *SMAD4* mutations (53%), which was strikingly higher than background mutations of *SMAD4* in CRC(***Raskov et al., 2020***) (***Figure 9A***). As previously reported, *KRAS* mutations are widespread in CRC(***Raskov et al., 2020***), but to a lesser extent in GINS3 (12%) (***Figure 9A and Figure 9-figure supplement 1A***), in line with its inferior activity of *KRAS* signaling detected in the discovery cohort. GINS5 was previously identified as tumors with high TMB and MSI, and thus displayed overall rich mutations in driver genes (***Figure 9A***), especially *BRAF* (***Figure 9-figure supplement 1B***), which was associated with CRC showing a high level of MSI. Conversely, GINS5 presented low chromosomal instability (CIN), featured by slight copy number variation (CNV), whereas an evident CIN phenotype was assigned to GINS3 that possessed heavy CNV burden, including amplifications and deletions (***Figure 9A-D***). We also observed that the broad amplifications of Chr20 were particularly evident for GINS3 (***Figure 9E-F***). Liu et al. demonstrated that tumors with TMB-high and CNV-low showed favorable response to immunotherapy(***Liu et al., 2019***), further validating the enhanced remission potential for immunotherapy in GINS5. Subsequently, we identified four CpG island methylator phenotypes (CIMP) from the TCGA-CRC cohort using the beta value of 5000 CpG island promoters with the most variation (***Figure 9-figure supplement 1C***). As previously reported, high CIMP (CIMP-H) was parallel with high MSI (MSI-H)(***Raskov et al., 2020***), and our results consistently displayed that tumors with MSI-H or CIMP were predominantly assigned to GINS5 (***Figure 9G***). In this study, we determined seven DNA methylation-driven genes via our introduced pipeline (***Figure 9A and Supplementary Methods***). Specifically, the methylation silencing of *SMOC1* was strongly enriched in GINS3, and *TMEM106A* silencing prevalently occurred in GINS4 (***Figure 9A***). The expression levels of these two genes were significantly inversely correlated with their methylation levels (***Figure 9H-I***). Collectively, these findings suggested that GINS subtypes were endowed with specific genetic alterations that presumably drive biological characteristics.

**Figure 9.**
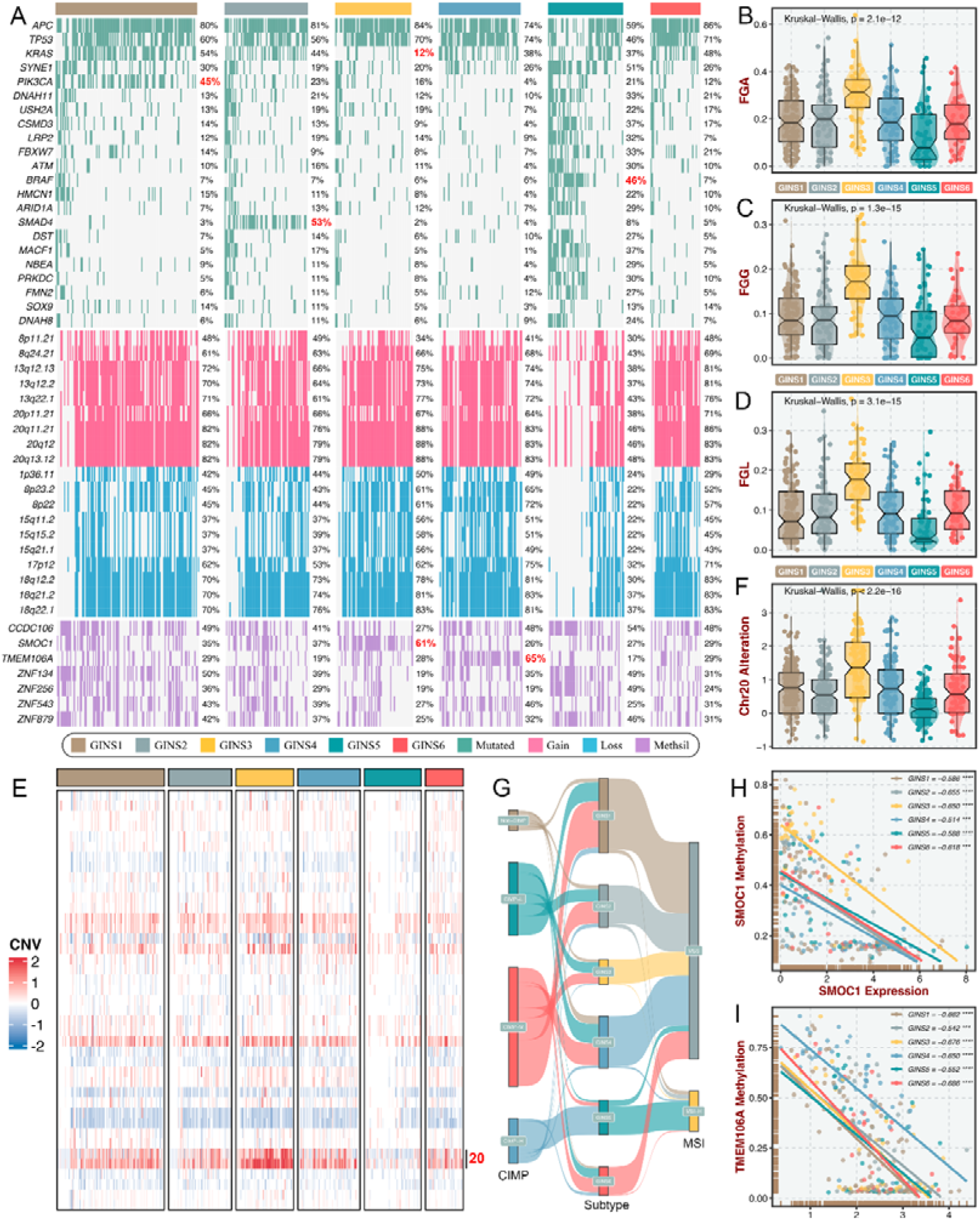
Multi-omics alteration characteristics of six subtypes. **A.** Genomic alteration landscape according to GINS taxonomy. The mutational genes were selected based on mutation frequency >10% and MutSigCV q-value <0.05. The focal gain and loss regions were selected based on CNV frequency >40% and GISTIC q-value <0.05. The methylation silencing (Methsil) genes were identified based on our pipeline (***Supplementary Methods***). **B-D.** The distribution of fraction of genome alteration (FGA), fraction of genome gained (FGG), and fraction of genome lost (FGL) among six subtypes. Kruskal-Wallis test. **E.** Heatmap showed the distribution of broad copy number changes across six subtypes in the TCGA-CRC dataset. **F.** The distribution of Ch20 alterations in six subtypes. Kruskal-Wallis test. **G.** Sankey plot showed connections among GINS subtypes, CIMP phenotypes, and MSI phenotypes. **H-I.** The expression levels of *SMOC1* and *TMEM106A* were significantly inversely correlated with their methylation levels. \*\*\**P* <0.001, \*\*\*\**P* <0.0001.

### Stromal contribution to the subtype transitions

The tumor transcriptome originated from cancer cells and TME, thus, it is conceivable that stromal components might impact the subtype assignments of CRC. Previous reports suggested that the subtype derived from stromal contents is a strong indicator of tumor aggressiveness and poor prognosis(***Isella et al., 2017***; ***Isella et al., 2015***), which was consistent with the inherent characteristics of stromal-derived GINS2 subtype. Indeed, most of GINS2-discriminant genes from the PAM classifier belonged to stromal genes (71.1%), followed by GINS4 (47.5%) (***Figure 10-figure supplement 1A***). To explore the extent of stromal contribution to the GINS subclasses, we leveraged the transcriptional profiles from CRC patient-derived xenografts (PDXs), for which the transcriptome is a mixture of human RNAs (deriving from cancer cells) and mouse RNAs (deriving from stromal cells) (***Figure 10A***). Hence, the stromal transcriptome of PDX samples can’t be detected by human microarray or RNA-seq^44^. In Uronis cohort (chip data)(***Uronis et al., 2012***) with 27 matched human CRC samples and PDX derivatives, the subtype assignments were incongruent between PDXs and their original counterparts. Subtypes with rich stromal components (e.g., GINS2 and GINS4) in human CRC samples were inclined to transform into subtypes with high tumor purity (e.g., GINS1 and GINS3) in corresponding PDX derivatives (***Figure 10B***). Another RNA-seq cohort with larger samples, Isella cohort(***Isella et al., 2017***), including 116 matched liver metastatic CRC and mouse xenografts, further validated these findings (***Figure 10C***).

**Figure 10.**
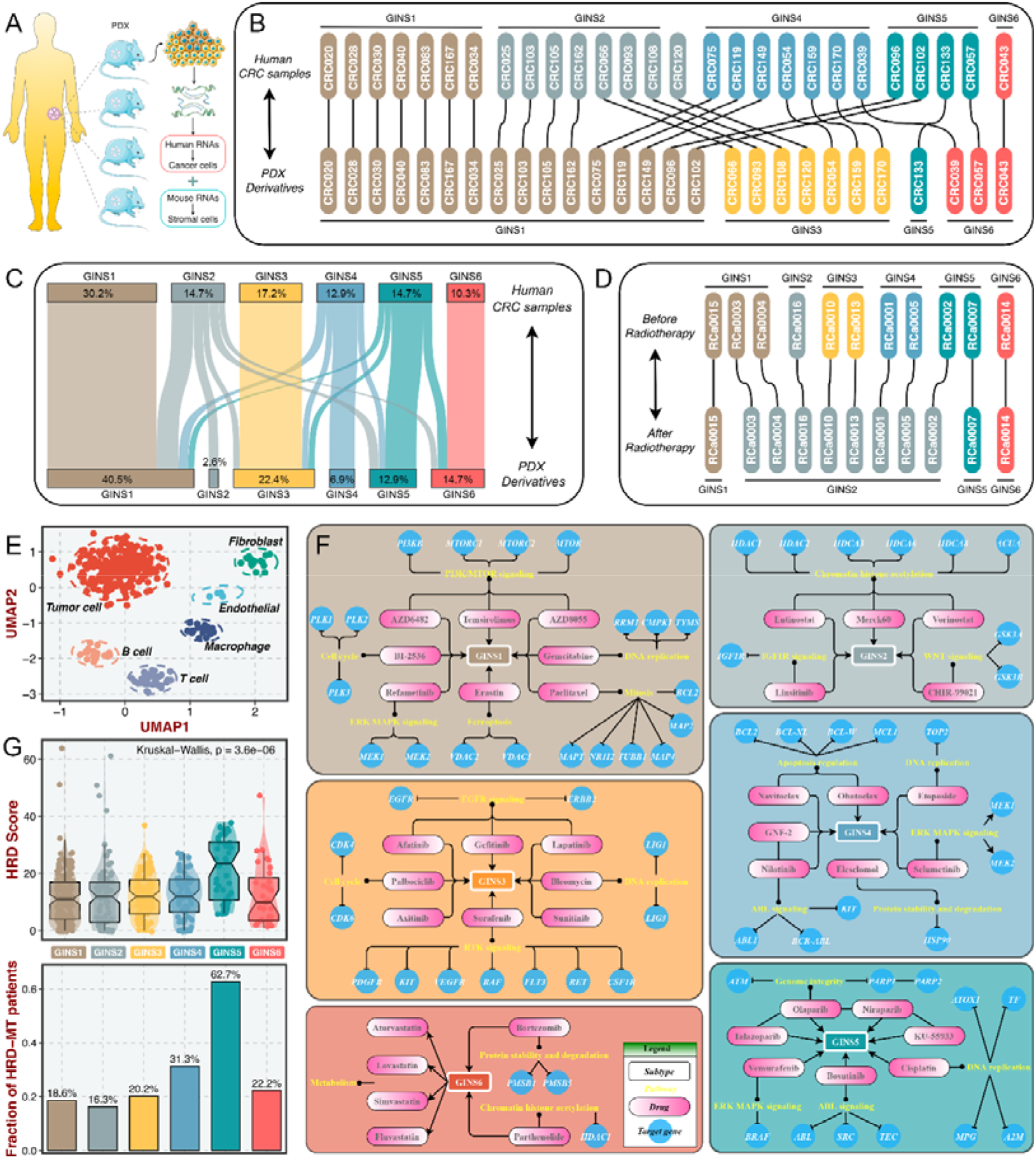
Stromal contributions and potential therapeutic agents. **A.** Schematic diagram showed the PDX transcriptome is a mixture of human RNAs (deriving from cancer cells) and mouse RNAs (deriving from stromal cells). **B-C.** Transcriptional classification of paired human CRC samples and PDX derivatives in Uronis cohort (**B**) and Isella cohort (**C**). **D.** Transcriptional classification of paired samples before and after radiotherapy. **E.** UMAP projected all cells annotated by reference component analysis (RCA) in spatial distribution. **F.** Identification of potential therapeutic agents for six subtypes. **G.** The distribution of HRD score and HRD pathway mutations among six subtypes. Kruskal-Wallis test.

Furthermore, a dataset (GSE56699) comprised 11 pairs of preoperative radiotherapy specimens and matched post-treatment surgical specimens(***Isella et al., 2015***), was utilized to investigate how the substitution of cancer cells by fibrous tissue, a typical reparative reaction triggered by radiotherapy, impacted the subtype assignments. We observed that the pretreatment specimens were confidently assigned to six subtypes, and most of the matched post-treatment biopsies were assigned to GINS2 (***Figure 10D***), confirming that stromal component served as the driven factor for GINS2 transitions. Additionally, in a single cell RNA-seq cohort derived from 11 patients with CRC(***Li et al., 2017***), UMAP projected all cells annotated by reference component analysis (RCA) in spatial distribution (***Figure 10E***). We applied the PAM-centroid distance classifier to perform subtype assignments for these 11 samples (***Figure 10-figure supplement 1B-C***). GINS2 displayed a strikingly higher fraction of fibroblasts relative to other subtypes (***Figure 10-figure supplement 1D***). A previous study reported that tumors with a high level of fibroblasts were resistant to radiotherapy(***Isella et al., 2015***). Thus, we further examined the associations between the GINS subtypes and radiotherapy in GSE56699. As expected, GINS2 possessed superior cancer-associated fibroblasts (CAF) score and worse prognosis across all treated samples (***Figure 10-figure supplement 2A-B***). Isella et al. demonstrated that CAF score was a stronger indicator of negative prognosis(***Isella et al., 2015***). In GINS2 samples, a higher CAF score certainly predicted worse prognosis (***Figure 10-figure supplement 2C***). We also observed that resistant tumors were predominantly enriched in GINS2 (***Figure 10-figure supplement 2D***). In summary, stromal signals remarkably contributed to the transitions of GINS2, which featured rich fibrous components and was resistant to radiotherapy.

### Identification of potential therapeutic agents for six subtypes

To facilitate the subtype-based targeted interventions, we introduced an integrated pipeline to identify potential therapeutic agents for each subtype(***Yang et al., 2021***) (***Figure 10-figure supplement 3A and Supplementary Methods***). Three pharmacogenomic datasets, CTRP, PRISM, and GDSC, store large-scale drug response and molecular data of human cancer cell lines, enabling accurate prediction of drug response in clinical samples(***Yang et al., 2021***). As mentioned above, stromal components could obscure the expression patterns of cancer cells in clinical samples. A purification algorithm termed *ISOpure*(***Quon et al., 2013***) was adopted to eliminate the contamination of stromal signal in the discovery cohort prior to conducting drug response prediction, and further yielded a purified tumor expression profiles comparable to cell lines(***Yang et al., 2021***). After purification, the proportion of stromal-rich subtypes (e.g., GINS2 and GINS4) was obviously decreased, suggesting the impact of stroma components had been eliminated (***Figure 10-figure supplement 3B***). A PDX dataset, GSE73255(***Isella et al., 2017***), is naturally uncontaminated by human stromal components. Hence, we tested our pipeline in the discovery cohort and GSE73255, and ultimately identified intersecting subtype-specific agents in two datasets (***Figure 10-figure supplement 3A***). To demonstrate the stability of drug response assessment, we examined whether the estimated response of four *EGFR* pathway inhibitors was concordant with their clinical efficacy—with a stronger clinical benefit in *KRAS*-mutant patients(***Raskov et al., 2020***). Our results indicated that patients with *KRAS* mutations squinted towards possessing a lower drug response (***Figure 10-figure supplement 3C***), in line with how *EGFR* pathway inhibitors behaved clinically(***Raskov et al., 2020***).

Taking the intersections of two datasets, we determined 41 specific-subtype agents for six subtypes (***Figure 10-figure supplement 4 and Supplementary File 14***). Interestingly, the targeted pathways of several candidate drugs were consistent with the biological and genomic peculiarities of corresponding subtypes (***Figure 10F***). For example, GINS1-specific drugs, BI-2536, gemcitabine, and paclitaxel target proliferative pathways; AZD6482, AZD8055 and temsirolimus target activated *PI3K*/*mTOR* signaling arise from *PIK3CA* mutations, which was strikingly harbored in GINS1; GINS2-specific drugs, linsitinib targets *IGF1R* signaling and CHIR-99021 targets WNT signaling; six *EGFR*/*RTK* signaling inhibitors, afatinib, gefitinib, lapatinib, axitinib, sorafenib, and sunitinib were specifically designed for GINS3; GINS6 featured dysregulated lipid synthesis, which might be targeted by four anticholesterol drugs containing atorvastatin, lovastatin, simvastatin, and fluvastatin. These results not only identified candidate compounds for each subtype, but also supported our previous findings. Notably, we observed that four *PARP* inhibitors were specific for GINS5, including olaparib, niraparib, talazoparib, and KU-55933 (***Figure 10F***). Previous reports have demonstrated that tumors with homologous recombination deficiency (HRD) are sensitive to *PARP* inhibitors(***Liu et al., 2021***). In the TCGA-CRC cohort, we next compared the HRD score and the proportion of HRD pathway mutations(***Liu et al., 2021***) among six subtypes. As expected, tumors with higher HDR score and mutations were predominantly assigned to GINS5 (***Figure 10G***), suggesting its stronger potential to benefit from *PARP* inhibitors. Overall, we provided more subtype-based targeted interventions for GINS taxonomy.

## Discussion

To address the snapshot effect of transcriptional analysis(***Chen et al., 2021***; ***Sahni et al., 2015***; ***Li et al., 2019***), we leveraged a relatively stable gene interaction network to discover the heterogeneous subtypes of CRC from an interactome perspective. As previously reported, biological networks maintain relatively constant irrespective of time and condition, preferably characterizing the biological state of bulk tissues(***Chen et al., 2021***; ***Sahni et al., 2015***; ***Li et al., 2019***). In the biological network, gene interactions are highly conservative in normal samples but broadly perturbed in diseased tissues(***Sahni et al., 2015***). Here, we constructed a large-scale interaction perturbation network from 2,167 CRC tissues and 308 normal tissues, deciphering six GINS subtypes with particular clinical and molecular peculiarities. Notably, although the GINS subtypes were dramatically associated with published classifications, only a limited overlap between our classifier genes with the signature genes of all previous classifications, suggesting a significant molecular convergence but also distinct specialties.

Considering that the stability and reproducibility of molecular subtypes are fundamental for clinical application, the GINS taxonomy was rigorously validated in 19 external datasets (n =3,420) with distinct conditions. Our six subtypes not only maintained comparable proportions, but also shared analogical transcriptional and clinical traits in the discovery cohort and 19 validation datasets. To provide a rapidly accessible clinical tool, we shrunk the 289-gene centroid-based classifier into a 14-gene random-forest miniclassifier. The qPCR results from 214 clinical CRC samples further demonstrated the 14-gene miniclassifier could afford the stability and interpretation in clinical practice.

Importantly, the GINS taxonomy also conveyed clear biological and molecular interpretability and laid a foundation for future clinical stratification and subtype-based targeted interventions (***Figure 11***).

**Figure 11.**
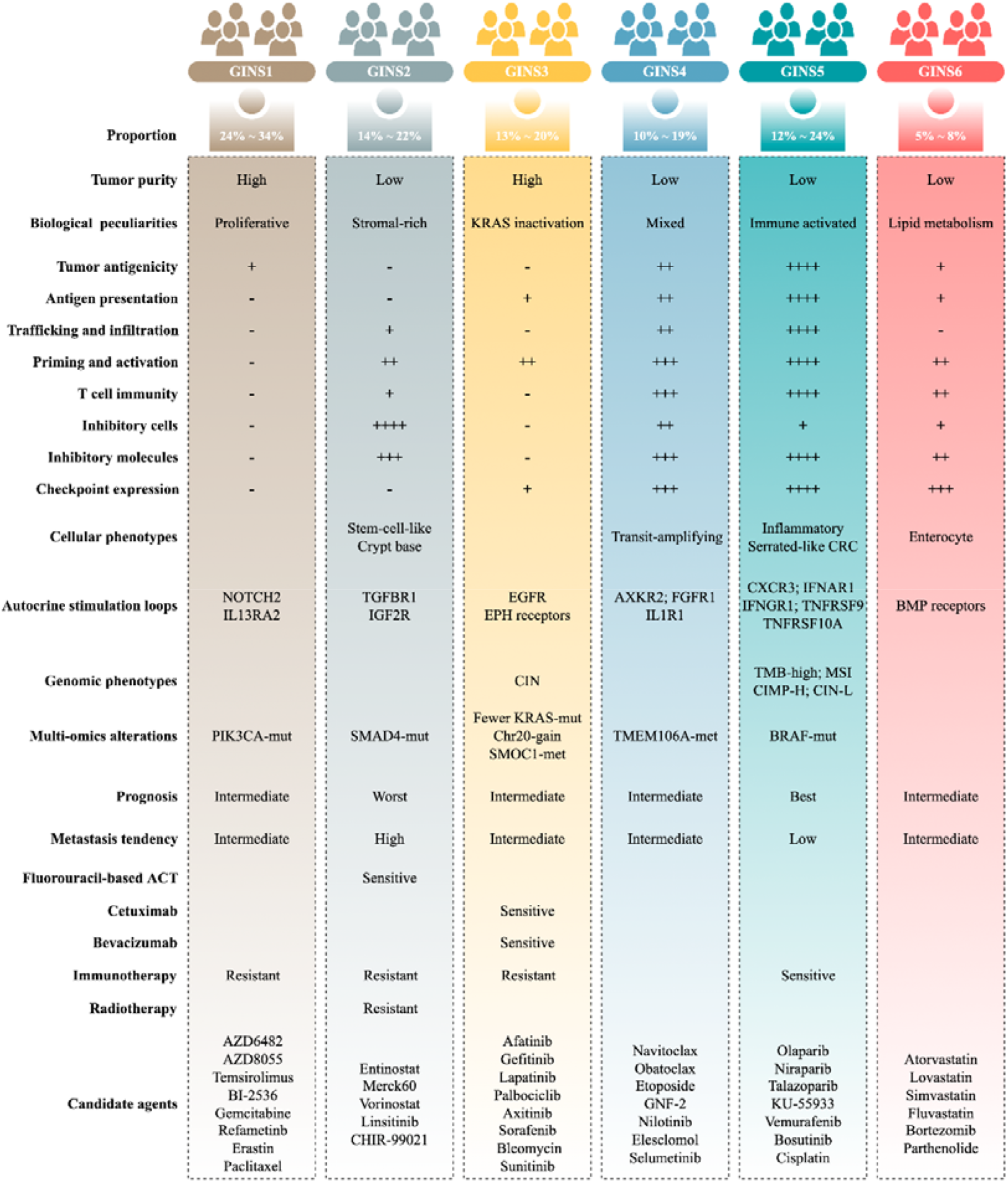
Summary of the main characteristics of six GINS subtypes.

GINS1, a proliferative subtype (24%∼34%), is endowed with elevated proliferative activity, high tumor purity, immune-desert, and *PIK3CA* mutations. This subtype displays a moderate malignant phenotype in spite of the high tumor purity, coincident with a previous study(***Mao et al., 2018***). In addition, GINS1 tumors reasonably develop resistance to immunotherapy due to their lower TMB and immune-desert TME. Indeed, current findings didn’t reveal an effective intervention for GINS1 in clinical settings. To further improve clinical outcomes of this subtype, we identified eight potential therapeutic agents for GINS1, including BI-2536/gemcitabine/paclitaxel targeting proliferative pathways, AZD6482/AZD8055/temsirolimus targeting activated *PI3K*/*mTOR* signaling arise from *PIK3CA* mutations, refametinib, and erastin.

GINS2, a stromal-rich subtype, is characterized by abundant fibrous content, immune-suppressed, stem-cell-like, *SMAD4* mutations, unfavorable prognosis, and high potential of recurrence and metastasis. In line with previous studies(***Guinney et al., 2015***; ***Isella et al., 2017***; ***De Sousa et al., 2013***; ***Sadanandam et al., 2013***; ***Marisa et al., 2013***; ***Isella et al., 2015***), CRC patients with stem and mesenchymal transcriptional traits squint towards displaying the malignant phenotypes. Notably, stromal contents remarkably contributed to the transitions of GINS2 into other subtypes, suggesting PDX or cell lines are not applicable surrogates for assessing the GINS taxonomy(***Sadanandam et al., 2013***). For GINS2 tumors, patients are suitable for fluorouracil-based ACT but resistant to radiotherapy due to a high level of fibroblasts(***Isella et al., 2015***). Unlike GINS1/3, the immunotherapeutic resistance of GINS2 is mainly due to highly infiltrating immunosuppressive cells, such as fibroblasts, MDSC, Treg cells, and M2 macrophages, and is therefore dubbed as the immune-suppressed phenotype.

GINS3, a *KRAS*-inactivated subtype, was featured by high tumor purity, immune-desert, activation of *EGFR* and ephrin receptors, CIN, fewer *KRAS* mutations, and *SMOC1* methylation. This subtype is a typical wild-type *KRAS* subgroup, paralleling by the sensitivity to *EGFR*/*VEGFR* inhibitors, such as cetuximab and bevacizumab. Moreover, we also identified six *EGFR*/*RTK* signaling inhibitors, including afatinib, gefitinib, lapatinib, axitinib, sorafenib, and sunitinib, which could serve as additional supplements for routine agents. Different from GINS1/2, the immunotherapeutic resistance of GINS2 could be driven by sparse immune storage and high burden of CNV(***Liu et al., 2019***). As previously reported, CNV-high tumors tend to respond unfavorably to immunotherapy(***Liu et al., 2019***).

GINS4, a mixed subtype, is distinguished by moderate level of stromal and immune activities, transit-amplifying-like phenotype, and *TMEM106A* methylation. This subtype is deemed as the intermediate state of GINS2 and GINS5. Thus, further interventions should focus on how to convert GINS4 into GINS5 with better prognosis and sensitivity to immunotherapy, such as adoptive T-cell immunotherapy, cancer vaccine, and reprogramming the microenvironment(***Liu and Sun, 2021***).

GINS5, an immune-activated subtype, is associated with serrated-like CRC, stronger immune activation, plentiful TMB and NAL, MSI, and CIMP-H, *BRAF* mutations, and favorable prognosis. This subtype commonly exhibits decent clinical outcomes due to the stronger immune activation. Spontaneously, GINS5 tumors respond well to immunotherapy. We also documented that tumors with high HDR score and mutations were predominantly assigned to GINS5, suggesting its nonnegligible potential to benefit from *PARP* inhibitors(***Liu et al., 2021***). Indeed, four *PARP* inhibitors, including olaparib, niraparib, talazoparib, and KU-55933, were identified for more targeted or combined interventions for GINS5 tumors.

GINS6, a metabolic reprogramming subtype, is linked to accumulated fatty acids, enterocyte-like, and *BMP* activity. The lipid metabolisms are the most significant metabolic processes in GINS6. Also, the metabolomics results further demonstrated that GINS6 featured by abundant fatty acids metabolites, indicating GINS6 tumors could be intervened by metabolic inhibitors targeting fatty acid metabolisms. Interestingly, our pipeline determined four anticholesterol drugs containing atorvastatin, lovastatin, simvastatin, and fluvastatin, were specific for GINS6. Statins have been reported to attenuate cellular energy and outgrowth of cancers(***Ali et al., 2019***; ***Beckwitt et al., 2018***), might provide extra options for this subtype.

From an interactome perspective, our study identified and diversely validated a high-resolution classification system, which could confidently serve as an ideal tool for optimizing decision-making for patients with CRC. The multifariously biological and clinical peculiarities of GINS taxonomy improve the understanding of CRC heterogeneity and facilitate clinical stratification and individuation management. Additionally, candidate specific-subtype agents provide more targeted or combined interventions for six subtypes, which also need to be validated in clinical settings. In this study, the GINS taxonomy could be measured and reproduced using a simple PCR-based assay, making it attractive for clinical translation and implementation. Nevertheless, a prospective multicenter study is still imperative to further confirm the biological and clinical interpretability of six subtypes. To conclude, we believe this novel high-resolution taxonomy could facilitate more effective management of patients with CRC.

## Materials and methods

### Data source and specimen collection

A total of 6,216 patients and 308 normal samples were enrolled from public databases. A merged discovery cohort consisted of 19 datasets (n =2,167), another 19 independent datasets (n =3,420) were used for validation, and 17 datasets including immunotherapy or radiotherapy annotations, cancer cell lines, patient-derived xenografts (PDX), and single-cell sequencing were applied for exploration. ***Supplementary File 15*** summarizes the data sources and details of this study. We also enrolled 214 clinical CRC samples from The First Affiliated Hospital of Zhengzhou University for further validation (***Supplementary File 3***).

### Construction of the gene interaction-perturbation network

Our gene interaction-perturbation pipeline applied the discovery cohort (n =2,167) composed of 19 independent datasets from the same chip platform (Affymetrix Human Genome U133 Plus 2.0 Array, GPL570) as the tumor sample input and the GTEx cohort (n =308) as the normal sample input (***Figure 1, Supplementary Methods and Supplementary File 15***). A pathway-derived analysis requires constructing the protein interaction functional networks projected by candidate genes(***Chen et al., 2021***). The initial background network from the Reactome database(***Jassal et al., 2020***) included 6,376 genes and 148,942 interactions, and fitted the biological scale-free network distribution in the discovery cohort (*R* =-0.852, *P* <2.2e-16; ***Figure 1-figure supplement 1A***). The perturbation degree of gene interactions in the background network could measure the biological state of individual patients(***Chen et al., 2021***). The global network perturbations are quantified via the interaction change of each gene pair, which is reasonably and effectively utilized to characterize the pathological condition at the individual level(***Chen et al., 2021***; ***Sahni et al., 2015***). In high-throughput profiles, we need to compute the relative perturbations of all gene pairs based on the benchmark vector. Since gene interactions are highly conservative within normal samples, we selected the average interactions of all normal samples as the benchmark vector. Gene interactions in each patient should be compared with the benchmark network, and the corresponding difference represents the gene interaction perturbation for that patient. Indeed, tumor samples displayed remarkably stronger perturbations relative to normal samples (*P* <2.2e-16; ***Figure 1-figure supplement 1B***). The interaction perturbations of normal samples were much denser and less than tumor samples (***Figure 1-figure supplement 1C***). Collectively, 92.6% of all 148,942 gene pairs exhibited more dispersion in tumor samples than in normal samples by comparing the coefficient of variation (CV) of interaction perturbations (*P* <2.2e-16; ***Figure 1-figure supplement 1D***). These results revealed that the interaction perturbations of normal samples were more stable, whereas a broader variation existed in tumor samples, making it possible to discover heterogeneous subtypes in CRC samples.

### Statistical analysis

The detailed methods and statistics were described in ***Supplementary Methods***. All data processing, statistical analysis, and plotting were conducted in R 4.0.5 software. All statistical tests were two-sided. *P* <0.05 was regarded as statistically significant.

## Additional information

## Acknowledgements

Not applicable.

## Additional Files

### Supplementary Methods

#### Publicly available data generation and processing

##### Human CRC bulk cohorts

The integrated discovery cohort contained 2,167 patients with CRC from 19 independent datasets who fulfilled the following criteria: (1) available expression profiles obtained using the same chip platform (Affymetrix Human Genome U133 Plus 2.0 Array, GPL570) with raw CEL data; (2) overall survival (OS), relapse-free survival (RFS), fluorouracil-based adjuvant chemotherapy (ACT) or bevacizumab treatment information available. Another 19 independent datasets comprised 3,420 patients with CRC from different platforms and sequencing techniques (microarray or RNA-seq) were used for validation.

##### Normal tissue data

The Genotype-Tissue Expression (GTEx, https://gtexportal.org) project has established a tissue biobank with all samples from normal tissues rather than paracancerous tissues (e.g., TCGA). The RNA-seq data of 308 normal colorectal samples were obtained from the GTEx portal.

##### Immunotherapy cohorts

Three datasets (n =414) with immunotherapeutic annotations and expression profiles were derived from the following studies: Hugo and colleagues(***Hugo et al., 2016***) (GSE78220, n =27), Rose and colleagues(***Rose et al., 2021***) (GSE176307, n =89) and Mariathasan and colleagues(***Mariathasan et al., 2018***) (IMvigor210, n =298). Patients in Hugo cohort were treated with *PD-1* blockade, patients in Rose cohort were treated with *PD-1*/*PD-L1* blockade, and patients in Mariathasan cohort were treated with *PD-L1* blockade. All patients were performed with next-generation sequencing.

##### Multi-omics data for TCGA-CRC

The TCGA-CRC multi-omics data, including RNA-seq (raw count), proteome (Reverse Phase Protein Array), HumanMethylation450 array, whole-exome sequencing (VarScan MAF files), and copy number variation (CNV) data, were derived from TCGA portal (https://portal.gdc.cancer.gov). The neoantigen load (NAL) and homologous recombination deficiency (HRD) score measured by TCGA officials were available in TCGA pan-cancer atlas(***Thorsson et al., 2018***).

##### CRC cell lines

The Cancer Cell Line Encyclopedia (https://sites.broadinstitute.org/ccle, CCLE) stores the processed multi-omics data of cancer cell lines. We retrieved 55 CRC cell lines with both transcriptome and metabolomics data (including 225 metabolites) for further explorations(***Li et al., 2019***).

##### PDX model data

Two CRC PDX datasets were generated from two previous studies(***Isella et al., 2017***; ***Uronis et al., 2012***). Uronis cohort (chip data) encompassed 27 matched human primary CRC and mouse xenografts and Isella cohort (RNA-seq data) included 116 matched liver metastatic CRC and mouse xenografts.

##### Radiotherapy cohort

The GSE56699 comprises 72 primary rectal cancer with radiotherapeutic annotations, including 11 pairs of preoperative radiotherapy specimens and matched post-treatment surgical specimens(***Isella et al., 2015***).

##### Single-cell RNA sequencing data

A total of 1,591 single cells from 11 patients with CRC using Fluidigm based single-cell RNA-seq protocol were achieved from Li et al. study(***Li et al., 2017***). The cell annotations were obtained from the reference component analysis (RCA).

##### Pharmacogenomic datasets

Drug response and molecular data of human cancer cell lines were generated from the Cancer Therapeutics Response Portal (CTRP, https://portals.broadinstitute.org/ctrp), Profiling Relative Inhibition Simultaneously in Mixtures (PRISM, https://depmap.org/portal/prism), and Genomics of Drug Sensitivity in Cancer (GDSC, https://www.cancerrxgene.org) datasets. The area under the dose-response curve (AUC) values measured drug sensitivity (the AUC values in CTRP range from 0 to 30, in PRISM and GDSC range from 0 to 1). AUC values were log-transformed, with lower values indicating increased sensitivity to treatment. Drugs with missing AUC values in more than 20% of cell lines were removed, and the remaining missing values were imputed using the k-nearest neighbors (KNN) imputation procedure. The hematopoietic and lymphoid tissue-derived cell lines were excluded. Ultimately, CTRP contains the response data for 481 drugs over 835 cell lines, PRISM contains the response data for 1,448 drugs over 482 cell lines, and GDSC contains the response data for 265 drugs over 828 cell lines.

##### Data processing

Published microarray data were obtained from the Gene Expression Omnibus (GEO, https://www.ncbi.nlm.nih.gov/geo). The robust multiarray average (RMA) preprocessing and normalization of raw CEL files from Affymetrix GeneChip® arrays were performed via *affy* package. Gene expression presence and absence were measured via the barcode algorithm(***McCall et al., 2011***), and genes not presented in all samples were discarded. To remove potential multicenter batch effects of discovery cohort, data were corrected using ComBat approach implemented in *sva* package (***Leek et al., 2012***). For Illumina and Agilent microarrays, the processed data were obtained from the authors. The RNA-seq data (raw count) from TCGA-CRC and GTEx were converted to transcripts per kilobase million (TPM) and further log_2_ transformed. For somatic mutation data, somatic variants were detected using TCGA VarScan2 pipeline. A binary mutation profile was used to indicate the presence or absence of a gene mutation, and then gene-level mutations were generated after filtering out the synonymous mutations. For CNV data, segmented CNV values were generated by circular binary segmentation algorithm implemented in *DNAcopy* package. Then, all genes were mapped onto this segmented data to obtain gene-level and broad-level CNV values using GISTIC2.0. For DNA methylation data, we firstly excluded probes with >20% of missing values across samples and the probes for sex chromosomes, and the remaining missing values were imputed using the KNN imputation procedure. Then, each gene with the average beta value of the promoter region probes was assigned the final methylation values. For drug outcomes, the RECIST guideline (version 1.1)(***Eisenhauer et al., 2009***) indicates treatment responders and non-responders are defined as having a complete response (CR)/partial response (PR) and stable disease (SD)/progressive disease (PD), respectively.

#### Description of previous CRC classifications

To compare our subtypes with previously reported CRC classifications, the discovery cohort was reclassified according to the previous subtype criteria, including consensus molecular subtypes (CMS)(***Guinney et al., 2015***), CRC intrinsic subtypes (CRIS)(***Isella et al., 2017***), colon cancer subtypes (CCS)(***De Sousa et al., 2013***), CRCAssigner (CRCA)(***Sadanandam et al., 2013***), and Cartes d’Identité des Tumeurs (CIT)(***Marisa et al., 2013***), respectively. CMS intergrated six independent classfications systems into four CRC groups (CMS1-4) based on gene transcription and are prognostic in early-stage and first-line metastatic settings. Isella et al. deployed human-specific expression profiling of CRC PDXs to assess cancer-cell intrinsic transcriptional features, and identified five CRIS subtypes (CRISA-E). CCS was developed using a previously developed unsupervised consensus-based clustering technique, and formed three clusters generated the most robust classification (CCS1-3). Sadanandam et al. discovored five CRCA subtypes (CRCA1-5) shared similarities to distinct cell types within the normal colon crupt and showed differing degrees of ‘stemness’ and Wnt signaling. The french national CIT program enrolled a a multicenter cohort of 750 patients with stage I to IV CRC and found six CIT subtypes (CIT1) based on cluster expression centroid classification.

#### Background network

The gene interaction-perturbation pipeline applies the discovery cohort as the tumor sample input and the GTEx cohort as the normal sample input. A pathway-derived analysis requires constructing the protein interaction functional networks projected by candidate genes(***Chen et al., 2021***). Data from the Reactome Pathway Database (https://reactome.org) focus on the reaction, which was utilized to establish the biological interaction networks(***Jassal et al., 2020***). Network nodes that absented in our cohorts were removed and existing nodes were integrated into the large background network with 148,942 interactions in total.

#### Interaction-perturbation-based program

The flowchart of this program is shown in ***Figure 1***. For each gene within the background network, we first calculated its rank in individuals based on its gene expression value in each sample (the smallest expression value corresponds to the minimum rank, and the largest expression value corresponds to the maximum rank). Thus, the expression matrix was converted into the rank matrix by ranking all genes according to the expression values in all samples. The *R_a,s_* represents the rank of gene *G_a_* in sample *s*. Subsequently, the delta rank matrix, which rows and columns denoted interactions in the background network and samples, is generated from the rank matrix. For instance, a delta rank (represented by δ*_i,s_*) was calculated by subtracting the ranks for each gene pair (e.g., *G_a_* and *G_b_*) connected by an interaction in the background network, as follows:

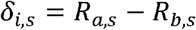

Where genes *G_a_* and *G_b_* are connected by interaction *i* in the background network.

Gene interactions are highly conservative within normal samples, and interaction perturbations are rare(***Sahni et al., 2015***). The within-sample delta ranks of gene pairs are highly stable among samples under normal conditions but are often widely disrupted after certain treatments, such as gene knockdown, gene transfection, drug treatment and tissue canceration(***Li et al., 2019***). Hence, we hypothesized that the background network is stable across all normal samples, and then the interactions within normal samples served as the benchmark network. We ranked genes according to their mean gene expression value among normal samples and similarly calculated the delta rank as the benchmark delta rank vector with elements denoted by 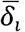, where *i* is an interaction in the background network. This vector measures the mean relative ranks of gene pairs in all normal samples. Gene interactions in each patient should be compared with the benchmark network, and the corresponding difference represents the gene interaction perturbation for that patient. Upon subtracting the benchmark delta rank vector from the delta rank of each sample, we finally obtained the interaction-perturbation matrix Δ with element Δ*_i,s_*. For an interaction *i* in the background network and an individual sample s,

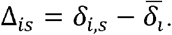

The interaction-perturbation matrix can measure the sample-specific interaction perturbation in the same whole background network effectively. Each column of the interaction-perturbation matrix represents the gene interaction perturbations for an individual sample, i.e., the sample-specific perturbation of the gene interaction.

#### Discovery of the gene interaction network-based subtypes

The subtypes were deciphered in the discovery cohort using the consensus clustering approach that required two elements, including features in rows and samples in columns. The clustering features needed to meet two aspects: (1) being able to significantly distinguish tumor from normal samples; (2) maintaining strong heterogeneity within all tumor samples with high variability. First, the differential analysis between tumor and normal samples for each interaction was performed. The top 30,000 interactions (approximately 20%) that differed significantly between tumor and normal samples and the top 30,000 interactions with high standard deviation (SD) among all CRC samples were selected. Subsequently, the intersection of two sets with 30,000 gene interactions over all CRC samples were retained for clustering work.

Furthermore, the perturbation matrix with the selected interactions was subjected to the consensus clustering procedure(***Wilkerson and Hayes, 2010***), a resampling-based clustering algorithm implemented in the *ConsensusClusterPlus* package(***Wilkerson and Hayes, 2010***). This procedure using the partitioning around medoids approach and 1-Spearman correlation distance was performed 5000 iterations on 80% of interactions and 80% of samples selected randomly. All derived partitions for a cluster *K* (2-10) were summarized by clustering the (samples x samples) co-classification matrix. Subsequently, the consensus score matrix, cumulative distribution function (CDF) curve, proportion of ambiguous clustering (PAC) score, and Nbclust were synthetically used to determine the optimal number of clusters. A higher consensus score between two samples indicates they are more likely to be grouped into the same cluster in different iterations. The consensus values range from 0 (never clustered together) to 1 (always clustered together) marked by white to dark brown. In the CDF curve of a consensus matrix, the lower left portion represents sample pairs rarely clustered together, the upper right portion represents those almost always clustered together, whereas the middle segment represents those with ambiguous assignments in different clustering runs. The proportion of ambiguous clustering (PAC) measure quantifies this middle segment; and is defined as the fraction of sample pairs with consensus indices falling in the interval (u1, u2) ∈ [0, 1] where u1 is a value close to 0 and u2 is a value close to 1 (for instance u1=0.1 and u2=0.9). A low value of PAC indicates a flat middle segment, and a low rate of discordant assignments across permuted clustering runs. PAC for each K is CDF_k_(u2) - CDF_k_(u1)(□enbabaoğlu etal., 2014). According to his criterion, we can therefore infer the optimal number of clusters by the K value having the lowest PAC. The Nbclust uses 26 mathematic criteria to select the optimal number.

#### Subtype Robustness

##### Internal robustness

(1) The subtypes were obtained using a consensus clustering procedure using both gene and sample resampling (5000 random subselections of 80% of the samples and 80% of the genes), such that these results are stable under conditions of gene and sample resampling. (2) The subtypes were obtained from a large set (n =2167) of samples processed with the same experimental procedure. (3) Moreover, we used several independent metrics to evaluate the performance of subtypes, such as the consensus score matrix, CDF curve, PAC score, and Nbclust.

##### External robustness

The GINS subtypes were validated on 19 independent datasets comprising 3,420 patients with CRC from different platforms and sequencing techniques (microarray or RNA-seq). Clinical and biological characteristics of the subtypes were found to be conserved in validation datasets.

#### Generation of the GINS classifier

To identify GINS subtypes in novel datasets using a small list of genes, we developed a gene expression-based classifier in three steps. First, our analysis included only samples that statistically belonged to the core of each subtype. Excluding samples with negative silhouette width has been shown to minimize the impact of sample outliers on the identification of subtype markers, as described in TCGA glioblastoma classification(***Verhaak et al., 2010***). Second, the significance analysis of microarrays (SAM)(***Tusher et al., 2001***) was employed to identify 762 genes significantly differentially expressed across the GINS subtypes. This is a well-established method that looks for large differential gene expression relative to the spread of expression across all genes. Sample permutation is used to estimate false discovery rates (FDR) associated with sets of genes identified as differentially expressed. The threshold was set to Benjamini–Hochberg-corrected FDR <0.0001. Third, the differentially expressed genes were further trained by prediction analysis for microarrays (PAM)(***Tibshirani et al., 2002***) to build a classifier. The PAM eliminated the contribution of genes that differentially express below a specific threshold, ΔPAM, relative to the subtype-specific centroids. The leave-one-out cross-validation (LOOCV) was performed to ultimately identify 289 genes that had the lowest misclassification error (1.8%). Using this strategy, a centroid-based classifier was built, and six centroids were computed on the gene-median centered discovery cohort. For each validation dataset, the distance to the six centroids of each sample was computed and samples were assigned to the closest centroid subtype. As previously reported(***Marisa et al., 2013***), the decision rule was based on the diagonal quadratic discriminant analysis method (DQDA) is defined as follows:

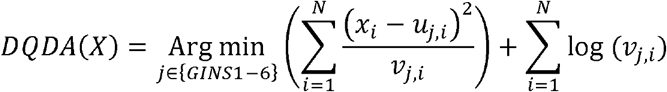

where *N* is the number of genes, *x* represents the expression normalized values, *u_j,t_* and *v_j,t_* were denoted by the mean and the variance of the gene *i* across samples of the subtype *j* from the discovery cohort. The classification confidence was assessed by identifying outliers (too distant samples) and mixed assignment samples (when a sample is close to several centroids). More specifically, a sample was defined as an outlier if its distance to the nearest centroid is superior to *n* times the median absolute deviation (MAD) of the distances of the samples used to compute the centroid; *n* is defined as the maximum (distances to centroid-median_distances to centroid_)/MAD_distances to centroid_. A sample has a mixed assignment if the difference of its distance to centroid is inferior to the 1st decile of the difference between centroids on data used to compute centroids.

#### Human tissue specimens

The human cancer tissues used in this study were approved by Ethnics Committee of The First Affiliated Hospital of Zhengzhou University and the TRN is 2019-KW-423. Overall, 214 frozen surgically resected CRC tissues were collected from The First Affiliated Hospital of Zhengzhou University. All patients provided written informed consent; received available standard systemic therapies; were aged 18 years or older; had adequate haematologic, renal, and liver function; had Eastern Cooperative Oncology Group performance status of 0 or 1; and had measurable disease according to Response Evaluation Criteria in Solid Tumours (RECIST, version 1.1)(***Eisenhauer et al., 2009***). Responders and nonresponders were defined as having a complete response (CR)/partial response (PR) and stable disease (SD)/progressive disease (PD), respectively. Detailed baseline data of patients with CRC are displayed in ***Supplementary File 3***.

#### RNA preparation and Quantitative Real-Time PCR (qRT-PCR)

Total RNA was isolated from CRC tissues using RNAiso Plus reagent RNA quality was evaluated using a NanoDrop One C (Waltham, MA, USA), and RNA integrity was assessed using agarose gel electrophoresis. An aliquot of 1 μg of total RNA was reverse transcribed into complementary DNA (cDNA) according to the manufacturer’s protocol using a High-Capacity cDNA Reverse Transcription kit (TaKaRa BIO, Japan). qRT-PCR was performed using SYBR Assay I Low ROX (Eurogentec, USA) and SYBR® Green PCR Master Mix (Yeason, Shanghai, China) to detect the expression of 14 genes. The expression value was normalized to *GAPDH*, and then log2 transformed for subsequent analysis. The primer sequences of the included 14 genes and *GAPDH* were shown in ***Supplementary File 4***.

#### Quantitative PCR miniclassifier

To facilitate translation of the GINS subtypes in clinical settings, we intended to develop a quantitative PCR (qPCR) miniclassifier and further validate our subtypes in 214 clinical CRC samples from The First Affiliated Hospital of Zhengzhou University.

Using 289 genes from the PAM classifier, we proposed a six-step pipeline to develop the simplified GINS miniclassifier in the discovery cohort:

1. For a gene, we extracted the expression of this gene in each subtype, and if the expression of one subtype was significantly higher than that of all other subtypes, this gene is defined as a specific gene of this subtype. In total, we identified 191 subtype-specific genes.
2. Bootstrapping method was further used to test these genes (from the first step). We extracted 70% samples randomly from the entire cohort and performed univariate Logistic regression analysis on these samples to assess the correlation between the gene expression and prognosis. This procedure was repeated 1000 times and 93 genes that were incorporated in all resample runs (achieved P < 0.05 in robustness testing) were kept for next step analysis.
3. To further select genes with the most information, the LASSO regression was applied for further dimension reduction. The LASSO algorithm is a popular method for regression with high-dimensional predictors. It introduces a penalty parameter λ to shrink some regression coefficients to exactly zero. The penalty parameter λ, called the tuning parameter, controls the amount of shrinkage: the larger the value of λ, the fewer the number of predictors selected. The 10-fold cross validation was used to determine the optimal values of λ via the one-standard-error rule. The optimal λ is the largest value for which the deviance is within one standard-error of the smallest value of deviance. In this process, 14 key genes were retained.
4. The discovery cohort was firstly divided into the training (70%) and testing (30%) datasets. With the expression profiles of 14 key genes, the random forest was employed to build a classifier with the following criteria: Gini impurity, 1,000 trees, minimal nodesize = 3, bootstrap sample, and 10-fold cross validation. The random forest analysis was repeated 1,000 times, and the model with the smallest of out-of-bag (OOB) error rate was considered as the optimal classifier.
5. The five indicators including accuracy, precision, recall, F1-score, and specificity were utilized to evaluate the performance of our miniclassifier in both training and testing datasets.

#### Gene set enrichment analysis

The *clusterProfiler* package(***Yu et al., 2012***) was utilized to perform downloaded from the gene set enrichment analysis (GSEA). The significance of enrichment was estimated using default settings and 1,000 gene permutations.

#### Gene set variation analysis

The gene set variation analysis (GSVA) was conducted via the *GSVA* package(***Hanzelmann et al., 2013***). GSVA is a non-parametric, unsupervised method for estimating variation of gene set enrichment through the samples of an expression data set. GSVA performs a change in coordinate systems, transforming the data from a gene by sample matrix to a gene-set by sample matrix, thereby allowing the evaluation of pathway enrichment for each sample.

#### Sample set enrichment analysis

To assess the extent to which six subtypes captured samples with stronger transcriptional signatures, we introduced a framework termed ‘Sample Set Enrichment Analysis’ (SSEA)(***Isella et al., 2017***). In SSEA, all samples are ranked by the integrated scores, and the ranked sample list was further subjected to the GSEA procedure to test whether the ‘sample set’ for each subtype enriches high-ranking samples. As previously reported(***Isella et al., 2017***), a signature score was computed for each signature and each sample, as follows:

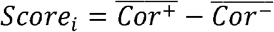

where 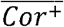 is the average expression value of the genes positively correlated with the subtype and 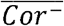 is the average expression value of the genes negatively correlated with the subtype in sample *i*.

For SSEA of mitogenic/anti-apoptotic autocrine loops(***Isella et al., 2017***), a total of 472 receptor/ligand interactions were downloaded from the Database of Ligand Receptor Partners (http://dip.doembi.ucla.edu/dip/DLRP.cgi). All samples in SSEA framework were ranked according to ‘receptor activation index’, which were computed by averaging the expression of receptor and its ligands, as follows:

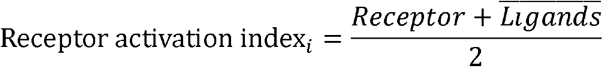

where *Receptor* is the mRNA expression of the receptor of interest and 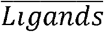 stands for the average expression of its corresponding ligands in sample *i*.

#### Cancer-immunity cycle

Based on the rationale that immunity within tumors is a dynamic process, Karasaki and colleagues(***Karasaki et al., 2017***) have proposed an immunogram for the cancer-immunity cycle (CIC) depicted by eight axes of the immunogram score (IGS): IGS1, T cell immunity; IGS2, tumor antigenicity; IGS3, priming and activation (actived dendric cell enrichment); IGS4, trafficking and infiltration; IGS5, recognition of tumor cells; IGS6, inhibitory cells; IGS7, checkpoint expression; and IGS8, inhibitory molecules. The gene sets of IGS1-IGS8 were retrieved from a previous study(***Karasaki et al., 2017***). The single [set enrichment analysis (ssGSEA) approach was leveraged to measure the IGS, and the immunogram radar displayed the normalized IGS value of six subtypes.

#### Assessment of immunotherapeutic potential

To predict the putative response to immune checkpoint blockade (ICB), CRC samples were scored using T-cell inflammatory signature (TIS) and Tumor Immune Dysfunction and Exclusion (TIDE) approaches. TIS proposed by Ayers et al. was used to predict clinical response to PD-1 blockade. The signature was composed of 18 inflammatory genes associated with antigen presentation, chemokine expression, cytotoxic activity, and adaptive immune resistance(***Ayers et al., 2017***). The TIDE algorithm (http://tide.dfci.harvard.edu/) integrates the expression signature of two primary mechanisms of immune evasions: T cell dysfunction and T cell exclusion, to model tumor immune evasion. Patients with higher TIDE score suggest the greater potential of tumor immune evasion; thus, these patients would derive worse immunotherapy response(***Jiang et al., 2018***). The immunophenoscore (IPS) was applied to assess the immune state of each sample. IPS is a scoring scheme that quantifies the immunogenicity of tumor samples using a variety of markers of immune response or immune tolerance. The higher the IPS z-score, the stronger the immunogenicity and immunotherapeutic potential of the sample(***Charoentong et al., 2017***).

#### Characteristics of multi-omics alterations

For mutation analysis, using the VarScan pipeline and MutSigCV algorithm(***Lawrence et al., 2013***), we retained genes with mutation frequency >10% and MutSigCV q-value <0.05. For copy number variation (CNV) analysis, the DNAcopy pipeline used Affymetrix SNP 6.0 array data to identify genomic regions that are repeated and infer the copy number of these repeats. The copy number values were further transformed into segment mean values, which were represented as log2 (copy-number/2). With −0.3 and 0.3 as cut-off points, genes were marked as deletion (<-0.3), neutral (−0.3∼0.3), and amplification (>0.3). GISTIC 2.0 software(***Mermel et al., 2011***) was applied to define the recurrently amplified and deleted regions. Subsequently, we retained focal segments with CNV frequency >40% and GISTIC q-value <0.05. For methylation profile analysis, we first assigned DNA methylation values for each gene with the average beta value of the probes mapped to the promoter region, including TSS200 (region from –200 bp upstream to the transcription start site (TSS) itself), 1stExon (the first exon), TSS1500 (from –200 to –1500 bp upstream of TSS), and 5′UTR in order. For each phenotype, we identified the epigenetically silenced genes (ESGs) using the following criteria: (1) excluding the CpG sites methylated in normal tissues (mean β-value of □>□0.2) or less than 10% of the tumor samples; (2) the DNA methylation data was divided into the methylation group and unmethylation group, according to the cutoff (β-value□=□0.3); (3) for each gene, if the difference between the corresponding gene mean expression in the unmethylated group and that in the methylated group was□>□1.64 standard deviations of the unmethylated group, the gene would be labeled as epigenetically silenced(***Liu et al., 2021***).

#### Assessments of TMB, NAL, and CNV burden

TMB was defined as the total non-silent somatic mutation counts in coding regions, encompassing missense, nonsense, frame shift insertion, frame shift deletion, in-frame insertion, in-frame deletion, and splice site mutation. The “maftools” R package was utilized to process mutation data and calculate the TMB of each patient. Neoantigens of the TCGA-CRC dataset measured by TCGA official were available in TCIA database (https://tcia.at/neoantigens)(***Charoentong et al., 2017***). The fraction of genome alteration (FGA), fraction of genome gained (FGG), and fraction of genome lost (FGL), were defined as the ratio of total CNV/gain/lost bases to all bases, respectively. The TCGA-CRC cohort was calculated based on copy number segment data as follows:

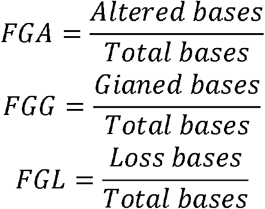

#### Estimating drug sensitivity in clinical cohort

To facilitate the subtype-based targeted interventions, we introduced an integrated pipeline to identify potential therapeutic agents for each subtype(***Yang et al., 2021***) (Figure S20A). Three pharmacogenomic datasets, CTRP, PRISM, and GDSC, store large-scale drug response and molecular data of human cancer cell lines, enabling accurate prediction of drug response in clinical samples(***Yang et al., 2021***). As mentioned above, stromal components could obscure the expression patterns of cancer cells in clinical samples. A purification algorithm termed *ISOpure*(***Quon et al., 2013***) was adopted to eliminate the contamination of stromal signal in the discovery cohort prior to conducting drug response prediction, and further yielded a purified tumor expression profiles comparable to cell lines(***Yang et al., 2021***). After purification, the proportion of stromal-rich subtypes (e.g., GINS2 and GINS4) were obviously decreased, suggesting the impact of stroma components had been eliminated (Figure S20B). A PDX dataset, GSE73255(***Isella et al., 2017***), is naturally uncontaminated by human stromal components. Hence, we tested our pipeline in the discovery cohort and GSE73255, and ultimately identified intersecting subtype-specific agents in two datasets. As previously reported(***Yang et al., 2021***), the model used for predicting drug response was ridge regression model implemented in the *pRRophetic* package(***Geeleher et al., 2014***). This predictive model was trained on mRNA expression profiles and drug response data of cancer cell lines with a satisfied predictive accuracy were evaluated by default 10-fold cross-validation, thus allowing the estimation of clinical drug response using only patients’ baseline gene expression data. We inputted the purified tumor expression profiles into the regression model to estimate the drug response of clinical samples, and ultimately determined the consensus agents in both the discovery cohort and GSE73255.

### Statistical analysis

All data processing, statistical analysis, and plotting were conducted in R 4.0.5 software. Correlations between two continuous variables were assessed via Pearson’s correlation coefficients. The chi-squared or fisher exact test was applied to compare categorical variables, and continuous variables were compared through the Wilcoxon rank-sum test or T test. The coefficient of variation (CV) is the ratio of the standard deviation to the mean and shows the extent of variability in relation to the mean of the population. The higher the CV, the greater the dispersion. R was used to measure the fitting level of the power law curve, The better the curve fitting level is, the closer R is to −1. Kaplan-Meier analyses were performed via the survival package. All statistical tests were two-sided. *P* <0.05 was regarded as statistically significant.

## Supplementary Figures

**Figure 1-figure supplement 1.**
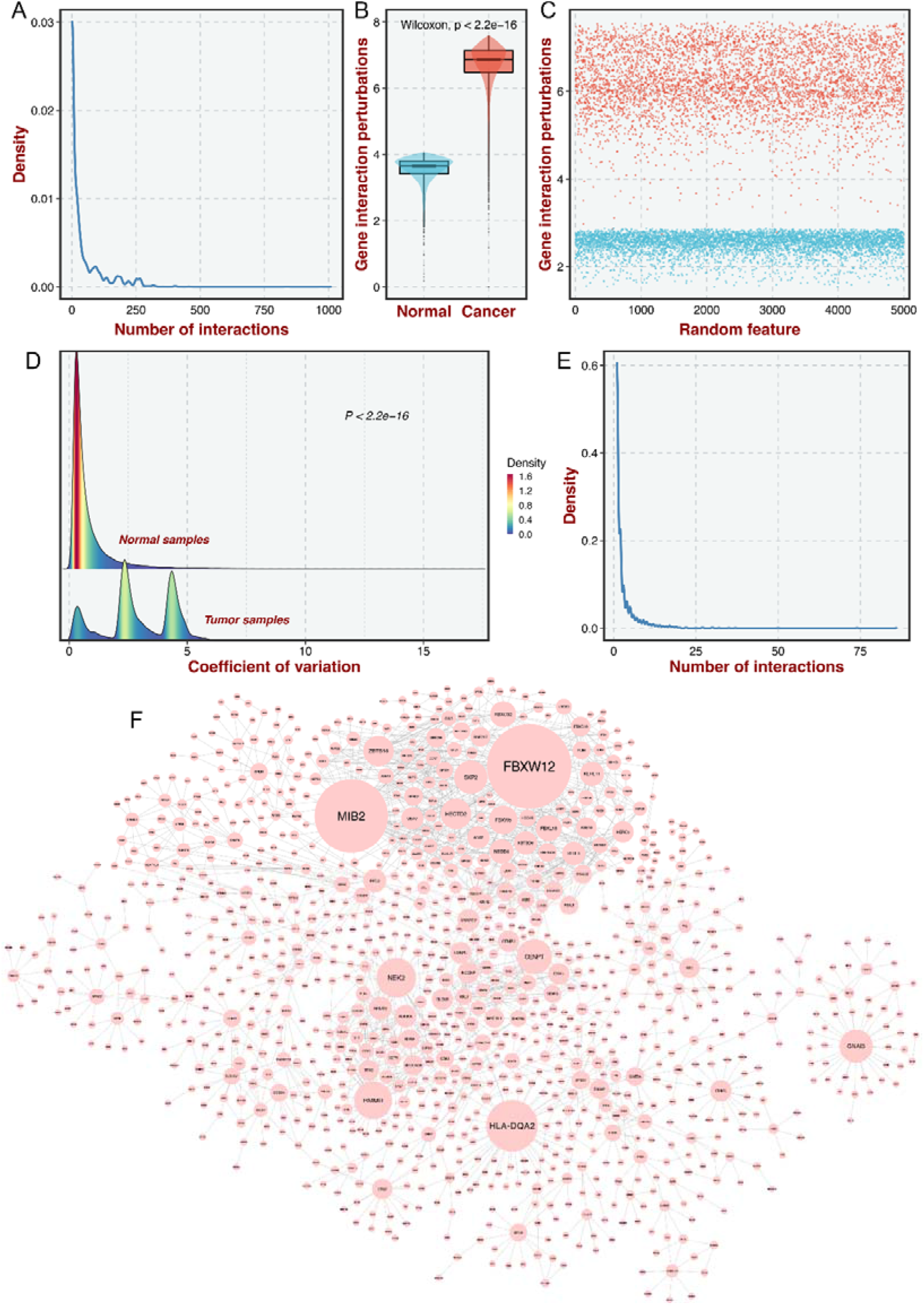
Construction of the gene interaction-perturbation network. **A.** As the number of interactions increased, the density decreased significantly, presenting a power distribution in the background networks. *R* was computed as the Pearson correlation between log_10_ (interaction number) and log_10_ (corresponding frequency), which was used to measure the fitting level of the power law curve. The better the curve fitting level is, the closer *R* is to 1. **B.** The distribution of gene interaction perturbations between normal and tumor samples. **C.** The scatterplot for the log2-transformed mean of the interaction perturbations in the 5000 randomly selected edges in both normal (blue points) and CRC (red points) tissues. The interaction perturbations of normal samples were much denser and less than tumor samples. **D.** 92.6% of all 148,942 gene pairs exhibited more dispersion in tumor samples than in normal samples by comparing the coefficient of variation (CV) of interaction perturbations. **E-F.** This new network with 1,390 genes and 2,225 interactions also met the scale-free distribution (**E**) and was visualized (**F**), the node size represents connectivity.

**Figure 2-figure supplement 1.**
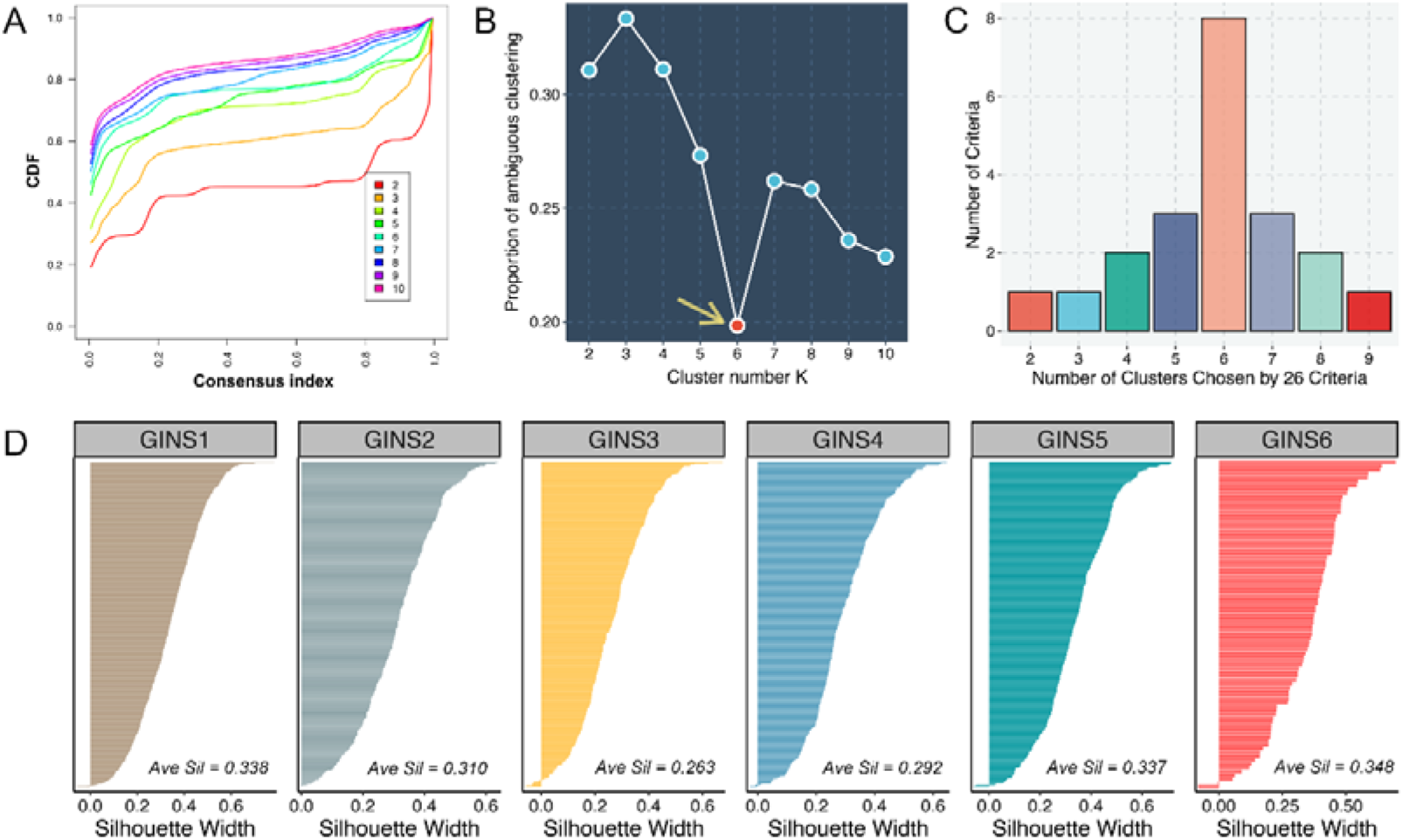
Six CRC subtypes were identified from the gene interaction-perturbation network. **A.** The CDF curves of consensus matrix for each *K* (indicated by colors). **B.** The proportion of ambiguous clustering (PAC) score, a low value of PAC implies a flat middle segment, allowing conjecture of the optimal *K* (*K* = 6) by the lowest PAC. **C.** Recommended number of clusters using 26 criteria of Nbclust package. **D.** Silhouette width of samples in each subtype.

**Figure 2-figure supplement 2.**
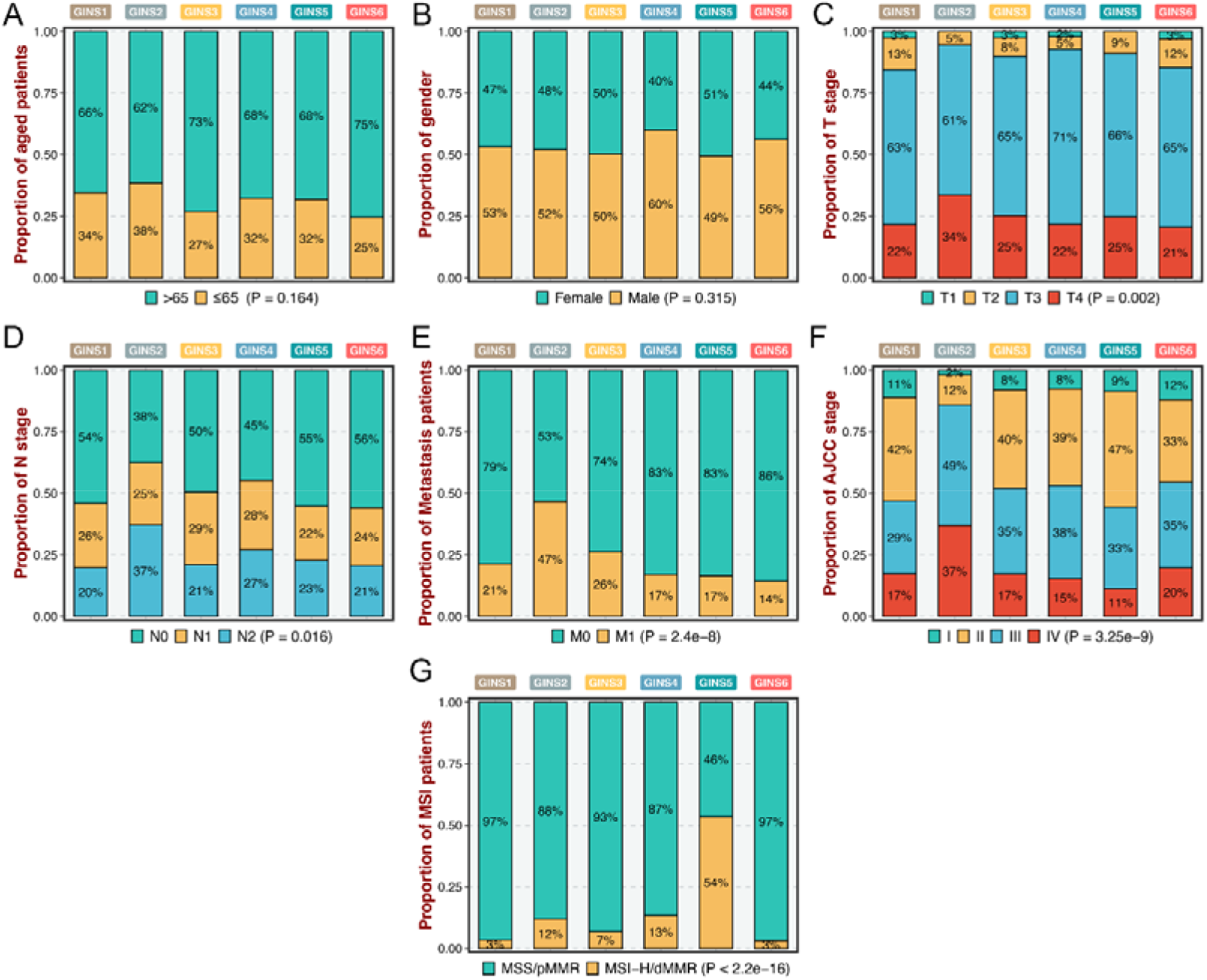
Clinical characteristics of six GINS subtypes. **A-G.** Barplots showed the distribution of age (**A**), gender (**B**), T stage (**C**), N stage (**D**), M stage (**E**), AJCC stage (**F**), and MSI status (**G**).

**Figure 2-figure supplement 3.**
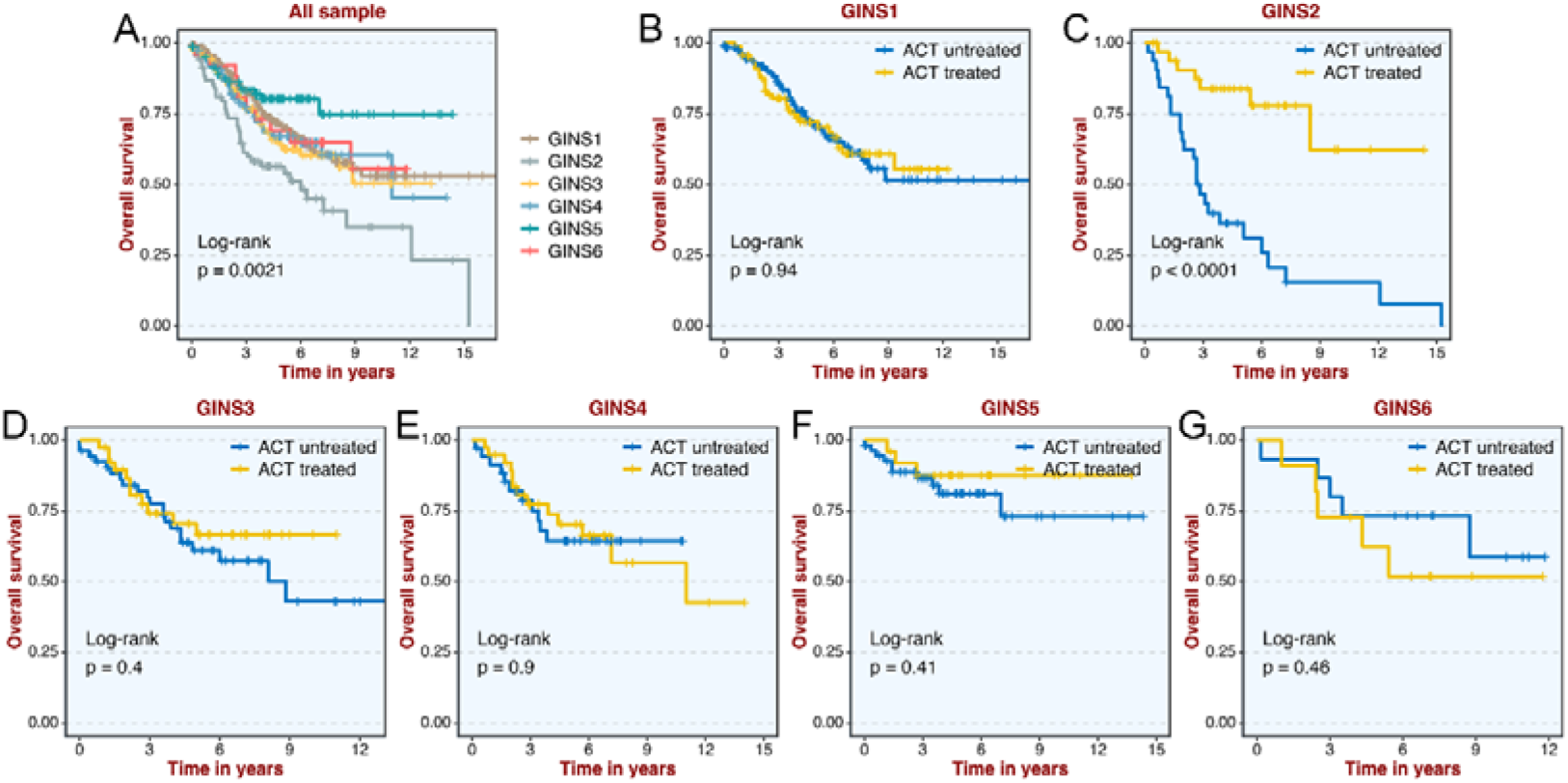
Association of GINS subtypes with ACT after surgery for 585 patients in GSE39582 dataset. **A-G** Kaplan-Meier curves of overall survival for six GINS subtypes across all (**A**), GINS1 (**B**), GINS2 (**C**), GINS3 (**D**), GINS4 (**E**), GINS5 (**F**), GINS6 (**G**) samples, respectively.

**Figure 2-figure supplement 4.**
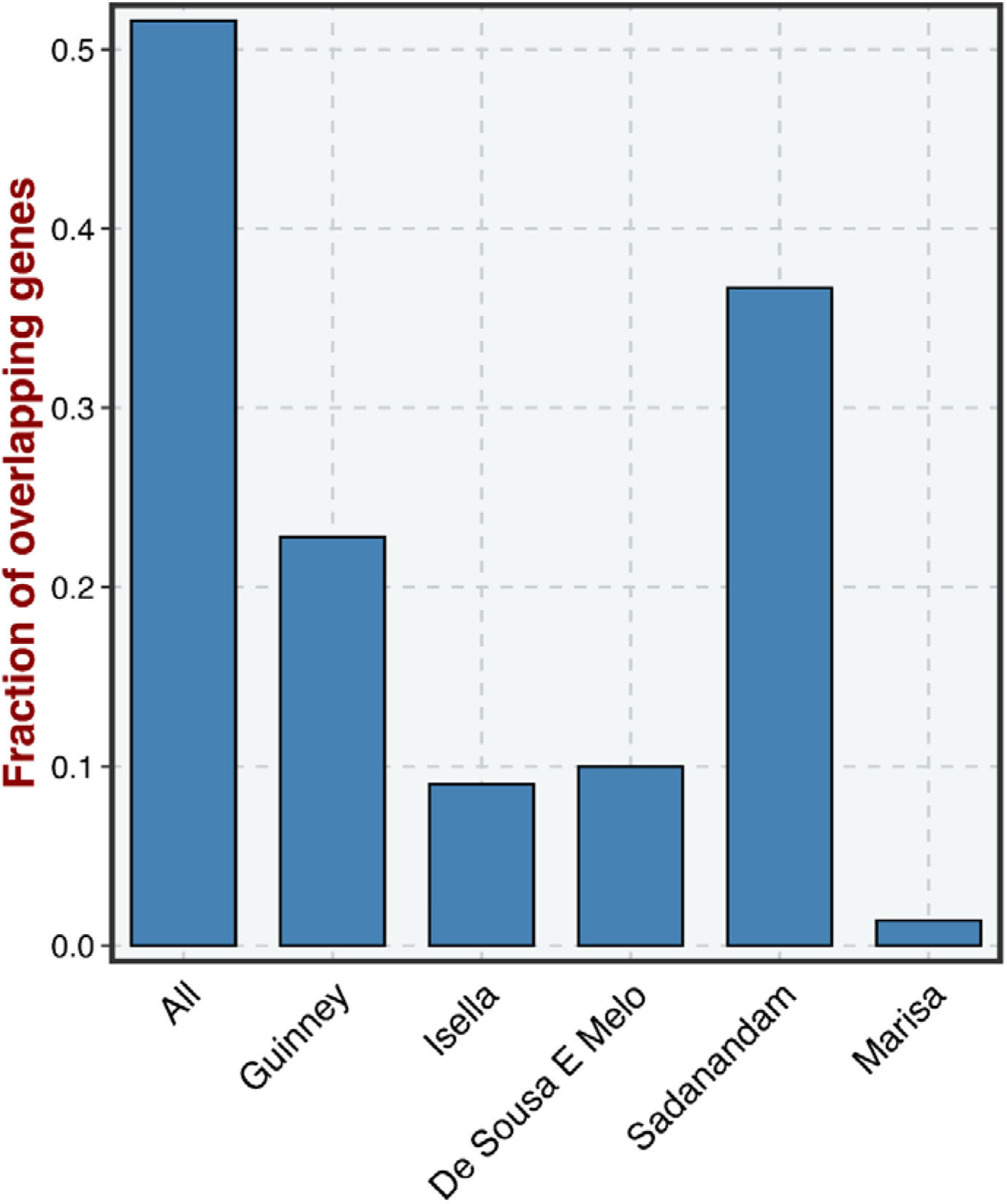
The overlapped genes between our classifier genes and the signature genes of all previous CRC classifications.

**Figure 3-figure supplement 1.**
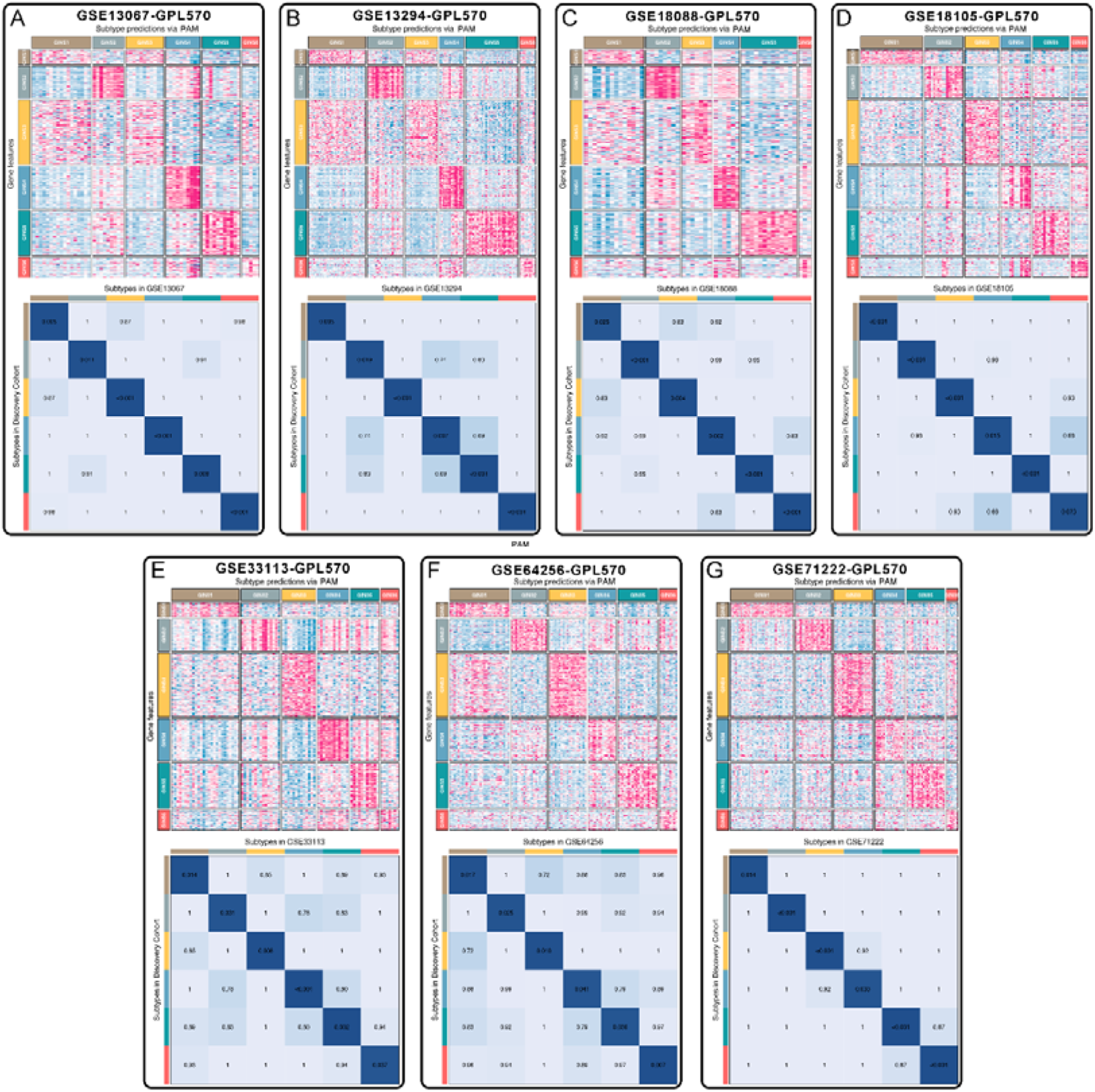
Subtype validation of seven datasets from the same platform (GPL570). **A-G.** Seven datasets, including GSE13067 (**A**), GSE13294 (**B**), GSE18088 (**C**), GSE18105 (**D**), GSE33113 (**E**), GSE64256 (**F**), and GSE71222 (**G**), were assigned in six subtypes according to the classifier. The top and left bars indicated the subtypes. In the heatmap, rows indicated genes from the classifier and columns represent patients. The heatmap was color-coded on the basis of median-centered log_2_ gene expression levels (red, high expression; blue, low expression). SubMap plots, located in the bottom panel, assessed expressive similarity between corresponding subtypes from two different cohorts.

**Figure 3-figure supplement 2.**
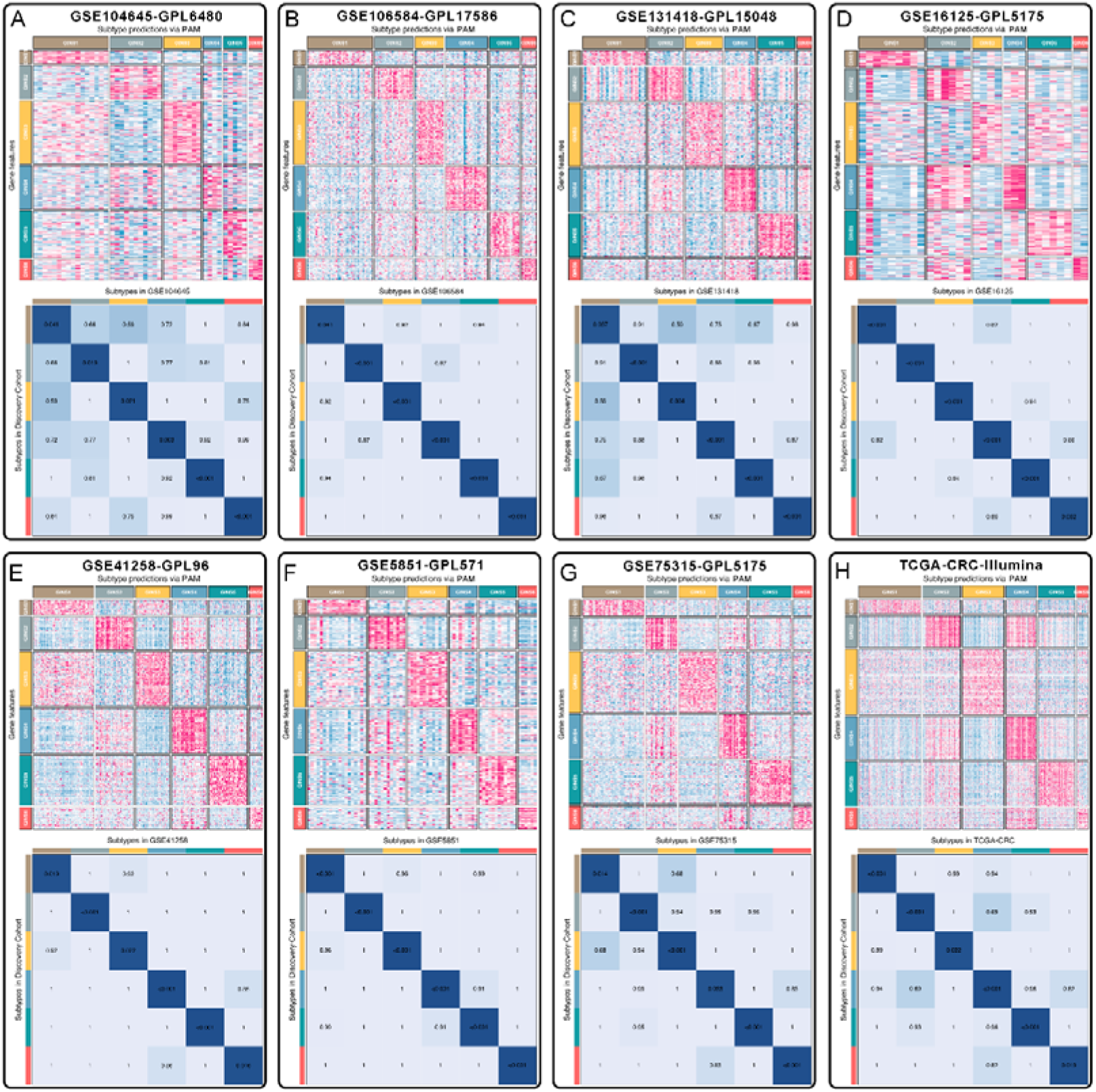
Subtype validation of seven microarrays from different platforms and a RNA-seq dataset. **A-H.** Eight datasets, including GSE104645 (**A**), GSE106584 (**B**), GSE131418 (**C**), GSE16125 (**D**), GSE41258 (**E**), GSE5851 (**F**), GSE75315 (**G**), and TCGA-CRC (**H**), were assigned in six subtypes according to the classifier. The top and left bars indicated the subtypes. In the heatmap, rows indicated genes from the classifier and columns represent patients. The heatmap was color-coded on the basis of median-centered log_2_ gene expression levels (red, high expression; blue, low expression). SubMap plots, located in the bottom panel, assessed expressive similarity between corresponding subtypes from two different cohorts.

**Figure 3-figure supplement 3.**
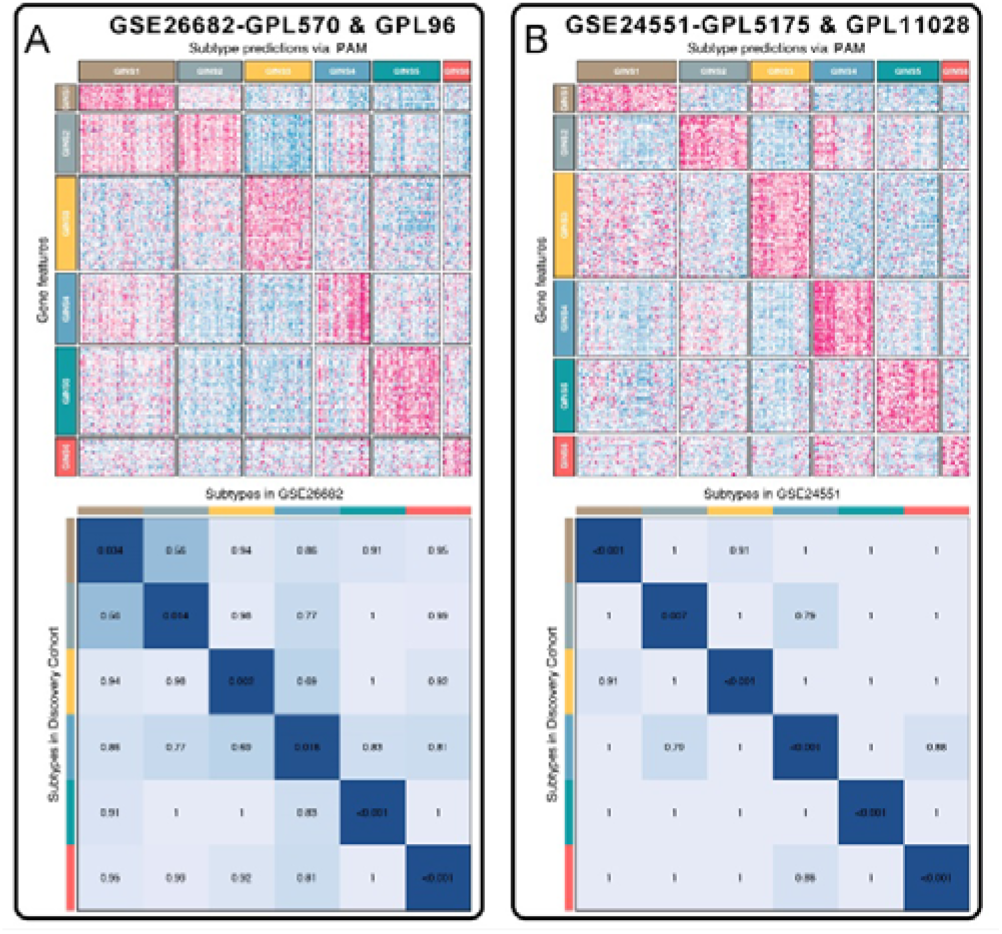
Subtype validation of two microarrays chip with two different platforms. **A-B.** Two datasets, GSE26682 and GSE24551, each chip from two different platforms (GPL570 & GPL96 for GSE26682 and GPL5175 & GPL11028 for GSE24551), were assigned in six subtypes according to the classifier. The top and left bars indicated the subtypes. In the heatmap, rows indicated genes from the classifier and columns represent patients. The heatmap was color-coded on the basis of median-centered log_2_ gene expression levels (red, high expression; blue, low expression). SubMap plots, located in the bottom panel, assessed expressive similarity between corresponding subtypes from two different cohorts.

**Figure 3-figure supplement 4.**
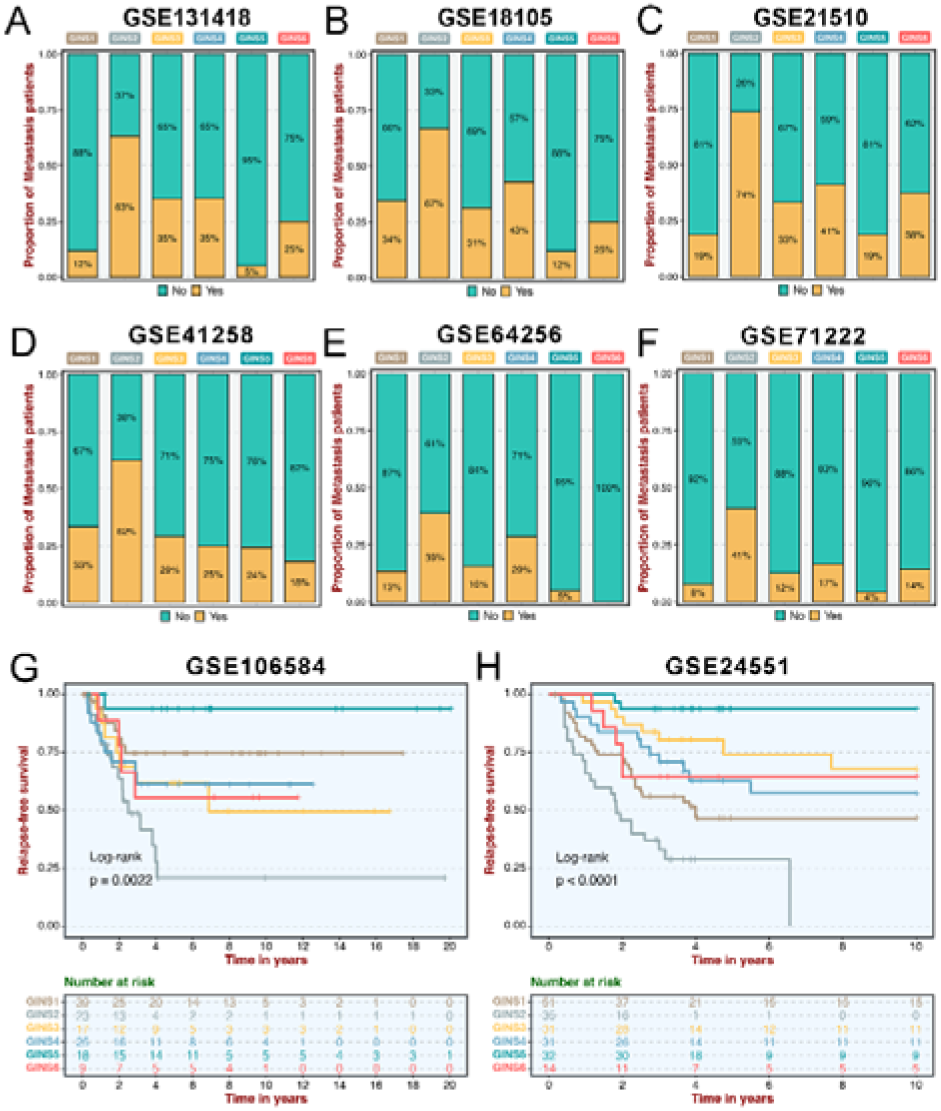
Clinical characteristics of six GINS subtypes in validation datasets. **A-F.** Barplots showed the distribution of metastasis patients to six subtypes in GSE131418 (**A**), GSE18105 (**B**), GSE21510 (**C**), GSE41258 (**D**), GSE64256 (**E**), and GSE71222 (**F**). **G-H.** Kaplan-Meier curves of overall survival for six GINS subtypes in GSE106584 (**G**) and GSE24551 (**H**).

**Figure 4-figure supplement 1.**
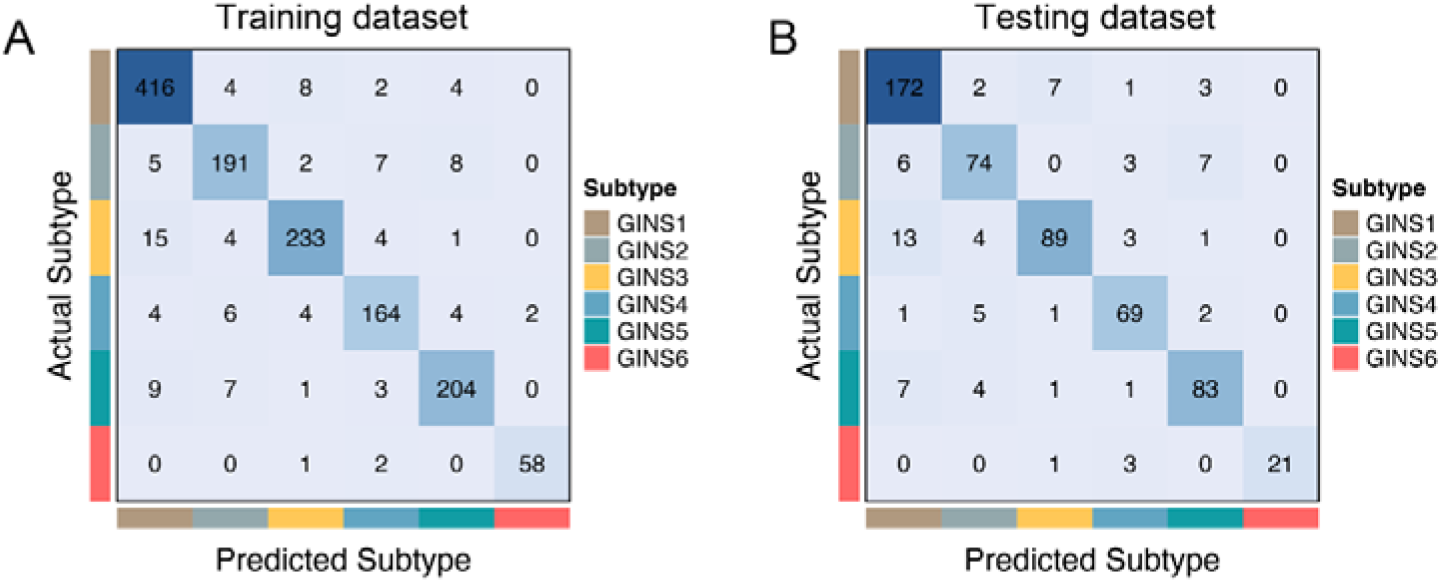
Performance of the miniclassifier. **A-B.** Confusion matrix displayed the general tendency of classification effect in the training (**A**) and testing (**B**) datasets, respectively.

**Figure 5-figure supplement 1.**
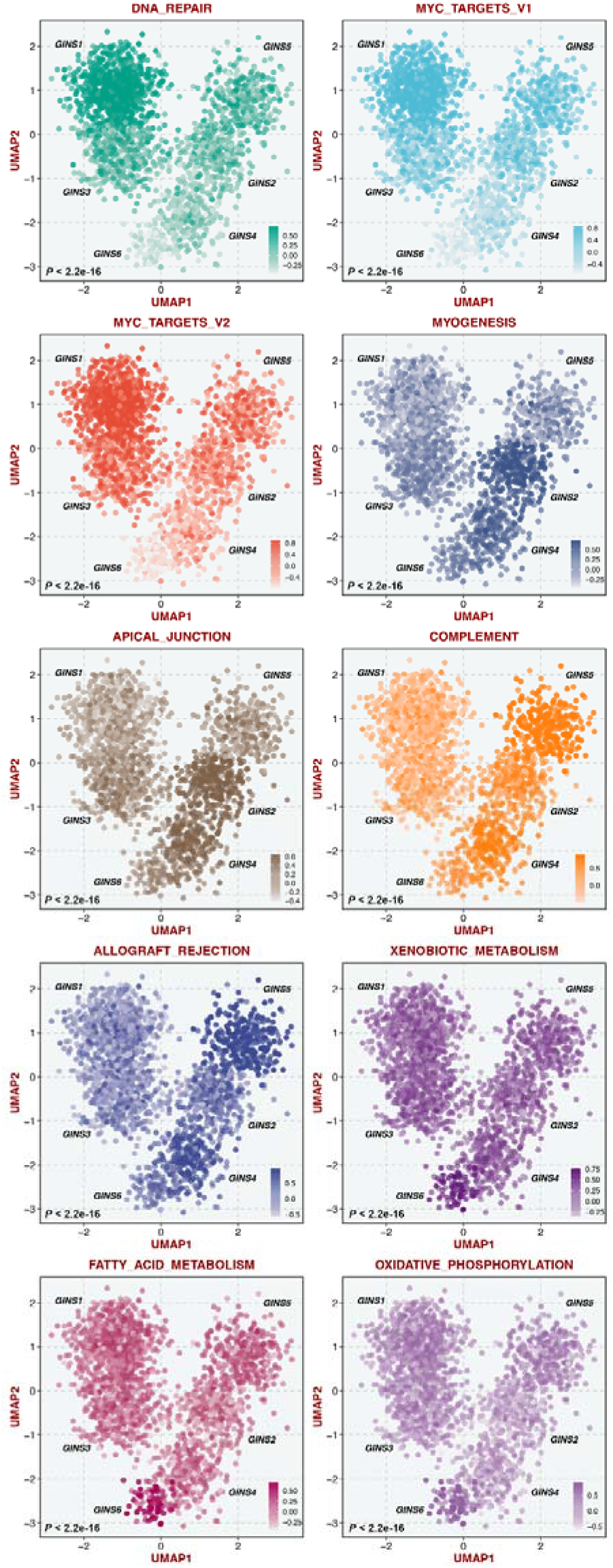
GSVA estimated differences in pathway activity across six subtypes.

**Figure 5-figure supplement 2.**
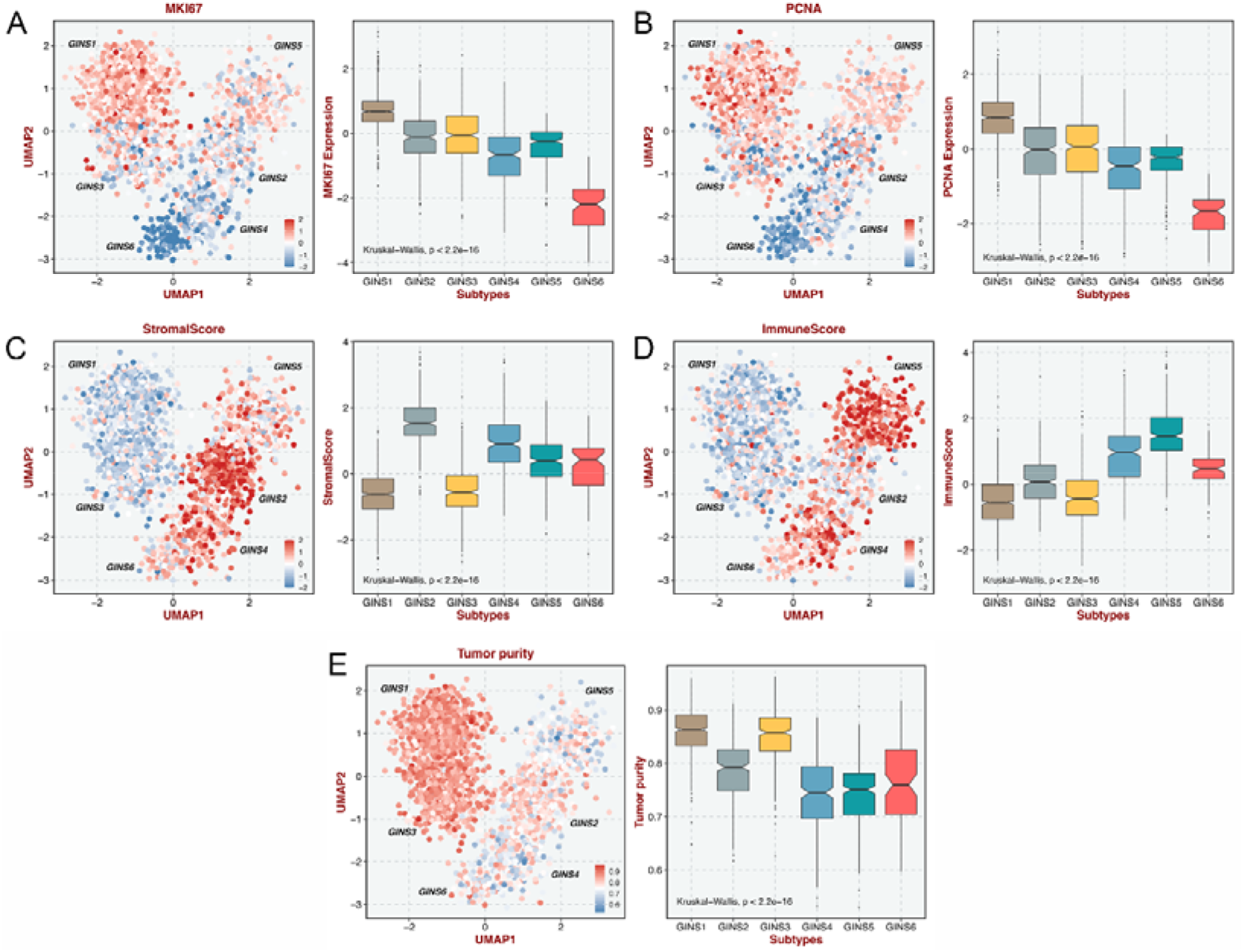
The distribution of *MKI67* (**A**), *PCNA* (**B**), stromal score (**C**), immune score (**D**), and tumor purity (**E**) in six subtypes.

**Figure 6-figure supplement 1.**
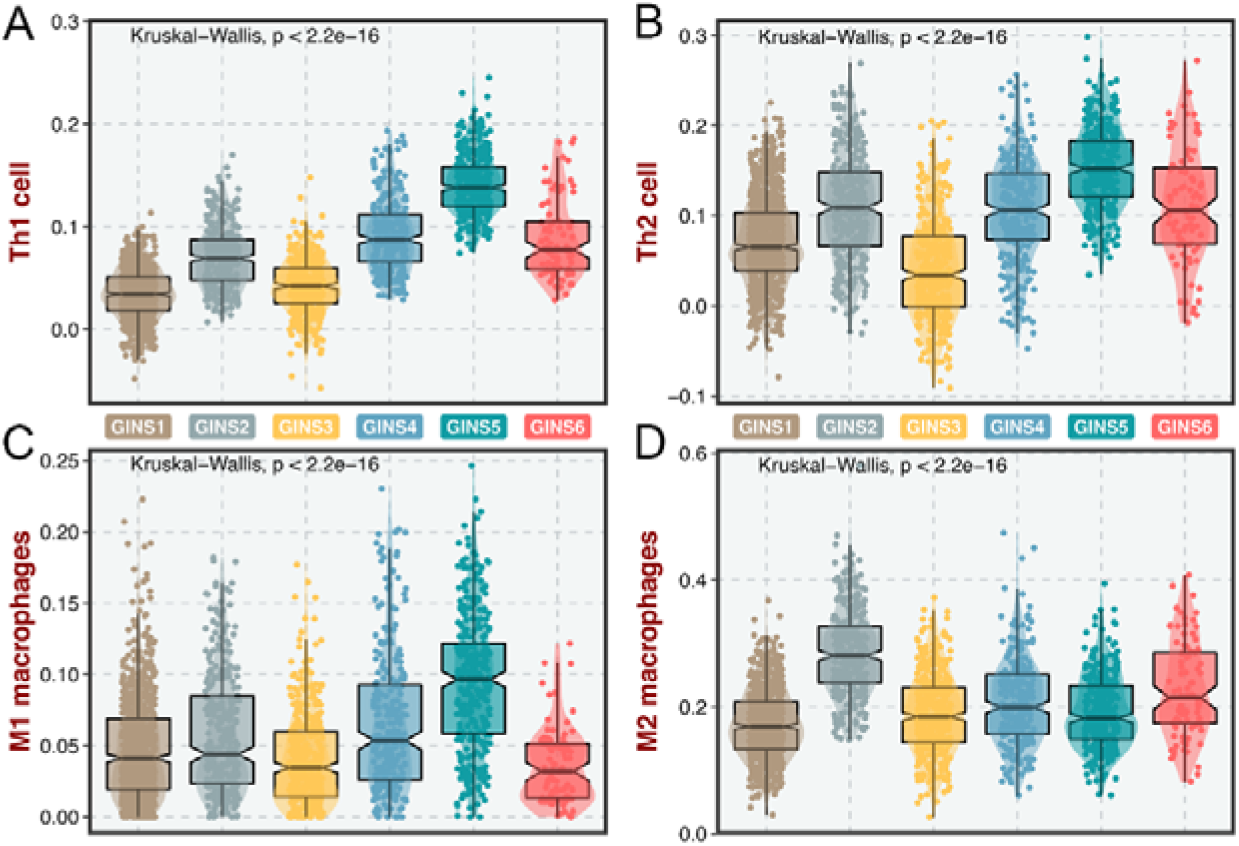
Immune cell infiltrations of six subtypes. **A-D.** The infiltration distribution of Th1 (**A**), Th2 (**B**), M1 (**C**), and M2 (**D**) cells.

**Figure 7-figure supplement 1.**
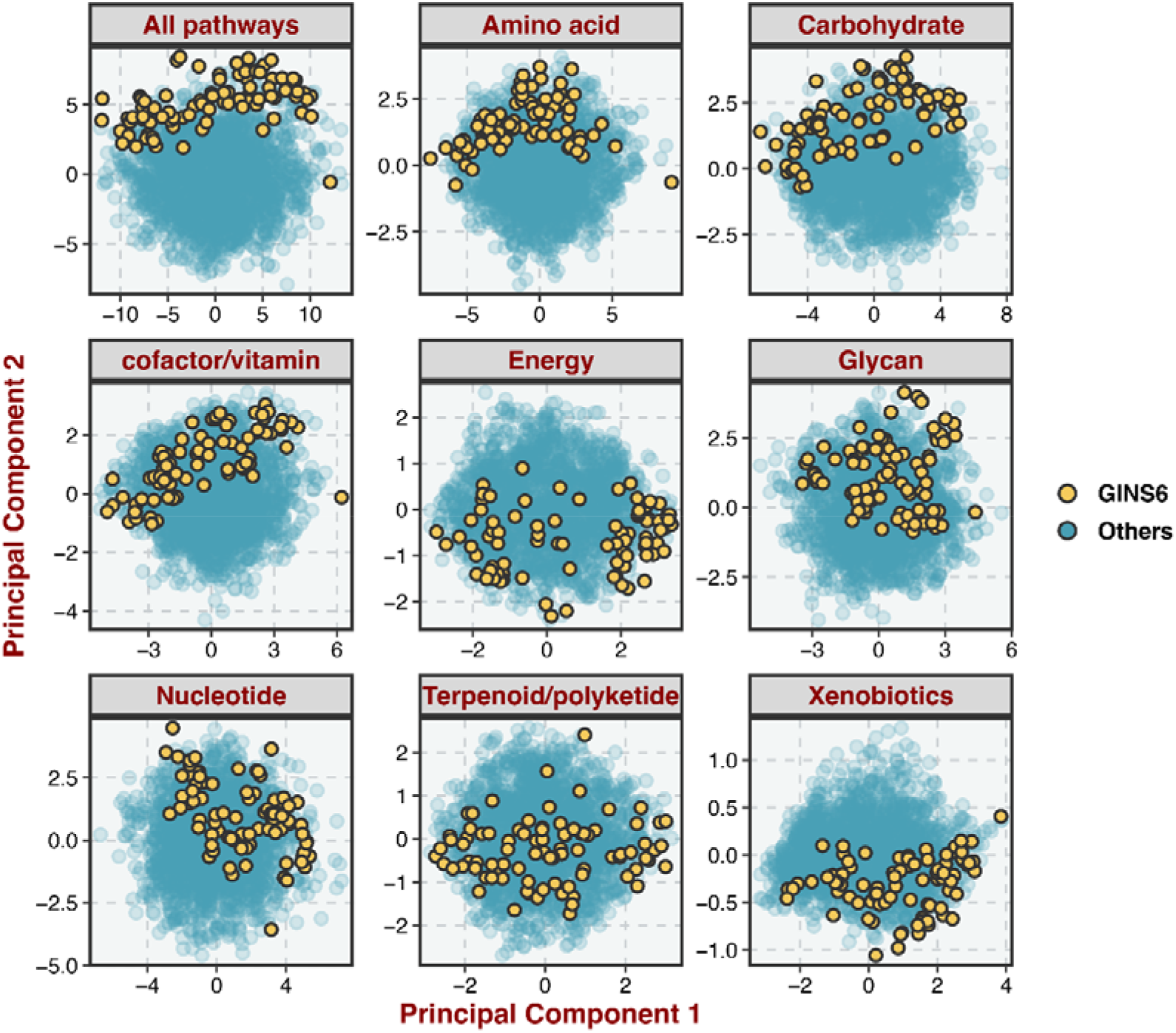
Principal component analysis of all samples in the discovery cohort for the mRNA expression of metabolic genes.

**Figure 8-figure supplement 1.**
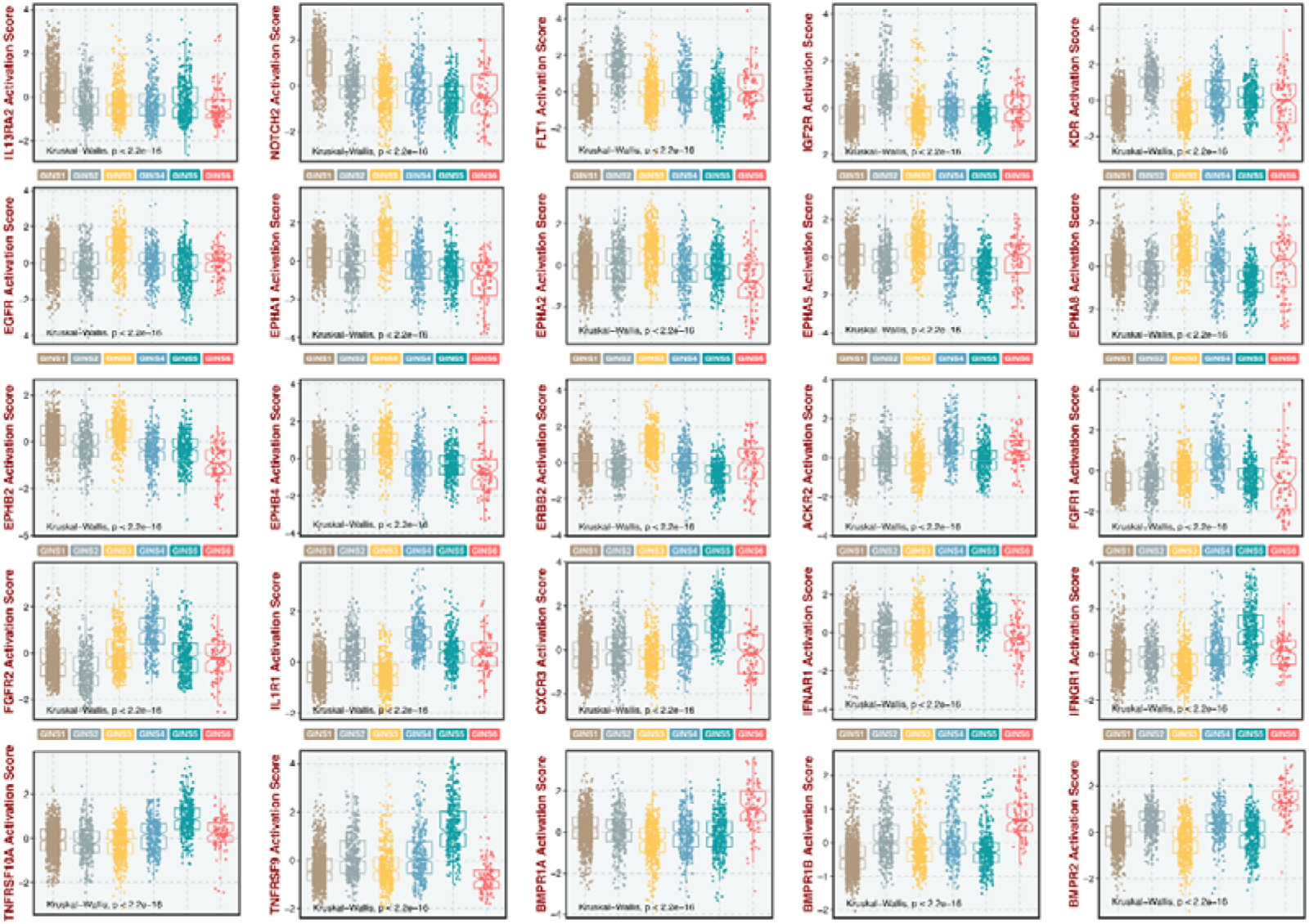
GSVA further estimated differences in Receptor-Ligand activities across six subtypes.

**Figure 9-figure supplement 1.**
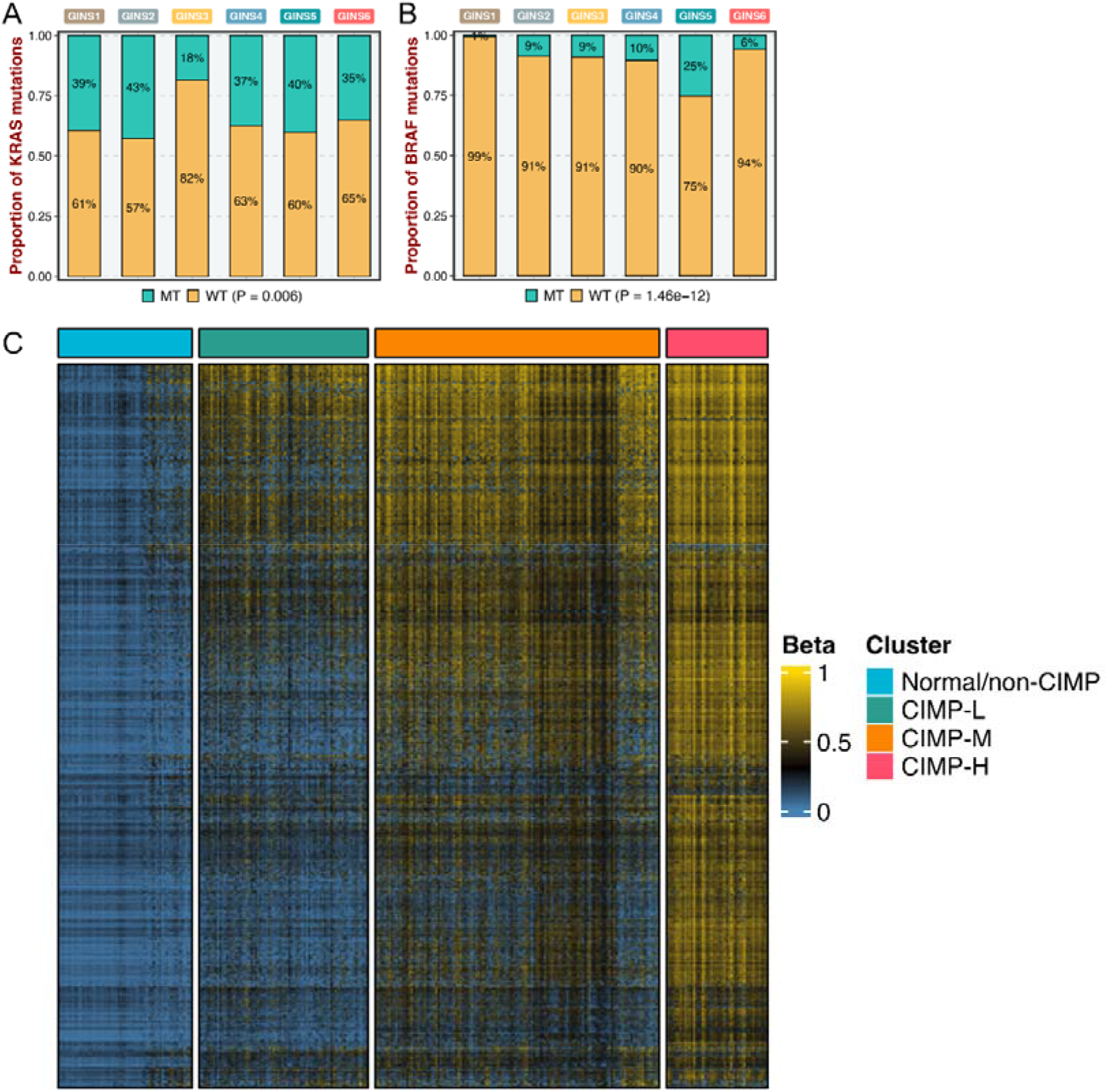
Genomic alterations of six subtypes. **A-B.** Barplots showed the distribution of *KRAS* (**A**) and *BRAF* (**B**) mutations to six subtypes in the discovery cohort. **C.** Four CpG island methylator phenotype (CIMP) were identified from the TCGA-CRC cohort using the beta value of 5000 CpG island promoters with the most variation.

**Figure 10-figure supplement 1.**
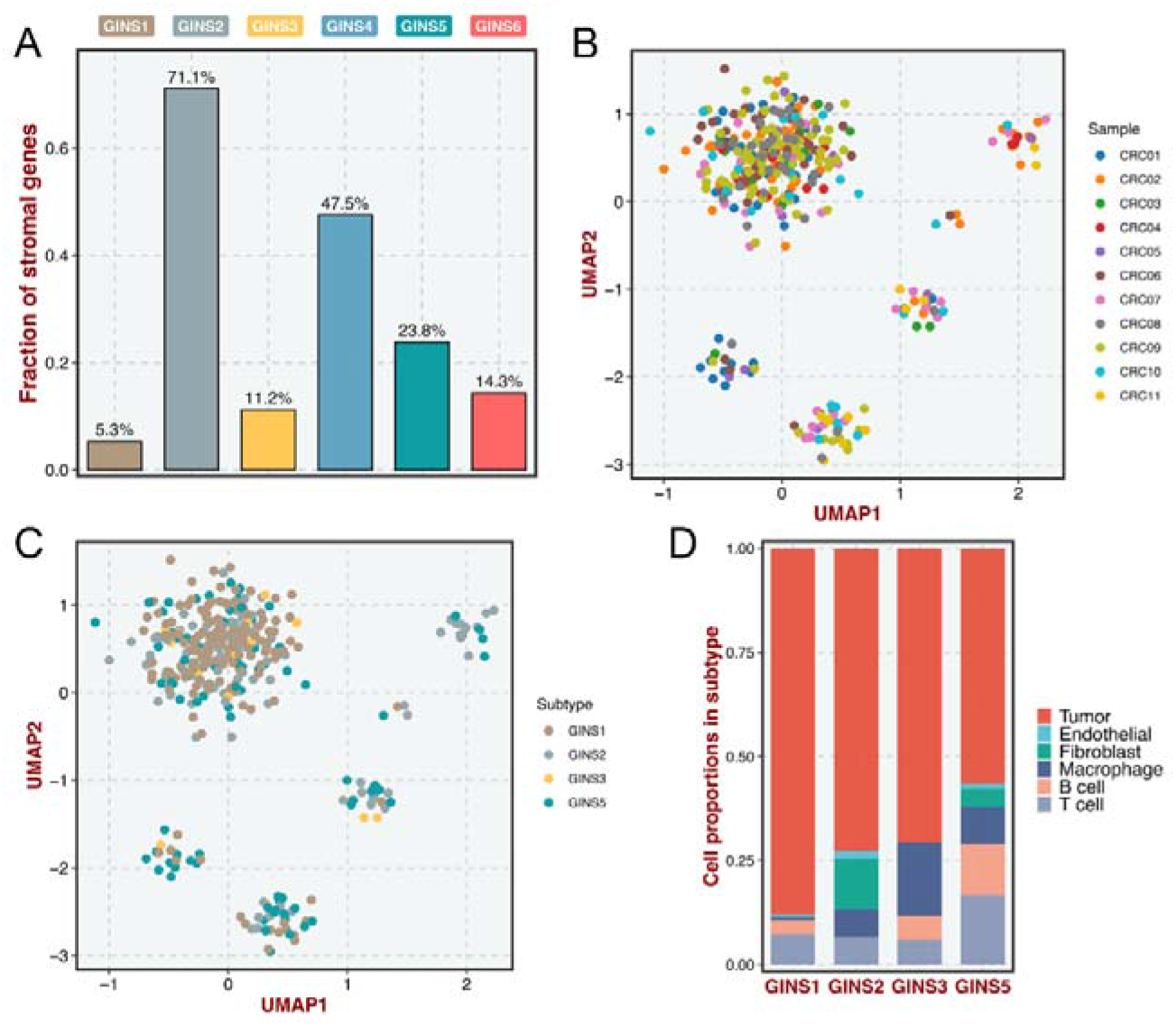
Stromal contribution to the subtype transitions. **A.** Fractions of subtype-discriminant genes of the PAM classifier belonged to stromal genes. **B-C.** The cell distribution of 11 patients (**B**) and GINS subtypes (**C**) according to the two-dimension UMAP analysis. **D.** The composition of cells in each subtype.

**Figure 10-figure supplement 2.**
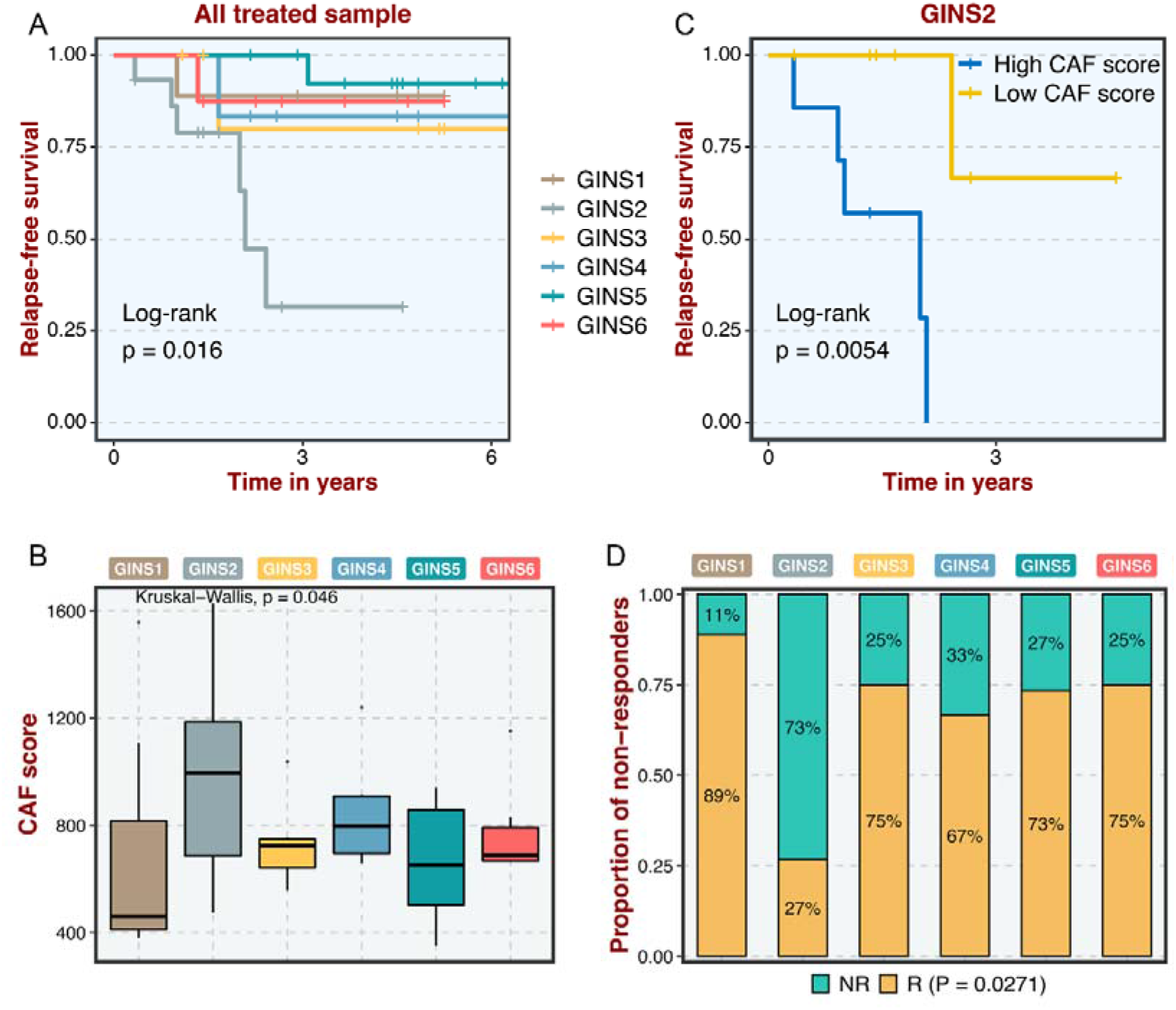
Clinical significance of six subtypes in GSE56699. **A.** Kaplan-Meier curves of overall survival for six GINS subtypes across all treated samples. **B.** The distribution of CAF score in six subtypes. **C.** Kaplan-Meier curves of overall survival for high and low CAF groups across GINS2 samples. **D.** The distribution of radiotherapeutic non-responders in six subtypes.

**Figure 10-figure supplement 3.**
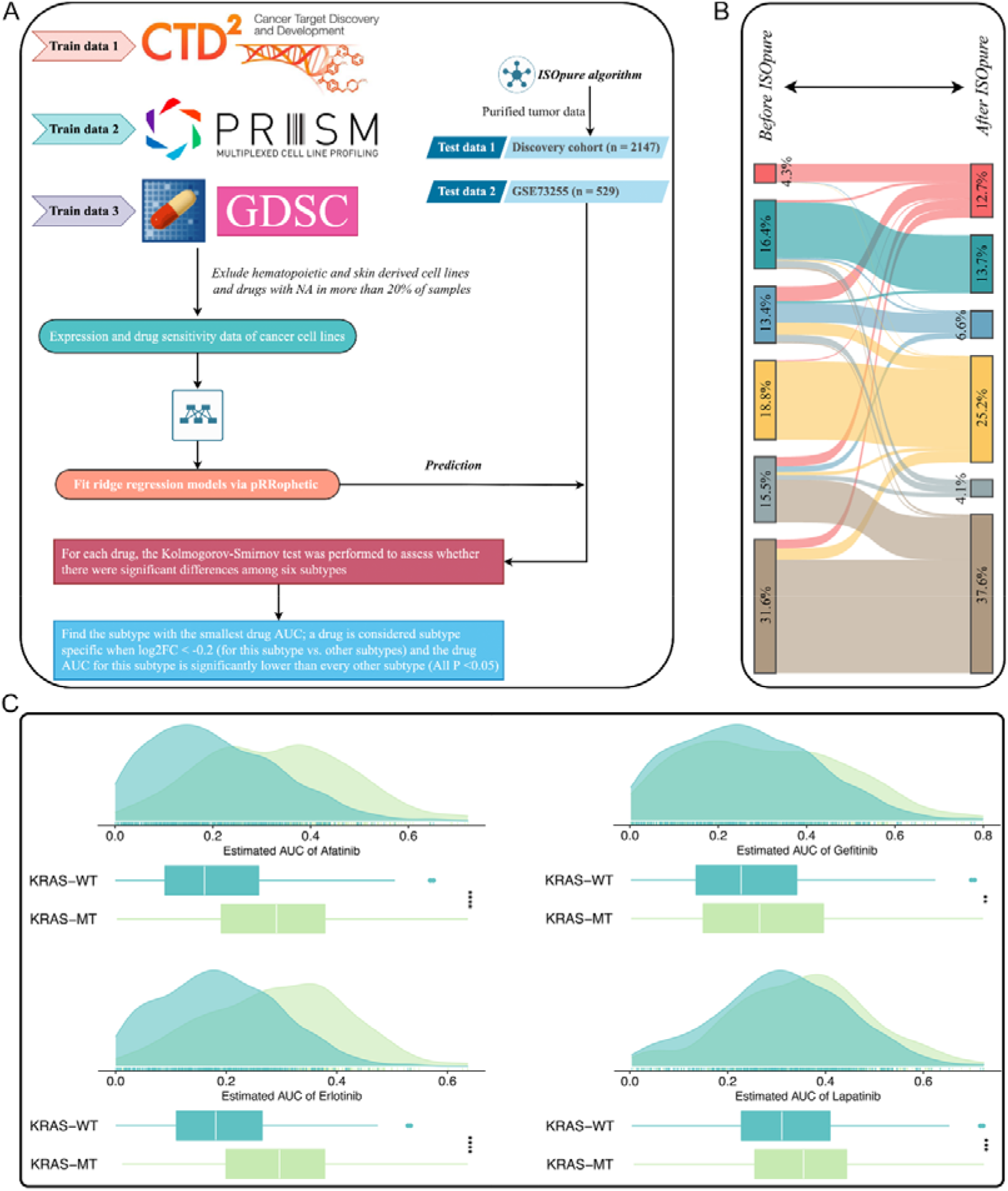
Identification of potential therapeutic agents for six subtypes. **A.** Overview of the analysis procedures. **B.** Transcriptional classification of paired samples before and after purification. **C.** Comparison of estimated drug sensitivity (logAUC) between *KRAS* mutant and wild type group. \*\**P* <0.01, \*\*\**P* <0.001, \*\*\*\**P* <0.0001.

**Figure 10-figure supplement 4.**
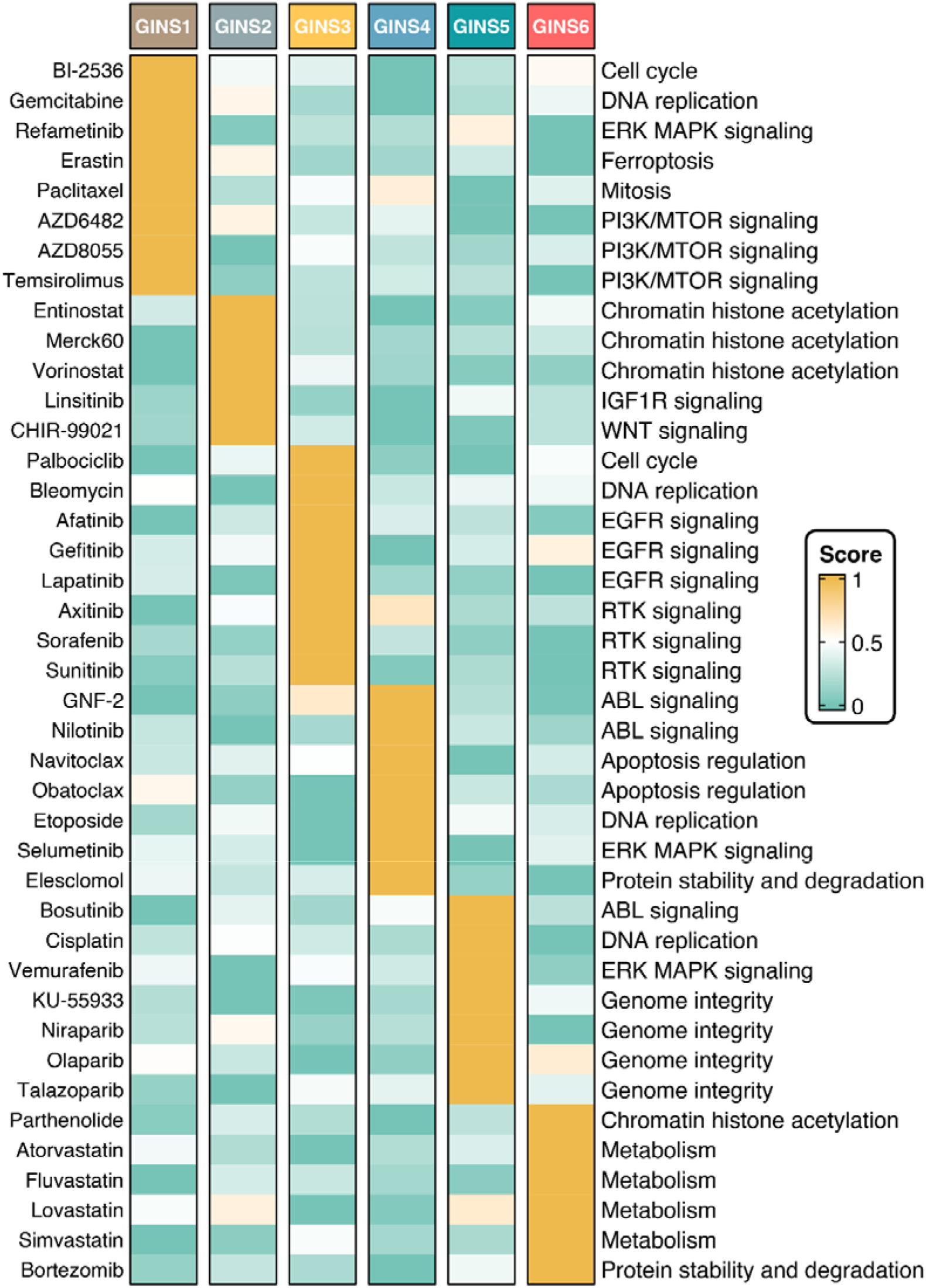
Candidate agents of six subtypes. Score represents the normalized drug sensitivity (logAUC).

## Supplementary Tables

**Supplementary File 1.** The PAM-centroid distance classifier included 289 subtype-discriminant genes.

**Supplementary File 2.** Clinical characteristics of six subtypes in validation datasets.

**Supplementary File 3.** Details of baseline information in our in-house dataset.

**Supplementary File 4.** The forward and reverse primers for qRT-PCR.

**Supplementary File 5.** Clinical characteristics of six subtypes in our in-house dataset.

**Supplementary File 6.** Publicly available gene signatures used in this study.

**Supplementary File 7.** Sample set enrichment analysis delineated the biological attributes inherent to GINS subtypes. \**P* <0.05, \*\**P* <0.01, \*\*\**P* <0.001.

**Supplementary File 8.** Gene cluster 3 and 10 were identified by Mfuzz analysis.

**Supplementary File 9.** The expression differences of 145 immunomodulators among six subtypes.

**Supplementary File 10.** The proteome (Reverse Phase Protein Array) data available from the TCGA portal included only 26 immunomodulators.

**Supplementary File 11**. GSEA results of 69 metabolic pathways from the Kyoto Encyclopedia of Genes and Genomes (KEGG) database.

**Supplementary File 12.** Metabolite-protein interaction network (MPIN) was established via nine GINS6-specific genes with broad and tight connections with lipid metabolites.

**Supplementary File 13.** Sample set enrichment analysis of ligand/receptor pairs expression in GINS subtypes. *NES and FDR are reported. NE, not enriched*.

**Supplementary File 14.** The targeted pahtways and molecules of candidate durgs for six subtypes.

**Supplementary File 15.** Details of data sources.

## Data availability

Public data used in this study are available in GEO, GTEx, TCGA, IMvigor210CoreBiologies, CCLE, GDSC, CTRP and PRISM databases. Essential scripts to develop the GINS taxonomy have been uploaded into Github (https://github.com/Zaoqu-Liu/GINS). Sequencing data available from GEO under accession codes GSE14333, GSE143985, GSE161158, GSE17537, GSE29621, GSE31595, GSE38832, GSE39084, GSE39582, GSE92921, GSE72970, GSE28702, GSE45404, GSE52735, GSE62080, GSE69657, GSE19860, GSE19862, GSE13067, GSE13294, GSE18088, GSE18105, GSE33113, GSE64256, GSE71222, GSE104645, GSE106584, GSE131418, GSE16125, GSE41258, GSE5851, GSE75315, GSE26682, GSE24551, GSE12945, GSE21510, GSE78220, GSE176307, GSE35144, GSE73255, GSE76402, GSE56699, and GSE81861. Normal tissue data is available from GTEx database (https://gtexportal.org). The TCGA-CRC multi-omics data, including RNA-seq (raw count), proteome (Reverse Phase Protein Array), HumanMethylation450 array, whole-exome sequencing (VarScan MAF files), and copy number variation (CNV) data, were derived from TCGA portal (https://portal.gdc.cancer.gov). Three datasets (n =414) with immunotherapeutic annotations and expression profiles were derived from the following studies: Hugo and colleagues(***Hugo et al., 2016***) (GSE78220, n =27), Rose and colleagues(***Rose et al., 2021***) (GSE176307, n =89) and Mariathasan and colleagues(***Mariathasan et al., 2018***) (IMvigor210, n =298). We retrieved 55 CRC cell lines with both transcriptome and metabolomics data (including 225 metabolites) from The Cancer Cell Line Encyclopedia (CCLE, https://sites.broadinstitute.org/ccle). Drug response and molecular data of human cancer cell lines were available from the Cancer Therapeutics Response Portal (CTRP, https://portals.broadinstitute.org/ctrp), Profiling Relative Inhibition Simultaneously in Mixtures (PRISM, https://depmap.org/portal/prism), and Genomics of Drug Sensitivity in Cancer (GDSC, https://www.cancerrxgene.org) datasets.

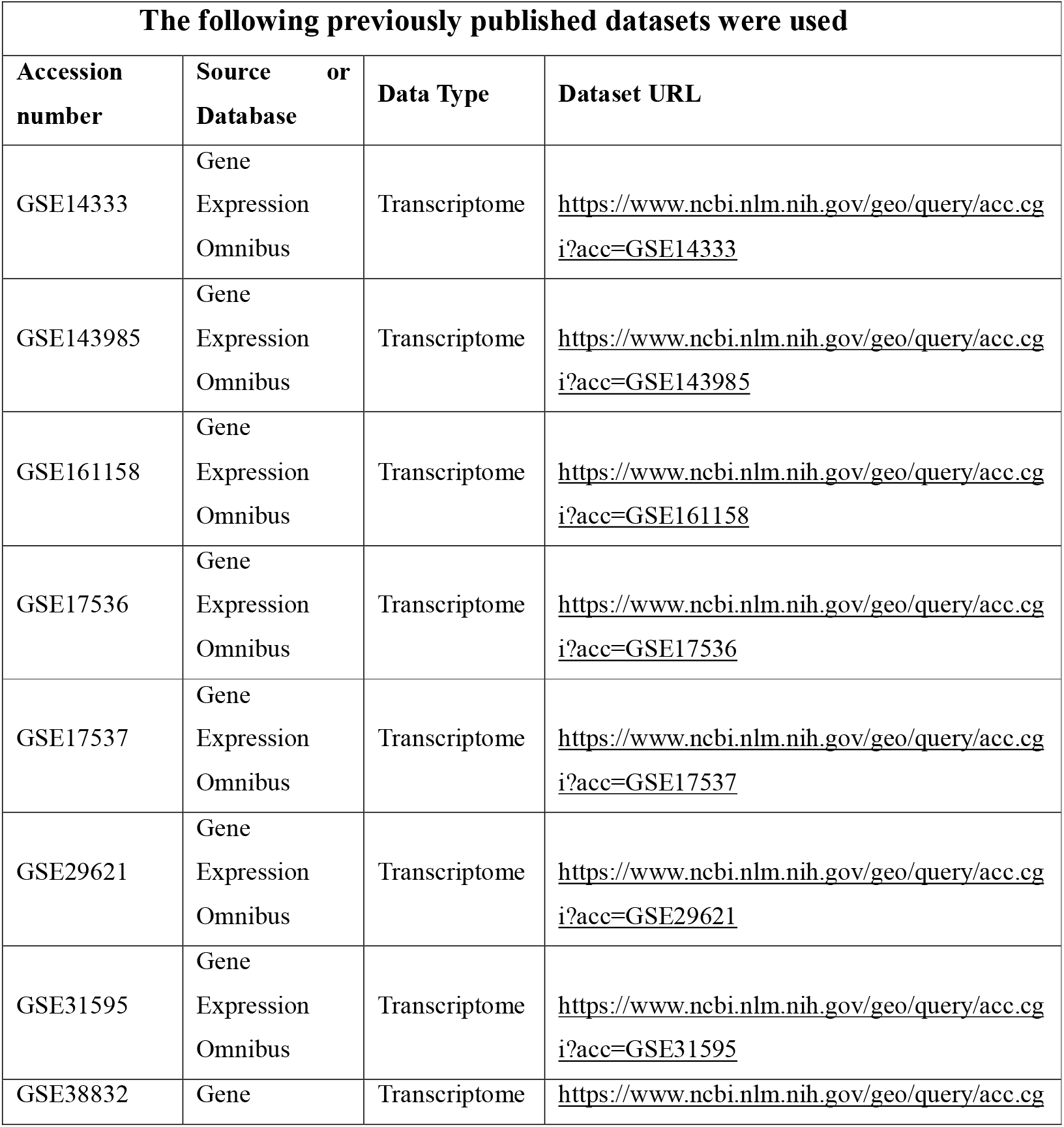
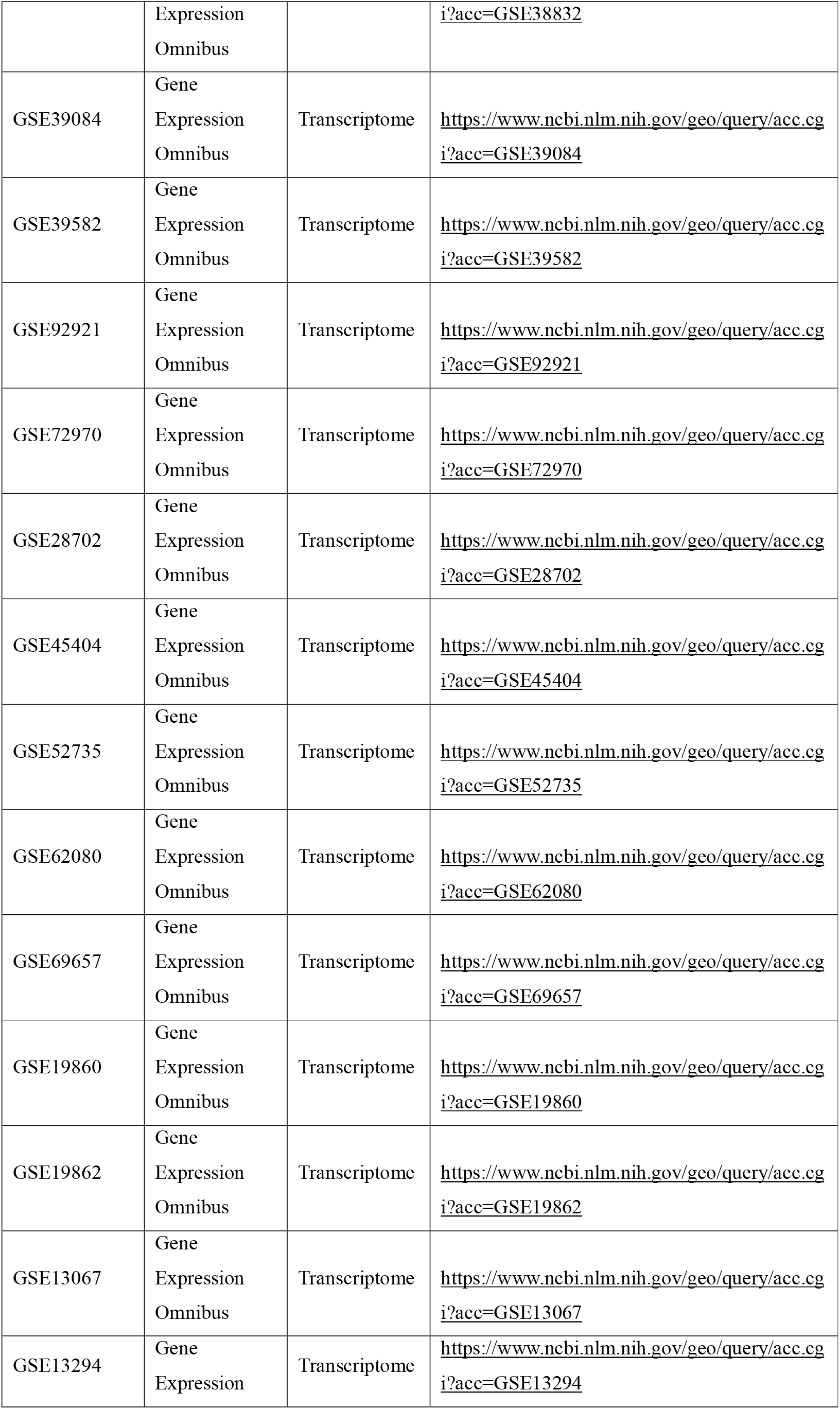
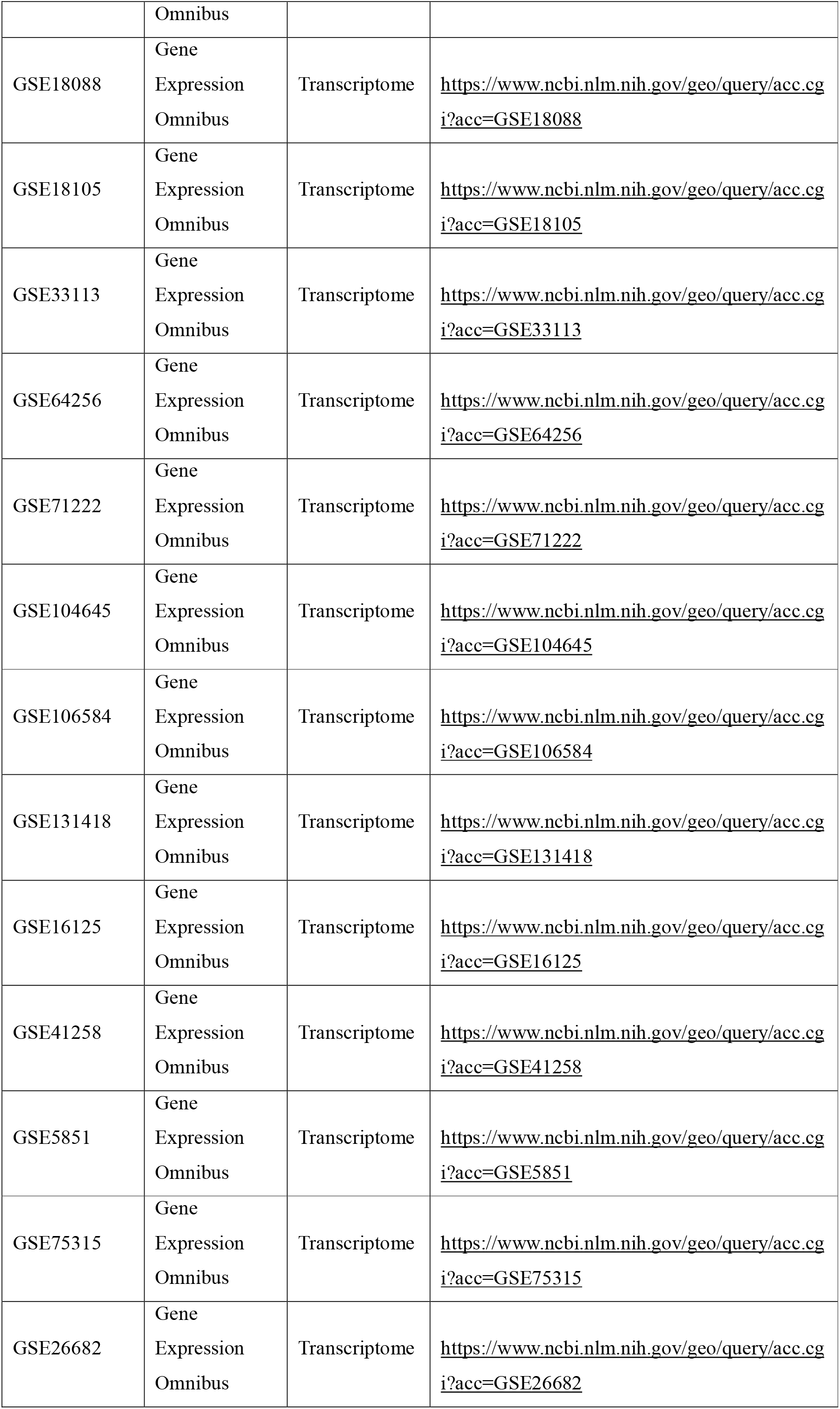
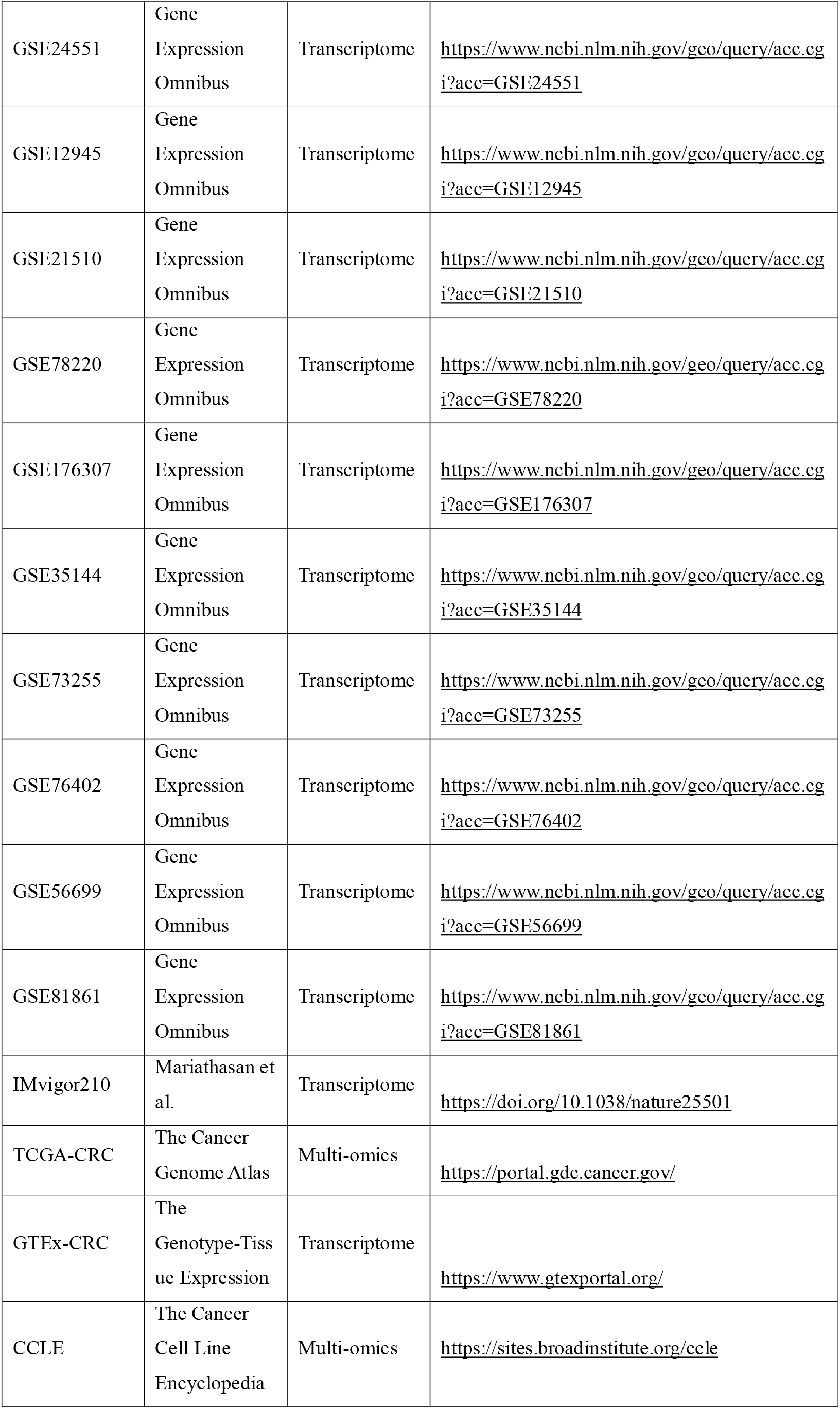
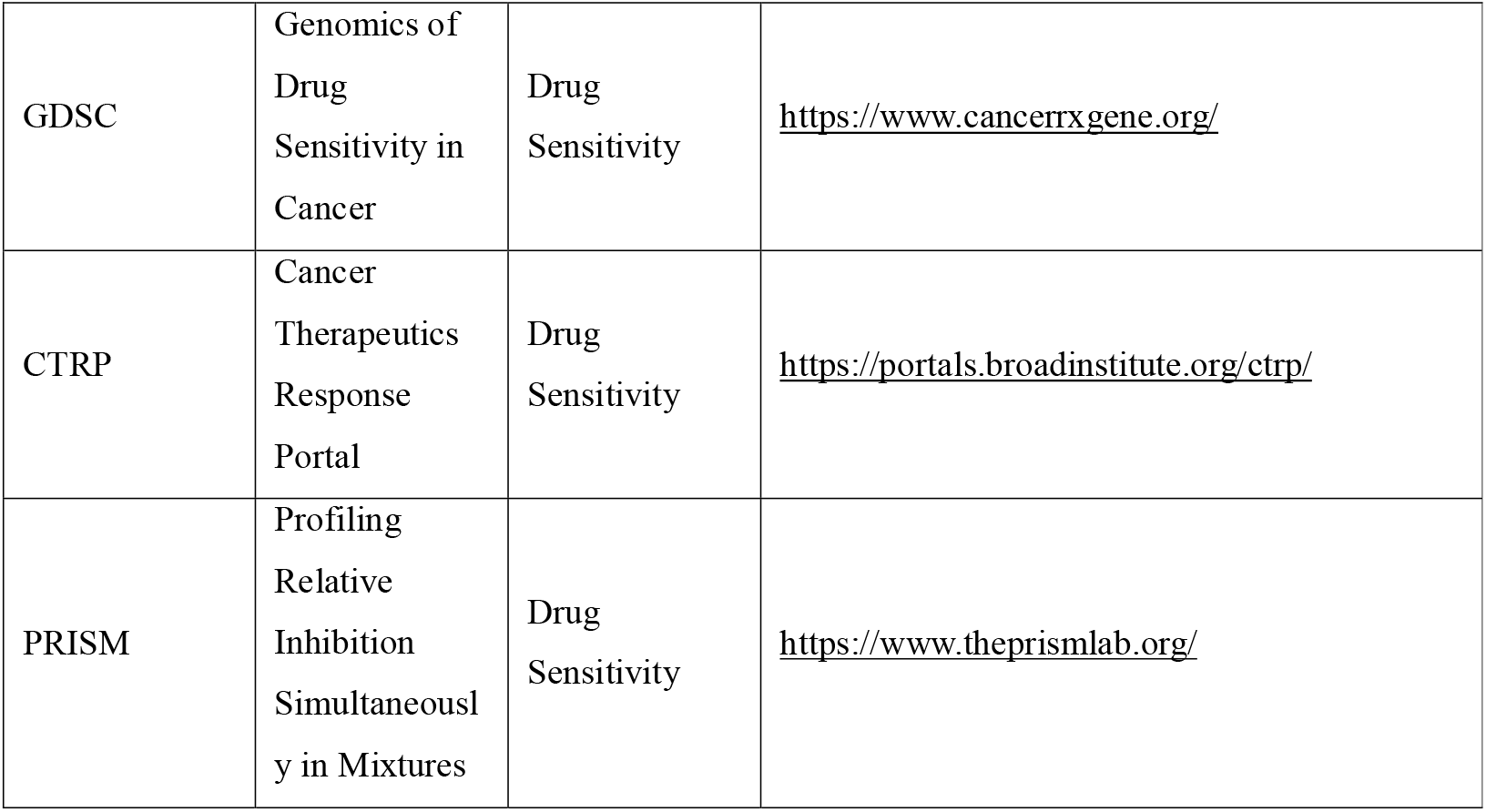

